# The ventral visual stream for reading converges on the transmodal language network

**DOI:** 10.1101/2025.09.17.676937

**Authors:** J.J. Salvo, M. Lakshman, A.M. Holubecki, Z.M. Saygin, M.M. Mesulam, R.M. Braga

**Affiliations:** Ken and Ruth Davee Department of Neurology, Northwestern University Feinberg School of Medicine; Department of Psychology, The Ohio State University; Mesulam Institute for Cognitive Neurology and Alzheimer’s Disease, Northwestern University Feinberg School of Medicine; Department of Psychology, Northwestern University

## Abstract

Reading bridges sensation and cognition. To derive meaning from written words, visual input is first processed in unimodal (i.e., sensory-specific) visual streams and then engages a distributed language network (LANG) that includes classic perisylvian language areas and supports transmodal (i.e., sensory-nonspecific) functions. A reading-relevant region in the inferotemporal cortex (ITC), sometimes called the visual word form area (VWFA), has been the subject of controversy because it displays properties of both systems: it responds to meaningless written pseudowords, implying a unimodal visual function, but also responds to meaningful speech, implying a transmodal function. We investigated whether precision functional mapping could help clarify this region’s role in reading. We characterized a stream of visual regions along the ITC that responded preferentially to visual orthographic forms (i.e., written pseudowords, consonant strings, and real words). Network mapping revealed that only the most anterior region of this “orthographic stream” was connected to the LANG network and accordingly showed responses to meaningful speech. Furthermore, this anterior region was more selective, responding preferentially to text-based stimuli, whereas the more posterior regions of the stream were additionally activated by perceptually similar images (i.e., number strings, foreign script). Our results support that connections to the LANG network may drive specialization along the orthographic stream for writing. This basal language network region may represent an interface between visual and transmodal language systems, thus serving as a critical nexus for reading.

## Introduction

Language comprehension is a “transmodal” function because the same meaning can be communicated through both visual and auditory sensory modalities (Mesulam, 1998). In reading, comprehension is derived from visual input that is initially processed through “unimodal” brain systems that are primarily dedicated to visual information processing. Understanding this interplay between transmodal and unimodal processes is therefore essential for clarifying the neuroanatomy of language and reading, and more broadly of how high-level cognition emerges from sensory processing.

Brain areas serving unimodal vs. transmodal functions follow distinct organizational patterns (Goldman-Rakic, 1988; Mesulam, 1998). Unimodal areas are organized as hierarchies through which sensory input is processed in stages, with information from earlier (upstream) stages integrated into increasingly abstract representations by subsequent (downstream) stages (Felleman & Van Essen, 1991; Mesulam, 1998). In the ventral visual stream, along the posterior half of the inferotemporal (ITC), earlier regions in the hierarchy are sensitive to stimuli in specific visual field locations (i.e., are retinotopic) as well as to basic visual attributes such as lines, shapes, and color (Desimone et al., 1985; Hégde & Van Essen, 2000; Grill-Spector & Malach, 2004). These posterior ITC regions transmit information downstream to more anterior ITC regions. The laminar pattern of neural projections between hierarchical stages implies that posterior-anterior projections are feedforward, while anterior-posterior projections are feedback (Felleman & Van Essen, 1991). This directionality is supported by response delays to presented visual stimuli (Schmolesky et al., 1998; Woolnough et al., 2021) and the finding that the later stages are sensitive to more abstracted features, such as categories of images including faces, scenes, or objects (Kanwisher, 2010).

In contrast to the unimodal hierarchies, networks in transmodal association cortex tend to be activated by cognitive processes that are not restricted to a single sensory or motor modality (Mesulam, 1990; 1998). These networks comprise regions distributed across multiple zones of association cortex which are linked by longer-range, reciprocal connections, rather than the feedback and feedforward projections of sensory hierarchies (Mesulam, 1990; Goldman-Rakic, 1988). This architecture may support emergent, integrative functions that are determined by interactions across the entire network and allow for widespread distribution of information throughout the brain.

A classic example of a transmodal distributed network is the language network (LANG; Geschwind, 1970; Mesulam, 1990; Monti et al., 2009; Price, 2010; Pallier et al., 2011; Lipkin et al., 2022), which canonically includes regions in perisylvian frontal and temporal zones. Recent findings suggest this network is much more widely distributed and includes regions such as ‘area 55b’ in posterior middle frontal cortex (Glasser et al., 2016), as well as in the supplementary motor area and possibly other anterior and posterior midline regions (Braga et al., 2020). The resulting distribution of regions matches the organization of other association networks, suggesting that the LANG network supports a transmodal function (Braga et al., 2020). Accordingly, the LANG network responds strongly to comprehensible language input, regardless of sensory input modality (Fedorenko et al., 2010; Scott et al., 2017; Salvo & Anderson et al., 2025). One framework describes this network as serving “core” transmodal aspects of language processing, while also communicating with an extended language network that includes unimodal sensory and motor regions necessary for language reception and production (Fedorenko et al., 2024). This framework implies that, during reading, the distributed LANG network interacts with and derives input from the visual streams located in the ITC. However, it is unclear what features of functional organization could support these interactions.

Of particular importance to reading, one of the downstream regions in the visual hierarchy shows preferential responses to strings (i.e., combinations) of recognizable letters, regardless of whether the strings form established words (Cohen et al., 2002; Dehaene et al., 2005; Dejerine, 1892; Woolnough et al., 2021). This region, often referred to as the visual word form area (VWFA), is thought to develop as one acquires reading ability (Dehaene et al., 2010; Saygin et al., 2016) and may be central to literacy. The VWFA has attributes of a unimodal visual region because it is situated within the posterior half of the ITC and shows responses to meaningless character strings (e.g., pseudowords; Baker et al., 2007; Cohen et al., 2002). However, the VWFA also displays transmodal properties, including activation for both visual and auditory linguistic stimuli (Forseth et al., 2018; Ludersdorfer et al., 2013; Yoncheva et al., 2010) as well as sensitivity to learned characteristics such as stimulus familiarity (e.g., native vs. foreign words, frequent vs. infrequent letter combinations; Nobre et al., 1994; Baker et al., 2007; Szwed et al., 2011; but see Vogel et al., 2012b). These properties suggest the VWFA may display properties of both unimodal visual and transmodal language systems, which challenges the notion of a strict network-to-function correspondence.

Interestingly, there is evidence that the LANG network includes a region where the VWFA is typically reported, approximately halfway down the posterior-anterior axis of the ITC (Lüders et al., 1986; Nobre et al. 1994). It is unclear whether this basal temporal language region is synonymous with the VWFA and can explain responses to speech within the vicinity of the VWFA, or whether VWFA and basal-language responses in the ITC are supported by different network connections. Attempts to map the network connectivity of the VWFA have led to divergent results, possibly because portions of the ITC suffer from functional magnetic resonance imaging (fMRI) signal dropout (Embleton et al., 2010; Olman et al., 2009). Some studies have provided evidence that the VWFA is connected to the language network (Stevens et al., 2017; Li et al., 2020). Others have argued that the VWFA is connected to the dorsal attention network, which serves visuospatial attention (Stevens et al., 2017; Vogel et al., 2014; Chen et al., 2019).

Recently, we demonstrated that using intrinsic functional connectivity (iFC) to define brain networks within individual participants can distinguish between adjacent transmodal language and unimodal auditory regions in lateral temporal cortex, despite these regions showing overlapping response profiles (Salvo & Anderson et al., 2025). Specifically, we used iFC to delineate LANG and showed that the network demonstrated robust responses to meaningful word combinations presented either visually (as written text) or auditorily (as spoken language; Salvo & Anderson et al., 2025; Scott et al., 2017). Here, to investigate the functional anatomy of reading, we asked whether individualized network definition based on iFC could similarly distinguish between transmodal language and unimodal visual regions in the ITC. We applied precision functional mapping to contextualize visual and language-related responses to multiple stimuli within each individual’s large-scale brain network organization. To limit signal dropout in anterior ITC, we used multi-echo fMRI. Results were replicated in an existing 7T fMRI dataset (Allen et al., 2022). We observed a conserved pattern across individuals, in which a stream of unimodal visual orthographic regions (responding to written words and reading-related stimuli) overlapped with the transmodal language network at a site within the mid-ITC (**Fig. 1**). We suggest that this region may serve as a reading nexus between visual and linguistic processing.

**Fig. 1:**
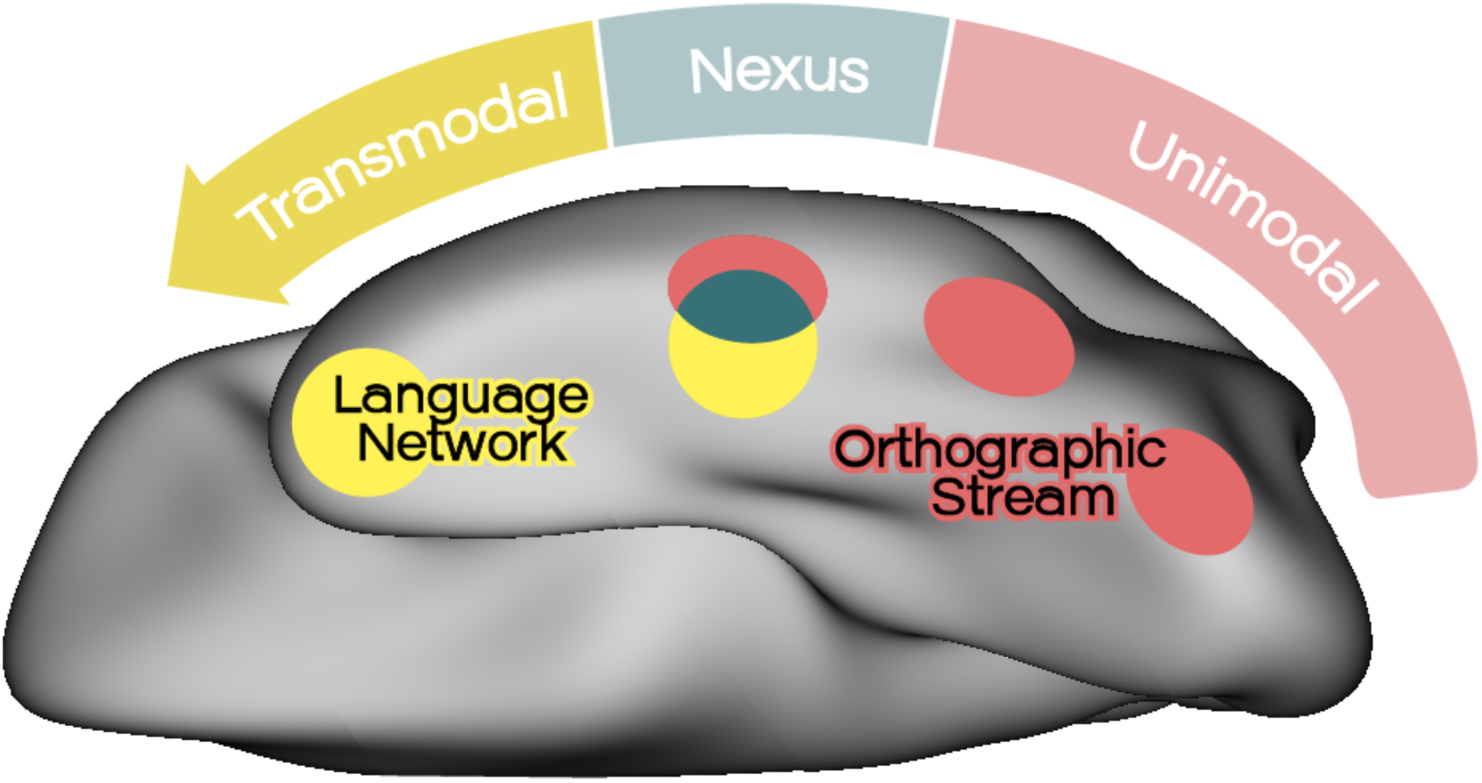
Overview of the reading nexus model. The basal language region (dark teal) represents a nexus between unimodal processing in the orthographic visual stream (red) and transmodal processing in the distributed language network (yellow). Written words are initially processed by the orthographic stream, which preferentially responds when viewing letter strings, even when these do not represent meaningful words. However, the anterior stream region is functionally different: it is connected to the LANG network as defined by iFC and responds when listening to speech. Further, the anterior stream region is more selective for reading-related visual input, selectively activating for familiar (i.e., learned) scripts, whereas the posterior stream regions don’t discriminate between familiar and unfamiliar (i.e., foreign) scripts, even though both types of writing share similar visual features. Thus, the nexus could represent an interface between visual and language systems, where visual processing engages the distributed LANG network for processing viewed meaningful written words linguistically.

## Methods

### Detailed Brain Network Organization (DBNO) Dataset

#### Subjects and sessions

Ten adults (6 female, ages 22–36, mean age 26.6, 9 right-handed) from the local community were recruited as part of the ‘Detailed Brain Network Organization’ (DBNO) study. Details regarding this dataset have been previously described in Kwon et al., 2025, Edmonds et al., 2024, and Salvo & Anderson et al., 2025. Participants had normal hearing and normal or corrected to normal vision, and no history of neurological illness. Informed consent was obtained as approved by the Northwestern University Institutional Review Board. Participants were compensated for participation.

Participants were invited to 8 magnetic resonance imaging (MRI) sessions, each featuring several cognitive tasks as well as two runs of a passive fixation (REST) task for network mapping using iFC. Prior to the first MRI session, participants were trained on all the tasks and were extensively coached about strategies for staying still in the scanner to improve data quality. While in the scanner, participants’ heads were padded using inflatable cushions to restrict head motion. Participants were also informed that the initial 1-2 MRI sessions would serve as a trial to assess compliance and whether they wanted to continue participation. Based on these criteria, 2 subjects were not invited back for additional sessions, leaving 8 subjects who completed all 8 MRI sessions (4 female, ages 22-36, mean age 26.75, 7 right-handed). This led to a total of 60.8 hours of fMRI data collected (∼7.6 hours per participant) for precision functional mapping.

#### Passive fixation task (REST)

To allow estimation of large-scale networks within each individual using iFC, participants completed a passive fixation task. Participants were shown a crosshair in the center of the screen and were instructed to fixate their gaze on the crosshair for the duration of the task, keeping their eyes open and blinking normally. The task lasted ∼7 minutes. Each participant completed 2 runs in each session (the first and final runs of the session) for a total of 16 runs and ∼112 minutes per participant. One participant (S2) completed an additional REST run to replace a poor-quality run that had been excluded in a prior session (see MRI quality control).

#### Cognitive tasks

Participants performed tasks targeting specific cognitive domains. Multiple runs of each task were collected to allow within-individual estimation of functional regions. The tasks targeted brain regions associated with language processing (SPEECHLOC) and recognition of visual categories (VISCAT, VWFACAT).

#### Speech-based language localizer (SPEECHLOC)

To identify regions responding to spoken language input, the SPEECHLOC task used auditory stimuli (speech) and lasted ∼7 minutes per run (based on Scott et al., 2017; for a full description, see Salvo & Anderson et al., 2025). In each trial, participants listened to audio clips of normal, intelligible speech and distorted, unintelligible speech. Audio clips were taken from recordings of TED talks, The Moth Radio Hour, and Librivox audiobooks. Two clips were taken from each recording, with one clip left intact and the second distorted by filtering as outlined below. This meant that the distorted clips preserved certain sound characteristics (speaker identity, intonation, speed, recording quality, etc.) to improve their effectiveness as control stimuli. Maps from this task were previously reported in these participants in Salvo & Anderson et al. (2025).

Audio was presented through Sensimetrics S14 MRI-compatible earphones. To ensure that the clips were audible over the scanner noise, at the start of each MRI session, prior to data collection, we tested sound volume while running a dummy version of the MR imaging sequence to mimic the conditions of the task. Participants were played two speech clips (one clear, one distorted) followed by the tone used as a button-press cue. Participants used the provided button box to adjust the volume to a comfortable level at which the speech could be clearly heard. They were then asked to describe the topic of the clearly presented story.

#### Visual categories task (VISCAT)

The VISCAT task targeted regions of the visual hierarchy responding preferentially to certain image categories. Participants were required to attend to a screen while images were presented and report if an image constituted a repeat of the previous image (i.e., a 1-back task). Images were cropped into a square of 256 x 256 pixels and set to grayscale. Images belonged to five categories: faces, scenes, pseudowords, objects and scrambled stimuli. The scrambled stimuli were created using images from the face, scene, and object categories, which were split into a grid of 13 x 13 squares and randomly shuffled in the 2 dimensions of the grid. Face, scene, object, and scrambled images were provided by Talia Konkle (Park et al., 2015; Konkle et al., 2011). Face images depicted individuals, subjectively appearing roughly between the ages of 10–40, over a white background, with 70 of the 120 individuals appearing male. Hair was included for all images, and upper shoulders were included for 80 images. Scene images depicted both urban and natural settings, with 41 of the 120 images photographed indoors. Object images depicted objects ranging from a thimble to a hot air balloon (all objects were displayed at the same size) over a white background. Pseudoword stimuli were generated by author J.J.S. using the Australian Research Council (ARC) Nonword Database (Rastle et al., 2002) and were checked by two lab members. Any pseudowords resembling real words were replaced.

Images were presented by category in 18s blocks, with 9 images presented in each block, including 1-2 repeats. Each image was displayed for 0.7s, followed by an interstimulus interval with a ‘+’ presented at the center of the screen for 1.3s. Blocks were counterbalanced within each run, and across runs. The fifth blocks in each run were followed by an 18s fixation period where a ‘+’ was presented in the center of the screen. Participants completed 1 VISCAT run per session,which lasted 4 minutes (240 s) and featured 75 unique images (plus 15 repeats). Across 8 runs, participants saw 720 images (600 unique, 120 repeats), about 32 minutes of data.

#### Visual word form area categories task (VWFACAT)

The VWFACAT task followed the same structure as VISCAT, targeting the VWFA through several categories of stimuli that resembled familiar written text to varying degrees (real words > pseudowords > consonant strings > foreign script > line drawings). Line drawings were used as a control, since they were stylistically similar to the written characters (a complex series of black lines on a white background) without appearing to be a script.

Real words were sourced from a list of the most frequently used 5050 English words in the Corpus of Contemporary American English (COCA) 2020 edition (Davies, 2020). Curses, proprietary (e.g. ‘iPad’), and potentially distressing (e.g. ‘tumor’) words were removed from the list. Word length was limited to 4-8 letters long, which accounted for 75% of the corpus. We next determined what percentage of words in this range fell into each word length category (e.g. 4-letter words accounted for 23% of all 4-8 letter words), and randomly selected words from the list while ensuring that the selected set maintained the word-length percentages of the broader corpus. We also ensured that selected words did not repeat the same word families (e.g. ‘sail’ and ‘sailed’).

A new set of pseudowords were generated using the ARC Nonword Database, distinct from those used in the VISCAT task. To create consonant strings, we first selected a new list of words from the 2020 COCA. We next replaced vowels in each word with randomly chosen consonants and shuffled the characters. Consonant strings were each checked, and those that still resembled real words were removed. For the foreign script condition, we employed hiragana and katakana, two Japanese scripts that are illegible for participants illiterate in Japanese. Characters resembling Roman characters were modified to appear less familiar to English readers (e.g. the hiragana character ‘て’, resembling a Roman ‘T’, was mirrored and rotated) by editing a Japanese font (KaoriGel) using the free program Glyphr. To reprint the consonant strings using Japanese characters, we created a new font (KaoriGelModified) with each selected Japanese character corresponding to a Roman character keyboard key. Prior to inclusion in our study, participants were screened to verify they were not familiar with the foreign script (i.e., they could not read in Japanese or other East Asian scripts).

Line drawings were taken from a stimulus set provided by Nishimoto and colleagues (2012) that was adapted from a set from Snodgrass & Vanderwart (1980). The latter study drew images from standardized categories (Battig & Montague, 1969), including animals, vehicles, and human body parts. Images thus depicted animate or inanimate subjects (e.g. a bird vs. a dress) and included objects with a range of actual sizes (e.g. a ladybug vs. a tractor). Each included image was shaped into a 256 x 256 pixel grayscale square to match VISCAT stimuli.

Participants performed a 1-back task using the VWFA stimuli, using the same block design as the VISCAT task. Participants completed 1 VWFACAT run per session, which lasted about 4 minutes (240 s) and featured 75 unique images (plus 15 repeats). Across 8 runs, participants saw a total of 720 images (600 unique, 120 repeats), about 32 minutes of data.

#### MRI data acquisition

Data were collected on a Siemens 3.0 T Prisma scanner (Siemens, Erlangen, Germany) at the Center for Translational Imaging at Northwestern University’s Feinberg School of Medicine in Chicago. Two anatomical images were collected: a T1-weighted image in the first MRI session (TR = 2,100 ms, TE = 2.9 ms, FOV = 256 mm, flip angle = 8°, slice thickness = 1 mm, 176 sagittal slices parallel to the AC-PC line) and a T2-weighted scan in the second session (TR = 3,000 ms, TE = 565 ms, FOV = 224 mm x 256 mm, flip angle = 120°, slice thickness = 1 mm). Both scans included volumetric navigators (Tisdall et al., 2012) from the Adolescent Brain Cognitive Development study (Hagler et al., 2019). Functional MRI data were collected using a 64-channel head coil using a multi-band, multi-echo sequence with the following parameters: TR = 1,355 ms, TE = 12.80 ms, 32.39 ms, 51.98 ms, 71.57 ms, 91.16 ms, flip angle = 64°, voxel size = 2.4mm, FOV = 216 mm x 216 mm, slice thickness = 2.4 mm, multiband slice acceleration factor = 6 (see Poser et al., 2006; Lynch et al., 2020).

#### Quality control

Head motion was estimated using FSL’s MCFLIRT command (Jenkinson et al., 2002). Across all task types, runs were automatically excluded from analysis if motion exceeded predetermined thresholds of framewise displacement (FD) > 0.4mm or absolute displacement (AD) > 2.0 mm. Runs were flagged for visual inspection if motion exceeded more stringent thresholds (FD > 0.2 mm, AD > 1.0 mm). The whole run was excluded if motion could be clearly seen in the raw data. Runs for the remaining three localizer tasks were also excluded if behavioral performance was poor, with thresholds specific to each task (as follows).

For REST, each participant retained at least 10 runs following quality control (range: 10-16). For all participants except one (S5), half of the data were set aside for replication analyses (S1: 56 min, S2: 56, S3: 56, S4: 56, S5: 60, S6: 49, S7: 49, S8: 49), leaving 49–60 minutes of data for each participant.

For SPEECHLOC, runs with more than 20% missing button-presses were excluded. At least 7 runs were retained per participant (S1: 8, S2: 8, S3: 8, S4: 7, S5: 7, S6: 8, S7: 7, S8: 8), leaving 49–56 minutes of data per participant.

For VISCAT, runs with more than 10% missing button-presses and/or incorrect answers were excluded. All 8 runs were retained for all participants, leaving 32 minutes of data per participant.

For VWFACAT, runs with more than 10% missing button-presses and/or incorrect answers were excluded. At least 7 runs were retained per participant (S1: 8, S2: 8, S3: 7, S4: 8, S5: 8, S6: 8, S7: 8, S8: 8), leaving 28–32 minutes of data per participant.

#### MRI data preprocessing

Blood-oxygenation-level-dependent (BOLD) data were preprocessed using a custom processing pipeline “iProc” (Braga et al., 2019) optimized for within-individual alignment of data from multiple runs and sessions. Additional steps were included to account for the multi-echo data (as outlined below from Salvo & Anderson et al., 2025 and reported in Edmonds et al., 2024). Registration steps were combined into a single interpolation to minimize blurring. Each individual’s data were processed separately. The first 9 volumes (∼12 s) of each run were discarded to remove a T1 attenuation artifact. Next, a mean BOLD template was created as an interim stage for data registration. This was done by averaging the first echo of all included runs of all tasks, to reduce bias toward one run. The participant’s T1 image was used to create a native space template. These templates were used to build four matrix transforms to align each BOLD volume: (1) to the middle volume of the same run for motion correction; (2) to the mean BOLD template for cross-run alignment; (3) to the native space template; and (4) to the 152 1-mm atlas (Mazziotta et al., 1995) from the Montreal Neurological Institute (MNI) International Consortium for Brain Mapping. The four transforms were combined and applied to the original volumes to register all volumes to the T1 anatomical template (matrices 1–3) and to the MNI atlas (matrices 1–4) in a single step. The MNI registration was used in the nuisance regression steps applied to the resting-state runs (see *Functional connectivity preprocessing* below). Visual checks were incorporated into each registration step of the pipeline.

Motion correction transforms were calculated using rigid-body transformation based on the first echo, which had less signal dropout and better preserved the shape of the brain. Registration matrices were calculated based on the first echo and applied to all echoes. Once data were projected to the T1 native space, the five echoes were combined to approximate local T2* (Heunis et al., 2021). Briefly, the tSNR was calculated for each echo, which was then weighted by the echo time. This weighted tSNR was then divided by the sum of all weighted tSNRs (i.e., for all echoes). The resulting image was multiplied by the echo’s original intensity image, and these were summed across all echoes to create the final optimally combined image.

#### Functional connectivity preprocessing

Nuisance variables were calculated for each run by extracting the average signal from deep white matter and from cerebrospinal fluid masks hand-drawn in MNI space that were back projected to native space. Additional nuisance variables included six motion parameters, whole-brain (global) signal, and their respective temporal derivatives. Data were then bandpass filtered using a range of 0.01–0.1 Hz using *3dBandpass* (AFNI v2016.09.04.1341; Cox, 1996; 2012). Next, data were projected to a standard cortical surface (fsaverage6, 40,962 vertices per hemisphere; Fischl et al., 1999) and smoothed with a 2 mm full-width at half-maximum (FWHM) kernel along the surface. This kernel size was chosen based on prior work (Braga & Buckner, 2017; Braga et al., 2019) to retain functional anatomic detail while limiting noise (i.e., speckling) in the functional connectivity maps.

#### Assessing signal dropout

To assess data quality and the interpretable zones of the inferior temporal cortex (i.e., regions that did not show evidence of signal dropout), we calculated average tSNR maps for each participant using all runs of the preprocessed, optimally combined data. Values were projected to the FreeSurfer surface mesh fsaverage6. As a visual aid for outlining the interpretable zones of ITC, author J.J.S. manually drew borders around tissue exhibiting the lowest tSNR values (about 20 or less) using the pen tool in Adobe Illustrator.

The tSNR maps revealed that in each participant, good tSNR was achieved throughout the cortex using the multi-echo BOLD sequence. Regions showing signal dropout were minimal and primarily within a restricted portion of anterior ITC (**Supp. Fig. S1**, see dashed lines), helping build confidence in the functional anatomy revealed in the middle and posterior half of the ITC. Nonetheless, tSNR was generally lower on the inferior temporal surface than on the lateral surface, possibly due to the distance of this surface from the skull and the MR coils.

#### Network estimation

Prior to any analysis of the task data, the independent REST data were used to create individual-specific estimates of large-scale networks using iFC following previously used procedures (Braga & Buckner, 2017; Braga et al., 2020). We only applied this analysis to half of the REST data, setting aside the remaining half for *Replication of the candidate language network from inferior temporal seeds* (see below).

A brief description of this process, previously reported in Salvo & Anderson et al. (2025), follows. First, functional connectivity matrices were calculated for each REST run by computing vertex-vertex Pearson’s product-moment correlations. The matrices were Z-normalized using the Fisher transform, averaged across all runs within each individual, and converted back to r values using the inverse Fisher transform. These within-individual average matrices were used to perform a seed-based analysis.

Seeds were manually selected to delineate eight networks, including LANG but also others as presented in Edmonds et al. (2024) and Kwon et al. (2025). Here, we focus on LANG and show the resulting network in **Fig. 2** but note that the initial definition of networks defined the LANG network based on its separation from surrounding large-scale networks defined using other seeds. We used an interactive platform for manual seed selection and correlation map viewing based on Connectome Workbench (Marcus et al., 2011). Seeds were chosen to target LANG in the left hemisphere in the posterior superior temporal cortex (pSTC), posterior middle frontal gyrus, inferior frontal gyrus, temporal pole, and supplementary motor area. These provided converging estimates of the LANG network (**Supp. Figs. S2–S9**). Seeds in the pSTC yielded comparable correlation patterns whether chosen generally in lateral temporal cortex vs. specifically in posterior lateral temporal cortex, closer to the typical site of Wernicke’s area (Tremblay & Dick, 2016). The identification of LANG was based on anatomical distinctions characteristic of the network (described in Braga et al., 2020). Comparisons to task activity during a speech listening (**Fig. 3**) and reading task (Salvo & Anderson et al. 2025) confirmed this network was recruited for linguistic processes.

To ensure that the estimated seed-based networks were not determined by observer bias, networks were also defined using a data-driven *k*-means clustering algorithm (**Fig. 2**). This process was carried out using two approaches. First, we used a Von Mises-Fisher distribution, based on Yeo et al. (2011). Second, we used a multi-session hierarchical Bayesian Model (MS-HBM), a method that stabilizes individual-specific network estimates by integrating priors from multiple levels (e.g., group-level, cross-individual and cross-run variation; Kong et al., 2019). For each individual, we generated clustering solutions with *k* values (i.e., number of clusters) between 7–20 and selected the lowest level of clustering that separated the networks of interest in a manner defined by the seed-based approach. For the Von Mises-Fisher maps in two participants (S2, S4), no *k*-value included all networks without over-splitting at least one network (i.e., the resulting networks did not match the distribution of regions defined using the *a priori* seed-based approach). In these two participants, two clusters were merged to yield the candidate language network. As shown in **Supp. Figs. S10–S11**, the combined networks better matched the seed-based correlation and task activation pattern in these subjects, including in the ITC.

We compared the network estimates from the Von Mises-Fisher and MS-HBM clustering approaches by eye, focusing on how well they captured the ITC region of LANG defined from the multiple seed-based estimates (**Supp. Figs S2–S9**). Results were largely comparable, but did show variation in some subjects, as shown in **Supp. Fig. S12**. Ultimately, we selected the Von Mises-Fisher approach as it included a prominent ITC LANG region in more subjects. We believe these differences reflect quirks of the clustering algorithms (e.g., the MS-HBM uses group-level priors that may be sub-optimal for delineating functional regions in lower signal quality zones).

#### Task activity maps

Data from the in-scanner tasks were analyzed using FSL’s FEAT (Woolrich et al., 2001). The surface-projected task fMRI data for left and right hemispheres was input into a separate general linear model. The task was modeled using explanatory variables that were convolved with a double-gamma hemodynamic response function then used as regressors using FEAT to define maps of parameter estimates as outlined below (also see *Cognitive tasks*). For each task, values from the contrast-of-parameter-estimates maps from each run were then normalized into z values by FEAT and averaged across all runs for a given task and individual.

For SPEECHLOC (**Fig. 3**), regressors included all presented audio clips for each of the two speech types (comprehensible and distorted). Beta values were calculated for the comprehensible and distorted speech conditions, and a contrast of parameter estimates was calculated between the conditions for each run.

For VISCAT (**Fig. 4**), regressors included all blocks of images for each of the five image categories (faces, scenes, pseudowords, objects, and scrambled images). Beta values were calculated for each condition, and for each run, a contrast of parameter estimates was calculated between each category and the remaining categories (e.g. pseudowords vs. faces, scenes, objects, and scrambled). We were concerned that the scrambled images might be more difficult to recognize compared to images in other categories, thereby affecting the contrast maps. Ultimately, we opted to omit the scrambled condition from these contrasts.

For VWFACAT, regressors included all blocks of images for each of the five image categories (real words, pseudowords, consonant strings, foreign script, and line drawings). Beta values were calculated for each condition, and a contrast of parameter estimates was calculated between each category and the control condition of line drawings (e.g. pseudowords vs. line drawings) for each run.

#### Thresholding

Task activity was initially visualized using qualitative thresholds, as reported in Salvo & Anderson et al. (2025), chosen by eye for each participant to emphasize larger contiguous regions containing higher values while removing smaller regions and speckling containing lower values (indicating noise). Based on this, thresholds were picked individually for each subject and task contrast map, prior to any assessment of overlap. We started at a threshold of z > 2.5 and lowered the threshold if the activation map appeared smaller than expected (e.g., comparing to prior work; e.g., Braga et al., 2020). To ensure that the results were not specific to these hand-picked thresholds, we created maps thresholded at the 90th percentile for each participant (**Supp. Figs. S13 & S14**), which produced very similar maps. Given their similarity, we used the qualitatively chosen thresholds for all further analyses.

#### Assessment of overlap

To assess overlap between the SPEECHLOC and VISCAT task contrast maps (**Figs. 5 & 6**), we binarized the thresholded maps and plotted them on the same cortical surface. We quantified the degree of overlap in mid-ITC using the Dice coefficient. The mid-ITC zone was defined by drawing a large mask by hand that surrounded the LANG network region in all subjects (shown in **Fig. 5**, lower panel). Only vertices within this mid-ITC mask were considered when calculating the Dice coefficient.

#### Replication of the candidate language network from inferior temporal seeds

To confirm whether mid-ITC regions active for both pseudowords and speech were functionally connected to the language network, we manually chose seeds in regions within the ITC that were active for both conditions (listening to speech and viewing pseudowords). We then applied these seeds to the independent held-out half of the resting-state runs to see if the resulting correlation pattern matched the language network.

### Replication using the Natural Scenes Dataset

#### NSD Overview

To investigate whether we could replicate the spatial relationships between transmodal language and unimodal visual regions in the ITC, we analyzed data from the Natural Scenes Dataset (NSD). The NSD includes functional MRI data collected at a greater field strength (7 T) and voxel precision (1.8 and 1.0 mm) than the DBNO dataset (3 T and 2.4 mm iso, respectively). Detailed descriptions of the participants, experimental paradigms, data quality, and initial data processing can be found in Allen et al., 2022. Additional in-house preprocessing for functional connectivity analysis was performed as described in Kwon et al., 2025, and Edmonds et al., 2024.

#### NSD participants and sessions

The NSD includes eight healthy human participants (two males and six females; 19–32 years old) recruited from the University of Minnesota community. Participants displayed normal or corrected-to-normal vision and had no known cognitive deficits. All participants were right-handed and fluent in English. Three participants were multilingual, with one having learned English as a second language. Participants completed an initial screening session and 30–40 MRI sessions that included two resting-state fMRI scans alongside a visual image recognition task. The study ended with two task-based scan sessions measuring responses to other task contexts. This resulted in 33–43 MRI sessions per subject along with additional behavioral assessments.

#### NSD resting-state data

Resting-state data were collected across 10 sessions (except for subject NSD-S1, who had 18 sessions), providing an average of 2 hours of resting-state fMRI data per participant, over an average of 77 days. In one form of this task, participants kept their eyes on a white, centered plus sign over a gray background for 300s. In a second form of this task, the white plus-sign turned red after 12s and remained red for 4s before returning to white for the remainder of the run. For runs following the second format, participants were asked to breathe in when the plus-sign turned red and hold their breath until the plus-sign returned to white. Each participant completed 100–180 minutes of resting-state runs.

#### NSD visual category functional localizer task (‘fLoc’)

Task data from the visual category functional localizer task (‘fLoc’) were collected in a single session at the start of the study for all subjects. Participants were presented with sets of stimuli sorted by visual category. All stimuli were presented on top of unique abstract backgrounds in grayscale. A central red dot was present throughout the task. Participants were asked to report with a button press whenever a ‘blank’ image was presented (i.e., a background was presented without a stimulus image). Each stimulus was shown for 0.5s, with each run lasting 312s and containing 48 images per category (480 non-blank images). Six runs were collected per participant for a total of 288 images per category (2,880 non-blank images) across 31.2 minutes. Each stimulus was shown only once.

The NSD fLoc task included images falling within 10 categories in 5 domains. Eight categories followed four ‘non-character’ domains: bodies (body and limb), faces (adult and child), places (corridor and house), and objects (car and instrument). The other 2 categories followed a ‘character’ domain, which included ‘words’ (pseudowords) and ‘numbers’ categories. For our key analyses, we studied the maps derived from contrasting the pseudoword and number categories against all other domains together (‘bodies’, ‘faces’, ‘places’, plus ‘objects’). Pseudoword stimuli were English-pronounceable pseudowords, while number stimuli were uncommon sequences of digits. Pseudoword/numeral length and font type, size, shading, angle, and distortion varied across stimuli. We also mapped activity during viewing of faces and scenes (contrasted against viewing the other stimuli) to replicate results from the DBNO dataset.

#### NSD quality control & preprocessing

As reported in Kwon et al. (2025) and Edmonds et al. (2024), we performed quality control following a two-step process. First, head motion was estimated using FSL’s MCFLIRT command (Smith et al., 2004). Whole runs were automatically excluded if (i) the maximum FD was > 0.4 mm and (ii) the maximum AD was > 2.0 mm. Second, any runs where the FD exceeded 0.2 mm or AD exceeded 1.0 mm were visually inspected for head motion and excluded if any motion could be clearly seen. Following quality control, two of the eight NSD participants were excluded due to excessive head motion.

The remaining NSD participants retained all 6 fLoc runs, and 6–35 resting-state runs per participant (NSD-S1: 35 runs; NSD-S2: 6, NSD-S3: 16, NSD-S4: 12, NSD-S6: 19, and NSD-S7: 18). For the two participants who provided 12–16 resting-state runs, half the data (6 to 8 runs) were assigned to a discovery dataset, and the remaining half were assigned to a validation dataset. For subjects with 18–19 runs, one-third of the data were assigned to the discovery dataset, and the remaining two-thirds of the data were assigned to two validation datasets (6–7 runs each). For subject S1, who provided 35 runs, half of the data (17 runs) were assigned to a discovery dataset, and the remaining half were assigned to two validation datasets (9 runs each). This is because we initially divided NSD-S1’s data into two datasets and began exploring half the data before determining that stable estimates could be achieved with six runs.

For iFC analysis, we performed additional preprocessing on the resting-state data following the steps outlined in the iProc pipeline (Braga et al., 2019, see above). Nuisance variables, including six parameters to account for head motion, as well as whole-brain, ventricular, and deep white matter signals, and their temporal derivatives, were calculated and regressed out of the data. Nuisance regression was performed using 3dTproject [AFNI version 2016.09.04.1341 (Cox et al., 1996)] on the native space-projected BOLD data resampled to a 1-mm isotropic resolution (i.e., the “func1p0mm” version of the NSD data). Data were then bandpass filtered at 0.01–0.1 Hz (using 3dBandpass from AFNI).

The original dataset included tSNR maps for each subject (see Allen et al., 2022). To visualize regions with low signal and reliability, we projected these values to a cortical surface representation (fsaverage, 163,842 vertices per hemisphere; **Supp. Fig. S15**). Notably, while the NSD data included high tSNR despite the smaller voxel size, it showed broader signal dropout in the ITC, particularly in the anterior half. This dropout provides important context for interpreting the successful and failed replications.

#### 7T iFC analysis

Preprocessed data were projected onto a standardized cortical surface containing 163,842 vertices (fsaverage) per hemisphere using FreeSurfer’s vol2surf command (Fischl et al., 1999) and smoothed along the surface using a 2.5-mm FWHM Gaussian kernel (as described in Edmonds et al. 2024). The highest-resolution fsaverage cortical mesh was used to minimize blurring during sampling and preserve fine-scale distinctions between networks. iFC matrices were calculated in each participant by computing vertex-vertex Pearson’s product-moment correlations for each run, *r*-to-*z* transforming, and then averaging across runs within each dataset. As for the DBNO dataset, these matrices were used for surface-based network estimation using an interactive platform for manual seed selection and correlation map viewing based on Connectome Workbench (Marcus et al., 2011).

We identified several cortical networks using the same procedure described above for the DBNO data, selecting candidate vertices for each network based on resulting iFC maps that were distinct from one another (other networks shown in Edmonds et al., 2024 and Kwon et al., 2025). Only the LANG network was assessed in this study. As with the DBNO data, we selected multiple seeds on the cortical surface to target different regions of LANG in each subject (**Supp. Figs. S16–S21**). We also performed data-driven clustering using a Von Mises-Fisher distribution (Yeo et al., 2011), picking a *k* value based on the network topography revealed by the seed-based maps. As before, network replication from mid-ITC seeds was performed using the independent held-out runs.

The multiple iFC maps (**Fig. 6**; **Supp. Figs. S16–S21**) provided converging evidence for a distributed organization of LANG including a mid-ITC region in multiple subjects. Of these subjects, NSD-S1, NSD-S2, and NSD-S4 showed the strongest evidence for this mid-ITC region in both the hand-picked seed maps and the data-driven clustering solutions. We focused replication analyses on these 3 subjects, noting that two other subjects (NSD-S3 and NSD-S6) also showed evidence of a mid-ITC LANG region (**Supp. Figs. S18 & S20**).

#### NSD identification of orthographic stream

To replicate the identification of an orthographic stream in the DBNO dataset (see Results), we used data from the fLoc task to separately identify pseudoword-active and number-active regions in the ITC using task maps provided as part of the NSD (Allen et al., 2022). As with the DBNO data, we thresholded the activity maps using both qualitative (hand-selected) and quantitative (95^th^ percentile) thresholds. Given the comparable results and sometimes superior filtering of activity “speckling” with the qualitative thresholds, we opted to use the qualitative thresholds for further analyses, but show the comparison of the two thresholds, along with the unthresholded maps, in **Supp. Fig. S22**.

To replicate the results showing differentiation of responses along the orthographic stream, we focused on two contrasts: pseudowords > non-character domains (i.e., all other categories except numbers) and numbers > non-character domains (i.e., all other categories except pseudowords). Task data were projected to the same cortical surface mesh, using the same registration procedures, as the resting-state iFC data. To test the stimulus preference of the ITC regions, we binarized the thresholded task activity maps and overlaid the results with the cluster-defined LANG estimate.

#### Recreation of candidate networks

As with the DBNO dataset, we tested whether mid-ITC seeds could replicate the LANG network in held-out data. For each patient, a seed was chosen in a region of the mid-ITC demonstrating activity for pseudowords but not for numbers. The seed was applied to the held-out set of resting-state data for each participant.

## Results

### The distributed language network includes a basal temporal region

In all DBNO participants, we defined a distributed language network (LANG) using resting-state iFC through two converging approaches: seed-based correlation and clustering. Network definition and identification were performed without consulting the inferotemporal surface. Following network definition, in 6 out of 8 participants, we observed that the iFC-defined LANG network included a region approximately halfway down the long axis of the ITC (**Fig. 2**).

Because tSNR, correlation, and task activation values were often lower in the ITC (**Supp. Fig. 1**), we used a lower threshold for the ITC-view of the seed-based maps (see colorbars in **Fig. 2**, left column). Importantly, we observed the putative basal LANG region multiple times in each participant, first using seeds targeting LANG in multiple cortical locations (**Supp. Figs. S2–S9**) and second in the data-driven clustering analysis (**Fig. 2**, right column). Additionally, this basal LANG region was left-lateralized (**Supp. Fig. S23**), matching the asymmetry of the remaining LANG network regions (Braga et al., 2020). These results support that a basal LANG region can be reliably observed when iFC is performed within individuals, using sequences that help address signal dropout in the ITC.

### The basal LANG region is activated by listening to speech

We previously showed that LANGdisplays transmodal responses by showing that the network is activated both during listening to speech (Salvo & Anderson et al., 2025; Scott et al., 2017) and reading comprehensible sentences (Braga et al., 2020; Salvo & Anderson et al., 2025). As predicted by its connectivity with LANG, we observed that the basal LANG region defined by iFC also displayed responses during listening to speech, despite its location in mid-ITC near canonically visual areas. Notably, this task-related activity overlapped precisely with the region showing connectivity to LANG, despite being defined using separate data and analysis approaches (**Fig. 3**). Speech responses were observed in mid-ITC even for the two subjects without a basal LANG region (S6 & S8), though these responses were less robust relative to other subjects (**Fig. 3**). These findings support that the basal LANG region serves a transmodal function (see Li et al., 2020; 2024). Speech responses in the basal LANG region were left-lateralized (i.e., stronger and larger on the left hemisphere) in all participants except S6 (**Supp. Fig. S24**), reinforcing the lateralization of the LANG network observed throughout the cortex (**Supp. Fig. S23**). All subjects also showed evidence of additional, more anterior speech-active regions approaching the temporopolar cortex, despite lower tSNR quality there (**Supp. Fig. S1**). In most participants, these temporopolar speech-active regions also overlapped with regions of the iFC-defined LANG network.

**Fig. 2:**
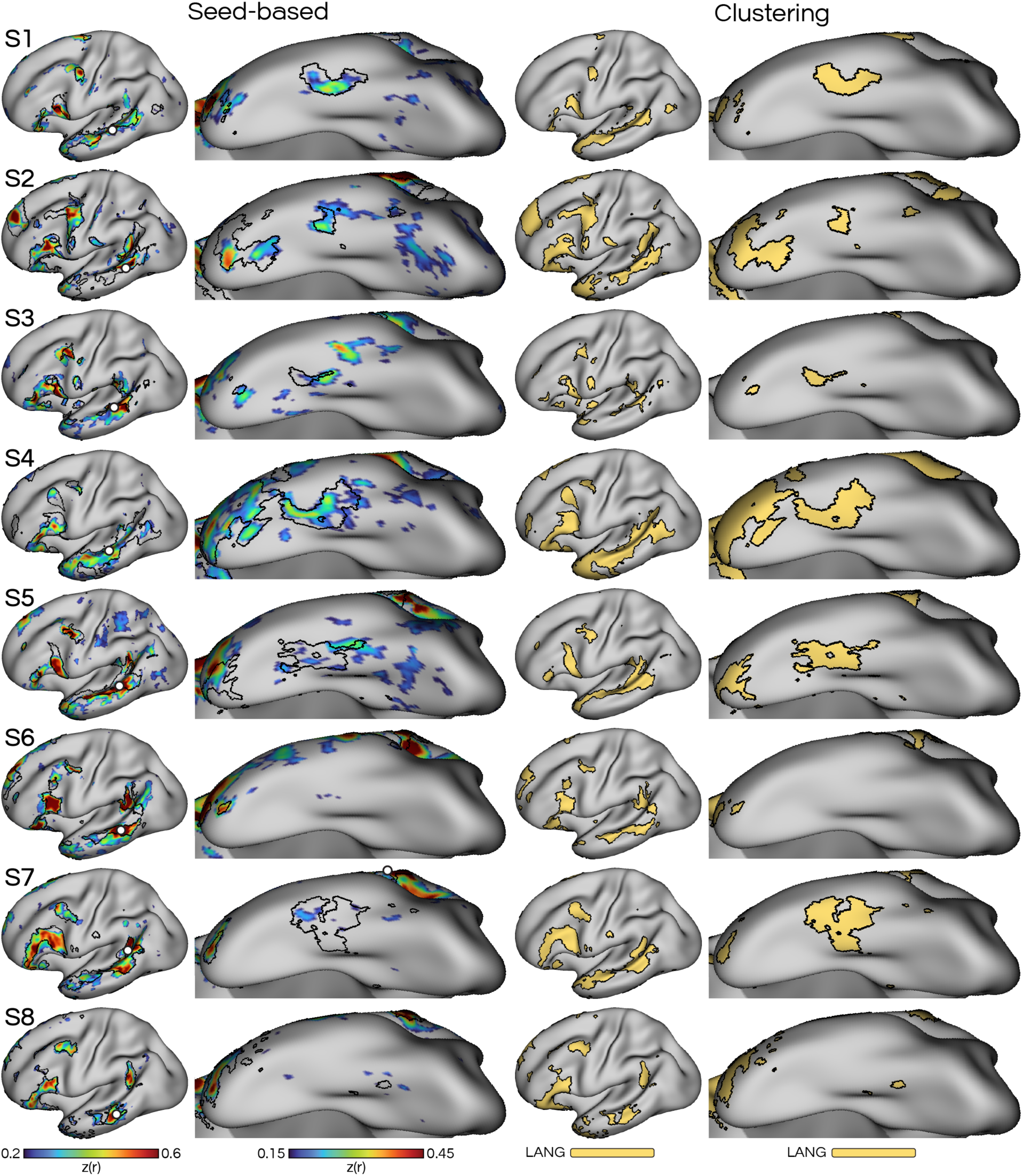
Functional connectivity estimates of the distributed language (LANG) network reveal a basal region in the inferotemporal cortex (ITC) in 6 out of 8 participants. Each row represents a different participant (S1–S8). *Left*: Maps show iFC (Z-scored correlation) maps from seeds (white circles) manually chosen in the lateral temporal cortex, which revealed a correlation pattern that included classic inferior frontal language regions as well as the basal LANG region. Correlations on the lateral surface were thresholded at z(r) > 0.2, with a color range of 0.2–0.6 (Turbo colormap), while for the ITC surface the threshold was lowered to z(r) > 0.15, color range = 0.15–0.45, to account for lower tSNR (see **Supp. Fig. S1**). *Right:* Data-driven clustering confirmed the organization of LANG seen in the seed-based analyses, including the presence of the basal LANG region. For two participants (S2 and S4), the clustering solution divided the seed-based candidate LANG network into two clusters, which were merged (see **Supp. Figs. S10 & S11**).

### Viewing images activates parallel streams along the posterior ITC

Given the LANG network’s role in transmodal linguistic processes, we asked whether the basal LANG region only responded to visual stimuli containing language-relevant information (i.e., writing or ‘orthographic forms’). We mapped category-preferring regions of the ventral visual pathway using our visual category task (‘VISCAT’) to identify regions responding preferentially when subjects viewed pseudowords, faces, or scenes, compared to the other categories (**Fig. 4**).

The image categories activated multiple, interdigitated regions along the ITC. These regions were broadly arranged as three parallel ‘streams’ that extended from the occipital pole towards the temporal pole (**Fig. 4**). The multiple regions in each stream potentially represent different stages of visual processing within each category (Felleman & Van Essen, 1991; Van Essen & Maunsell, 1983). Generally, we observed that the scene-related stream was positioned most ventral, the “orthographic stream” was most lateral, and the face-related stream was in-between. This organization replicated prior work demonstrating the presence of category preferring regions in the ITC (e.g., Kanwisher, 2010; Grill-Spector, 2003; Hasson et al., 2002), and accords with the suggestion that category-preferring visual regions may be arranged as an ‘archipelago’ (Kanwisher et al. 2010).

When we compared the two hemispheres, the orthographic stream was strongly left-lateralized (i.e., appeared larger on the left), while the face and scene streams were more bilateral (**Supp. Figs. S25 & S26**). This feature matched the left-lateralization of both the basal LANG region and the broader LANG network on the lateral surface (**Supp. Fig. 27**), supporting a relationship between lateralization of language and reading (Van der Haegen et al., 2012; Cai et al., 2008). Qualitatively, face-preferring regions may also have been larger on the right (Rossion et al., 2012).

### The orthographic visual stream converges on the transmodal basal LANG region

We observed that the most anterior region of the orthographic stream overlapped consistently with the boundaries of the basal LANG region as defined with iFC (black borders) in each individual (**Fig. 4**). Sometimes, the adjacent face stream region also overlapped with the LANG borders, though this occurred in fewer participants. This selective alignment suggested that the anterior-most orthographic stream region might be embedded in a different network from the more posterior regions, supporting a differentiation of function along the orthographic stream.

**Fig. 3:**
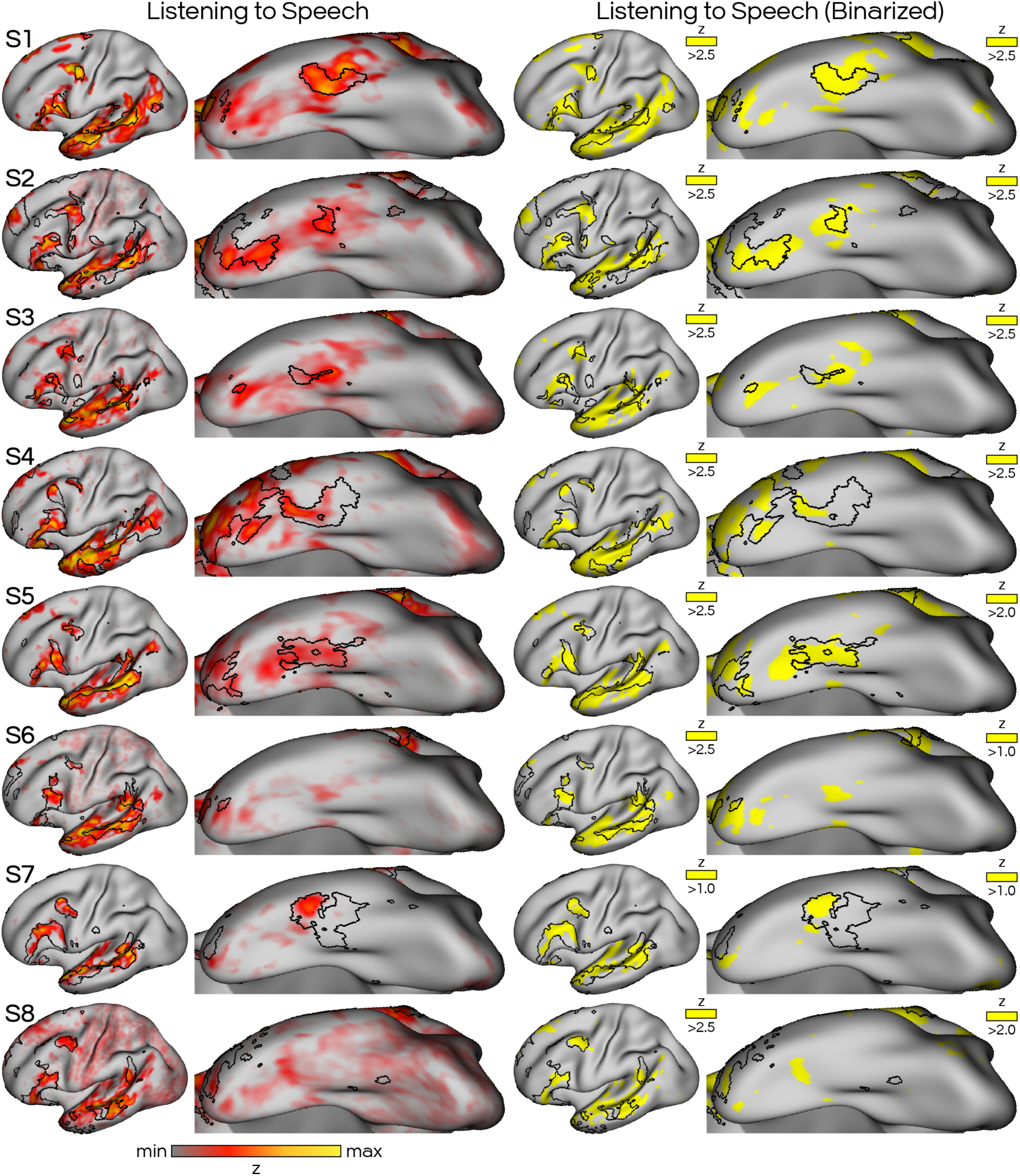
Listening to speech activates the distributed language (LANG) network, including the basal LANG region. *Left:* Heat map indicating Z-normalized beta values for the SPEECHLOC task contrast of listening to speech minus listening to filtered, incomprehensible speech. The iFC-defined LANG network (i.e., clustering map from Fig. 2) for each subject (S1–S8) is shown as black borders. *Right*: The task maps were subsequently thresholded and binarized (yellow) to more easily visualize overlap with visual category streams in later figures.

**Fig. 4:**
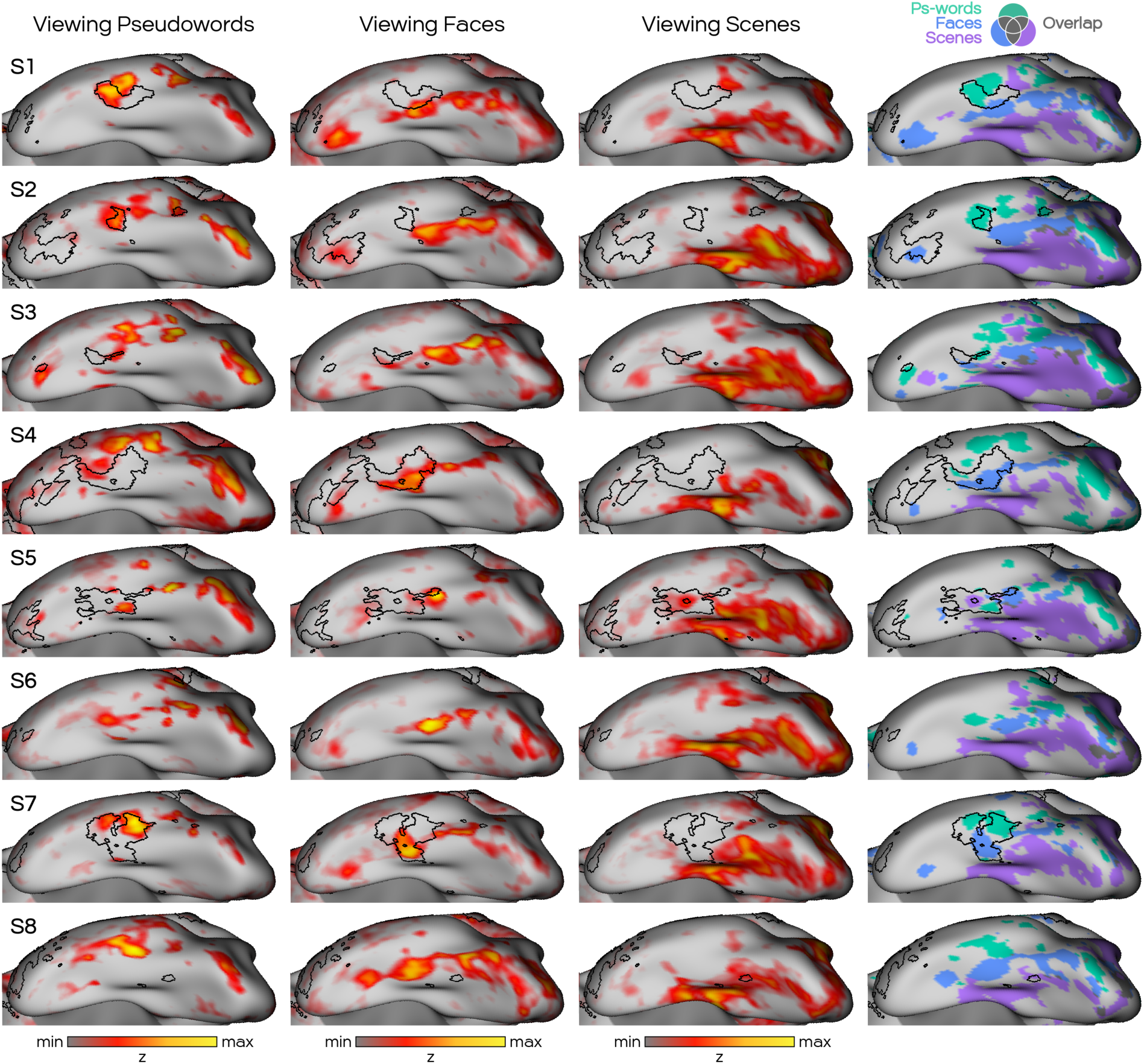
Activity evoked by viewing images of pseudowords, faces, or scenes (VISCAT task) reveals three parallel streams of category-preferring visual regions along the posterior inferior temporal cortex. *Left three columns:* Heat maps show activity evoked by each image category (pseudowords, faces, scenes) contrasted with activity for the other categories plus an ‘objects’ category. Color bars display the full range of positive Z-normalized beta values per participant and are shown with thresholds overlaid in **Supp. Fig. S13**. *Right column:* Binarized maps of activity from each of the three contrasts, displayed on the same surface. Although the regions are closely interdigitated, three parallel streams of regions can be observed, with the pseudoword-preferring stream most lateral (teal), followed by the face stream (blue), and scene stream (purple) most ventral. The boundaries of the LANG network defined by iFC (Fig. 2) are shown as black lines in each panel, which overlapped most consistently with the orthographic (Ps-words; teal) stream.

**Fig. 5:**
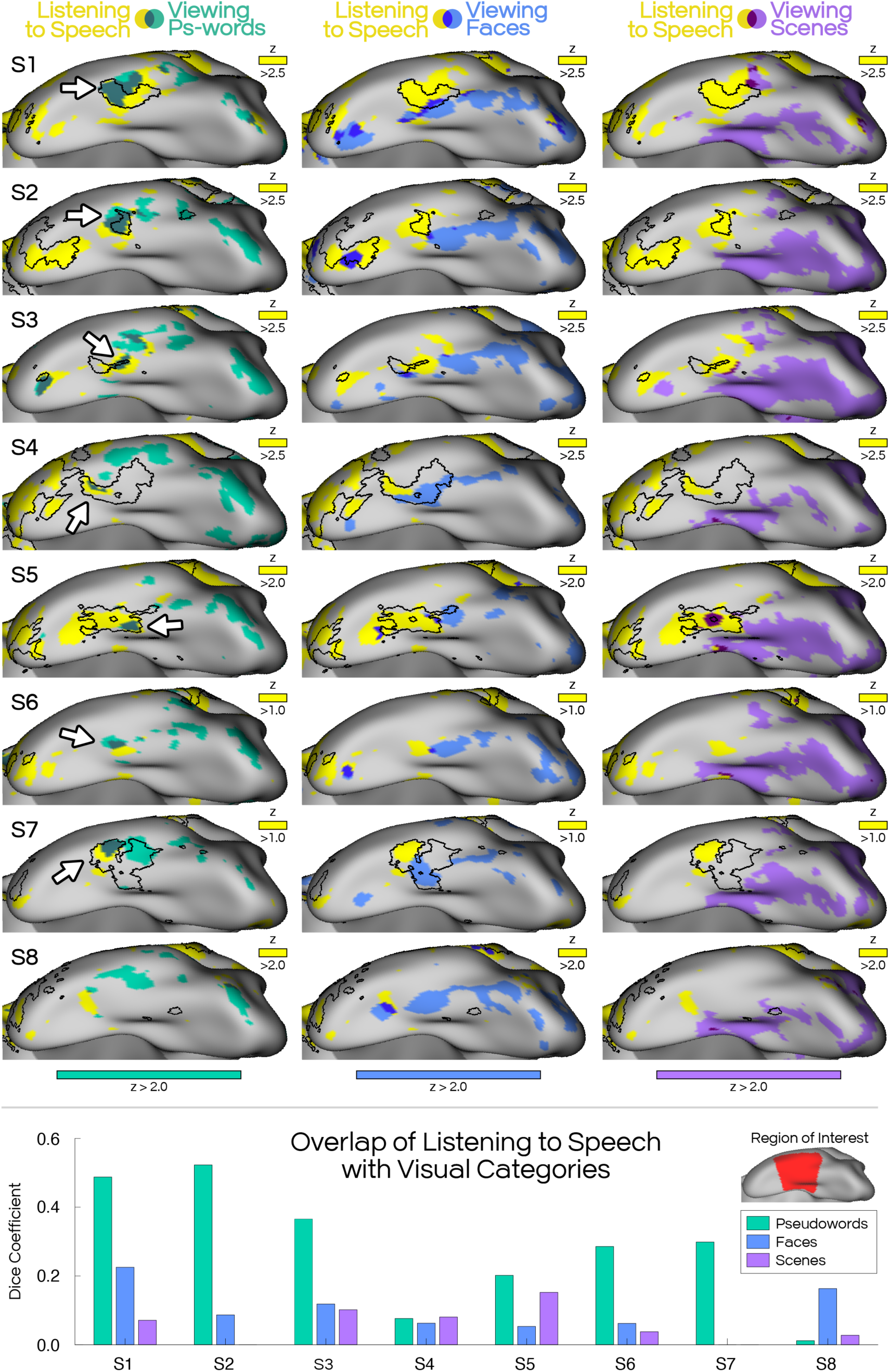
The basal LANG region activates both during listening to speech and viewing orthographic forms. *Top:* All but one participant demonstrated a mid-ITC region (see white arrows) that was active both when listening to speech (SPEECHLOC task; Fig. 3) and when viewing images of pseudowords (VISCAT task; Ps-words; Fig. 4). This overlap was consistently within the borders of the LANG network as defined by iFC (see black borders), despite each map being defined using independent data, tasks, and analysis approaches. Using the same thresholds in each participant, overlap with the transmodal LANG region was not seen as consistently for the face-(*middle column*) or scene-preferring (*right column*) streams. *Bottom:* The degree of overlap between speech-listening activity and each visual stream was quantified using Dice coefficients. Inset shows mask (red) used to isolate the mid-ITC regions of interest. Pseudowords (teal) demonstrated the greatest overlap with listening to speech, implying that auditory speech processing and viewing pseudowords converge more than other visual categories in mid-ITC.

Supporting this hypothesis, we observed that in 7 out of 8 participants, the most anterior region of the orthographic stream was also more active when participants listened to meaningful vs. distorted speech clips (SPEECHLOC task; see arrows in **Fig. 5**). In contrast, neither face-nor scene-related streams, immediately adjacent to the orthographic stream in the left mid-ITC, overlapped consistently across participants with regions active for listening to speech (see Dice coefficient graphs in **Fig. 5**). The coactivation of the basal LANG region by meaningless visual pseudowords and by meaningful auditory words suggests that this region displays a mixture of visual and transmodal language responses.

To further confirm that the anterior-most region of the orthographic stream was connected with the distributed LANG network, we chose seeds targeting ITC regions active during both viewing pseudowords and listening to speech. Correlation maps from these seeds were calculated using independent, held-out REST data (**Supp. Fig. S28**). The task maps of Participant S8 did not include an ITC region active for both pseudowords and speech, so a seed was chosen in the anterior-most pseudoword region, closest to speech activation. The correlation patterns were weak, possibly due to lower tSNR in the ITC, but were generally within or near the LANG network boundaries. Subjects S6 and S8 displayed maps that did not recreate the LANG network, in keeping with their main analyses, which failed to define a basal LANG region using iFC (**Fig. 2**).

### Replication of basal LANG region and parallel streams in the Natural Scenes Dataset

The above results demonstrated that (i) the ITC includes a basal LANG region in mid-ITC, (ii) category-preferring visual regions are arranged as parallel streams in posterior ITC, and (iii) the orthographic stream specifically overlaps with the basal LANG region. We replicated these findings in participants from the Natural Scenes Dataset (NSD).

As with the DBNO participants, a candidate language network was first identified for each NSD participant using resting-state iFC from seeds at or near lateral temporal cortex (**Fig. 6**). To build confidence in the estimated LANG network, we reproduced the network using seeds in multiple cortical zones (**Supp. Figs. S16–S21**). Three NSD subjects (NSD-S1, NSD-S2 and NSD-S4) demonstrated compelling evidence (i.e., correlations of r > ∼0.4 from multiple seeds) of basal LANG regions at or near mid-ITC, replicating our findings from the DBNO dataset. In these subjects, the presence of a basal LANG region was confirmed using data-driven clustering (black outlines in **Fig. 6**), which also confirmed this region was left-lateralized (**Supp. Fig. S29**). Two other subjects (NSD-S3 & NSD-S6) displayed some mid-ITC connectivity to LANG for a subset of the seeds (**Supp. Figs. S18–S20**). Note that signal dropout was more pronounced in the single-echo NSD data (see tSNR maps in **Supp. Fig. S15**), which may have influenced observable iFC and task activity (**Supp. Fig. S30**). Subsequent analyses therefore focused on the three participants with successful replications of the basal LANG region based on iFC (NSD-S1, NSD-S2, NSD-S4).

We plotted the maps from a visual category localizer task (‘fLoc’ task; similar to VISCAT), performed by the NSD participants. In all three NSD subjects, the pseudowords contrast identified a similar orthographic stream of active ITC regions (**Fig. 6**). This stream again spanned from the occipital pole to about halfway down the long axis of the ITC, replicating our findings from the DBNO participants (**Fig. 3**). These maps also indicated the presence of interdigitated regions broadly arranged into three parallel visual streams (akin to **Fig. 4**), with the scene-related stream being most medial, the orthographic stream most lateral, and the face-related stream in between. However, the pattern was less clear compared to the DBNO data (e.g., in NSD-S2 the orthographic stream was both lateral and medial to the face-related stream). Replicating our main findings, the anterior-most pseudoword-active region again overlapped with the borders of the basal LANG region in all three individuals.

**Fig. 6:**
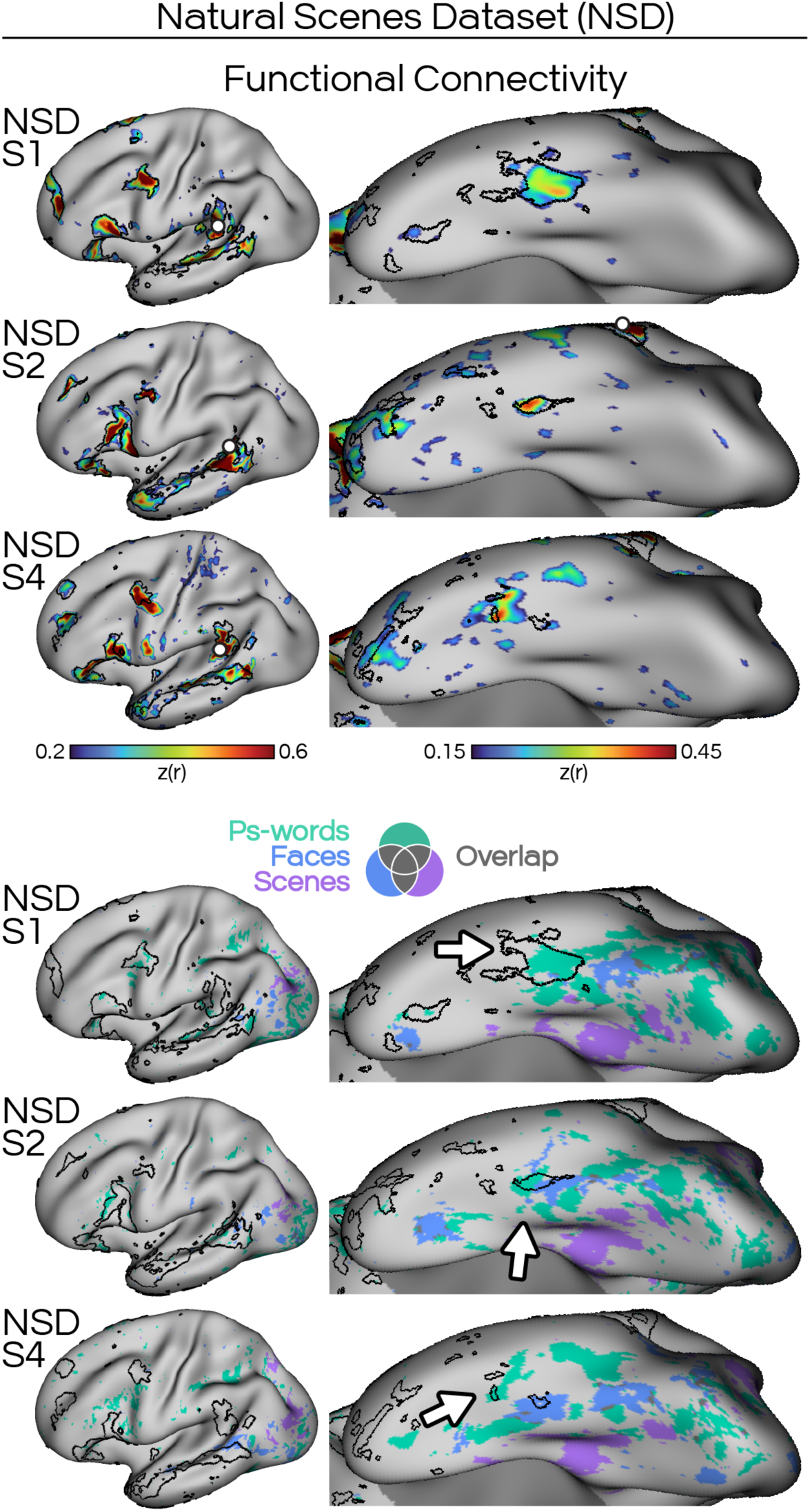
Replication that the orthographic stream converges on a basal LANG region in the Natural Scenes Dataset (NSD). *Top:* The presence of a basal LANG region was confirmed in 3 participants (*upper panel*) using iFC (remaining subjects shown in **Supp. Fig. S29**). *Bottom:* This basal LANG region overlapped with the anterior-most region of the orthographic stream (*lower panel; white arrows*) revealed by the NSD fLoc task, in which participants viewed pseudowords (teal), faces (blue) and scenes (purple).

### LANG network connectivity explains differences in visual selectivity along the orthographic stream

Our findings support that the anterior-most region of the orthographic stream (i.e., the basal LANG region) displays properties of both transmodal and unimodal visual systems, in that it responds to both meaningless visual words and when listening to speech (**Fig. 5**; and see Li et al., 2024). Importantly, this mixture of responses distinguishes the anterior-most region from the remainder of the orthographic stream, implying that the network connections to LANG might explain the functional heterogeneity observed within the stream.

Given that listening to speech activated only the anterior-most orthographic stream region, we asked whether other types of visual stimuli could selectively activate the posterior regions, in a double dissociation. Previous work had reported that the VWFA responds more to real words and consonant strings compared to samples of foreign scripts (e.g. Chinese and Hebrew characters; Baker et al., 2007), suggesting that familiarity could drive responses within the VWFA. Therefore, we investigated how different regions of the orthographic stream responded to stimuli that increasingly diverged from familiar written words.

We analyzed the VWFACAT task data, assessing responses as DBNO participants viewed written real words, pseudowords, consonant strings, and a foreign script (i.e. modified Japanese characters), compared to a baseline condition of viewing line drawings (**Supp. Fig. S31**). The maps were thresholded, binarized, and plotted on top of the orthographic stream as defined using the pseudowords condition of the VISCAT task (**Figs. 4 & 5**). Viewing pseudowords activated the same set of ITC regions in both tasks, and the same set of regions also activated during viewing real words and consonant strings (**Supp. Fig. S32**). However, foreign script stimuli activated only the posterior regions of the orthographic stream and failed to activate the more anterior region (**Fig. 7**).

Analysis of the NSD participants provided further evidence for this dissociation. In the NSD dataset, participants viewed pseudowords as well as strings of written numbers. Viewing number strings activated posterior regions of the orthographic stream but not the anterior region connected to the LANG network (**Fig. 7** & **Supp. Fig. S33**). Thus, the most anterior orthographic stream region was distinguished from the posterior orthographic regions both by its responsiveness to meaningful speech (**Fig. 5**) and by its lack of response to unfamiliar foreign scripts (**Fig. 7**). This implies that the connections to the distributed association regions of LANG could support the recognition of familiar orthographic visual forms in the basal LANG region.

As a final confirmation that this anterior-most region was connected to the LANG network, in the NSD data we targeted seeds to mid-ITC regions that were active during presentation of written pseudowords but not during presentation of number strings (teal in **Fig. 7**). We then plotted the iFC maps using held-out NSD data for participants NSD-S1 and NSD-S3 (who provided sufficient data for replication), as well as the original data for NSD-S2 (who provided only sufficient data for the discovery dataset; **Supp. Fig. S34**). The correlation maps robustly replicated the estimates of the LANG network from lateral seeds (**Supp. Fig. S29**), further supporting that the anterior-most orthographic stream region is connected to LANG.

**Fig. 7:**
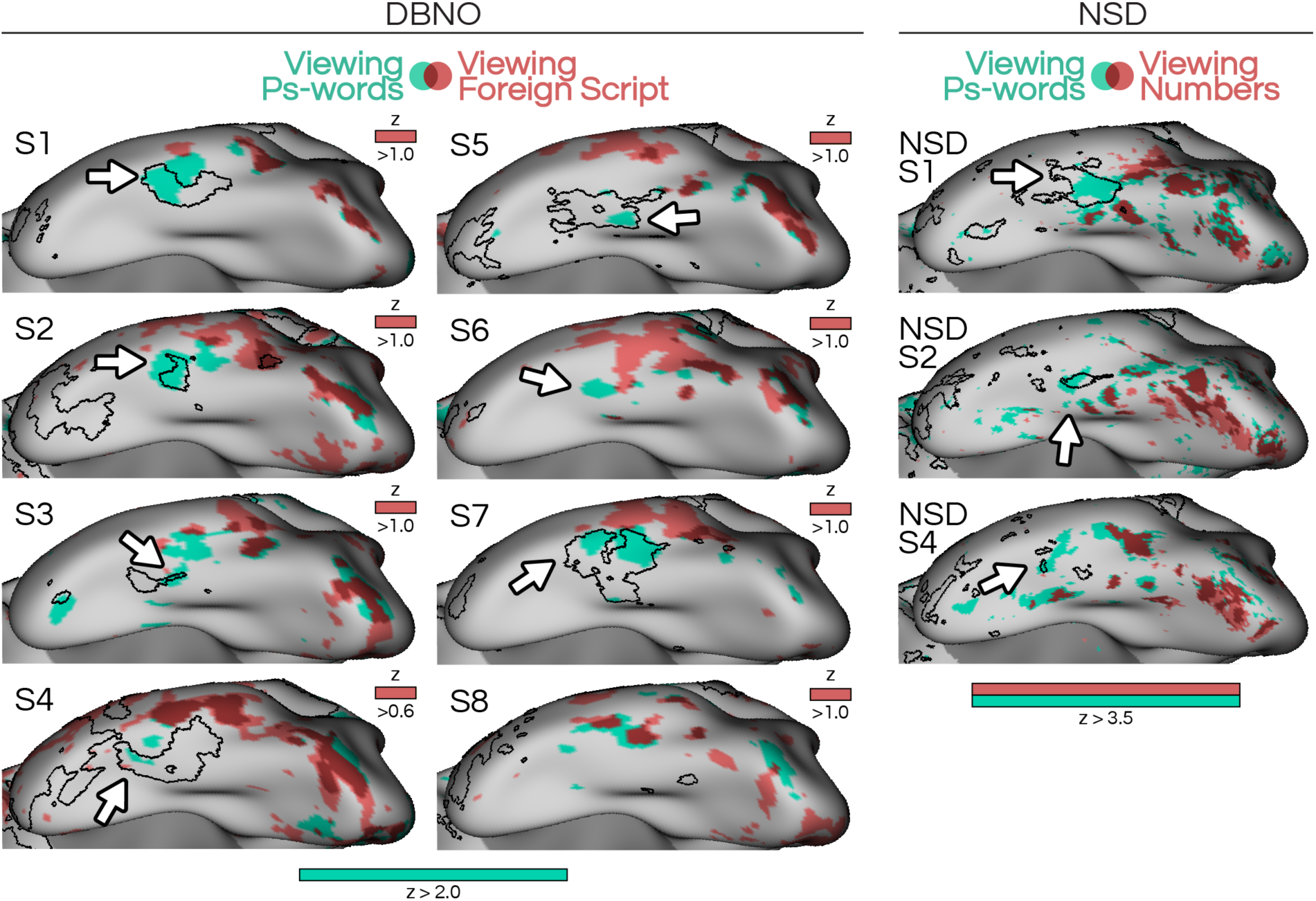
Connections to the LANG network differentiate visual responses along the orthographic stream. In all panels, the black borders delineate the LANG network (black borders) as defined using intrinsic functional connectivity (iFC) from resting-state data. In the DBNO and NSD data, viewing pseudowords activated the full set of orthographic stream regions, including the anterior-most region that overlapped with the basal LANG region. However, viewing visual stimuli that less closely resembled writing, including an unfamiliar foreign script (modified Japanese characters; red in left columns) or number strings (red in right column), only activated the more posterior parts of the orthographic stream (dark red overlap). These results confirm that the orthographic stream can be double dissociated, with the anterior-most region showing selective responses when listening to speech, while the posterior regions show selective responses for less familiar visual stimuli that resemble orthographic forms. Importantly, this differentiation of function is precisely predicted by the borders of the LANG network. All DBNO participants are displayed with z > 2.0 for viewing pseudowords (teal), and all but DBNO participant S4 (z > 0.6) are displayed with z > 1.0 for viewing foreign script (red). All NSD participants are displayed with z > 3.5 for both conditions.

## Discussion

We characterized the neuroanatomy of reading and language comprehension using precision functional mapping. First, we confirmed the existence of an inferotemporal (“basal”) region that was functionally connected to the distributed language network (LANG; **Fig. 2**). Next, we delineated a stream of visual regions that responded when a participant viewed orthographic forms, positioned next to parallel visual streams supporting face and scene processing (**Fig. 4**). Next, we showed that only the anterior-most of these orthographic stream regions, which was located in mid-ITC, reproducibly aligned with the basal LANG network region (**Fig. 4 & 4**). This mid-ITC orthographic stream region demonstrated mixed selectivity: it responded to visual displays of meaningless character strings (e.g., pseudowords, consonant strings), implying a visual function, but also to aurally presented intelligible speech, implying a transmodal language function (**Fig. 5**). In contrast, the posterior orthographic stream regions activated for other stimuli that shared visual features with written words but were less familiar, including foreign scripts and number strings (**Fig. 7**). We also reproduced the results in a separate dataset collected at 7 T (**Figs. 6 & 7**). The results suggest that the basal LANG region could serve as a nexus for reading, where sensory information from unimodal visual hierarchies interfaces with processes supported by the transmodal LANG network.

### The distributed language (LANG) network includes a basal region in mid-ITC

The classic language network is predominantly considered to include perisylvian regions in inferior frontal and lateral temporal cortex (e.g., Geschwind, 1970). However, there is now strong evidence that the network includes regions in frontal, temporal, and midline association zones (Dronkers et al., 2004; Fedorenko et al., 2010), which mirrors the organization of other association networks (Braga et al., 2020). Here, the use of multi-echo data allowed us to reliably define an ITC region of the LANG network using iFC (**Fig. 2**). This is important because the definition of the network, including its distributed architecture, was based on intrinsic (i.e., task agnostic) correlations of brain signal, and was therefore not confounded by the specifics of a given cognitive task. The presence of this basal LANG region was then confirmed by task activation during listening to speech (**Fig. 3**) and replicated in an independent dataset (**Fig. 6**). This finding supports multiple prior reports of a basal ITC region that responds to both spoken and written language, and that can lead to language-related deficits when stimulated or lesioned (Lüders et al., 1986; Sharp et al., 2004; Krauss et al., 1996; Ulvin et al., 2017; Snyder et al., 2023; Nobre et al., 1994; Forseth et al., 2018; Li et al., 2024). Further, ITC responses have also been reported in tasks including picture naming and naming based on spoken descriptions (Forseth et al., 2018; Snyder et al., 2023). These studies together provide robust evidence that LANG supports transmodal aspects of language and includes a region in mid-ITC, even though this part of the brain is more commonly associated with visual processing. Speculatively, the presence of these mid-ITC connections to the distributed LANG network could be a pathway through which visual input from the visual streams can lead to linguistic processing in the LANG network during reading.

### Category-selective visual regions are organized as parallel streams in posterior ITC

Our maps of the ITC also indicate that distinct category-selective regions may follow a shared organizational motif. We observed three parallel “streams” of regions extending in an arc from near the occipital pole towards the temporal pole (**Fig. 4**) that were each preferentially responsive to a particular visual category, i.e. faces, scenes, or orthographic forms. This description is consistent with previous work demonstrating multiple category-responsive visual regions in the ITC, organized as category-selective ‘archipelagoes’ (Kanwisher et al., 2010) rather than a single visual word form area (White et al., 2019) or fusiform face area (Kanwisher et al., 1997; Grill-Spector, 2003; Hasson et al., 2002; Levy et al., 2001; Malach et al., 1995; Weiner & Grill-Spector, 2010; Sayres & Grill-Spector, 2008). Different regions within these archipelagoes may represent different stages of a visual processing hierarchy (Van Essen & Maunsell, 1983; Goodale & Milner, 1992; Vinckier et al., 2007).

### The orthographic stream converges on the basal LANG region

We saw compelling evidence that the anterior-most region of the orthographic stream was functionally different from more posterior regions. First, the anterior region overlapped with the basal LANG region, defined using iFC, while posterior regions did not (**Figs. 4 & 5**). Second, this overlap with LANG was specific to the orthographic stream; anterior regions of the face- and scene-related streams did not overlap with LANG to the same degree (**Fig. 5; lower row**). Third, the anterior orthographic stream region resembled other LANG regions in its responsiveness during listening to speech (**Figs. 3 & 5**). Fourth, the full orthographic stream showed activity for real words, pseudowords, and consonant strings (i.e., images that included characters from a familiar Roman script), while the anterior region did not activate for foreign scripts and number strings (**Fig. 7**). This implies that the posterior regions are less selective in their responses and are likely sensitive to visual stimuli that share similar visual features to familiar writing, such as written numbers or unfamiliar characters. In contrast, the anterior region more selectively activates for visual stimuli that more closely resemble readable words and are deeply familiar through life-long, extensive experience with reading. Thus, our findings regarding the anterior orthographic region match the orthographic/speech activations of the VWFA described by Li et al. (2024). An intriguing hypothesis raised by the present results is that the network connections to the LANG network, which encompass multiple association areas including in frontal and temporal lobes, are fundamental to the acquisition of this experience-dependent specialization for recognizing written word forms.

### The basal language region as a nexus between vision and language

Our results suggest that the basal LANG region’s functional properties may be explained by the region’s precise positioning at the confluence between the distributed LANG network and the orthographic stream. It has been proposed that the mid-ITC acquires a preference for familiar orthographic forms as an individual learns how to read (Dehaene et al., 2010; Saygin et al., 2016), and that this process requires access to both the orthographic stream and transmodal language areas. For instance, the dual mode theory of reading argues that individuals learn to read by associating unfamiliar written words with familiar phonemes (Coltheart et al., 2001). Consistent with this theory, our results demonstrate that within the orthographic stream, the basal LANG region has the greatest selectivity for familiar, readable scripts and is also the only stream region activated transmodally by listening to speech (**Figs. 4 & 6**). Repeated engagement of the transmodal LANG network during the visual perception of orthographic forms could strengthen connections between the mid-ITC and perisylvian LANG regions involved in phoneme processing and articulation (see Dehaene et al., 2015). The distributed connections to the LANG network revealed here could support this widespread dissemination of reading relevant information. Similarly, more local connections with the visual stream could explain the basal LANG region’s responsiveness to artificial written language samples (e.g. pseudowords). Pseudowords may be processed similarly to real words, in that common or pronounceable letter combinations may be recognized and preferentially processed by the basal LANG region. However, these artificial stimuli lack syntactic or semantic features and thus do not activate the broader LANG network (**Supp. Fig. S27**).

Interestingly, the presence of the orthographic stream could also determine or relate to the basal LANG region’s development. Some evidence suggests that the mid-ITC may not respond to language, whether heard or read, in illiterate individuals (Monzalvo & Dehaene-Lambertz, 2013). Thus, while connectivity with the LANG network offers the potential for a mid-ITC language region to form, the formation (or expansion) of this region may follow acquisition of reading. Supporting such a link, we observed that both systems were lateralized to the same hemisphere, with larger regions on the left than the right (**Supp. Figs. S23-S25 & S27**). In contrast, face- and scene-processing regions were more bilaterally or right-hemisphere distributed (**Supp. Fig. S26**). Our results support that the orthographic stream itself is left lateralized, including more posterior regions, rather than just the basal LANG region (see also White et al., 2019). The left lateralization of language may be present in the early years of life (Dehaene-Lambertz et al., 2002; Dehaene-Lambertz et al., 2010; Holland et al., 2007; Szaflarski et al., 2006), even in the ITC, prior to the acquisition of reading (Saygin et al., 2016; see also Arcaro & Livingstone, 2021). If so, lateralization of language could subsequently influence lateralization of the whole orthographic stream through top-down influences.

Another possible explanation for the observed responses of the basal LANG region is that this region’s responses are shaped by articulation. For most of our participants, legible orthographic stimuli activated the basal LANG region (**Fig. 5**) but did not robustly activate the rest of the LANG network (see lateral views in **Supp. Fig. S25**) to the degree that listening to speech did (**Supp. Fig. S24**). In some subjects (e.g., see S3, S6 & S7 in **Supp. Fig. S25**), pseudowords instead activated lateral regions typically associated with articulatory processes (e.g., phoneme encoding; de Heer et al., 2018). These putative articulatory regions were located between the LANG network and the motor strip in the frontal lobe, and between the LANG network and auditory cortex in the temporal lobe. We have previously suggested that a similar arrangement of regions can be separated from the LANG network using iFC (see the intermediate, “INT”, network in the seventh and eighth figures of Braga et al., 2020). A plausible interpretation for these responses to familiar character strings is that participants are mentally articulating the characters (e.g., in inner speech) as they read them, including for difficult to pronounce consonant strings or pseudowords. The activation of the basal LANG region in isolation to the rest of the LANG network could similarly represent recruitment of articulatory or phonemic processes during the reading of pseudowords and consonant strings. The fact that we observed only partial overlap in the ITC between the orthographic stream and the LANG network in some individuals (e.g., see subjects S5 and S6 in **Fig. 5**), suggests that there may be further heterogeneity of responses within the basal LANG region.

The present results contribute to ongoing debate about the nature and specialization of word processing in the ITC (Vogel et al., 2012b). Although traditionally described as a single area (the visual word form area, or VWFA; Cohen et al., 2002), word processing in the ITC may in fact take place over a sequence of closely related stages through the orthographic stream. The similarity of responses across stages could explain diverging reports of the VWFA’s membership in distinct distributed networks: individualized studies of functional anatomy link the VWFA with perisylvian language areas (Stevens et al., 2017; see also Saygin et al., 2016; Li et al., 2020; 2024), while group-average studies have linked the VWFA to the dorsal attention network (Vogel et al., 2012a; Chen et al., 2019). This matches a framework which subdivides the VWFA into at least two regions with different orthographic preferences (e.g., VWFA-1 and VWFA-2; White et al., 2019; Woolnough et al., 2021). Our results may help reconcile these interpretations by attributing these different stimulus preferences to different regions along the orthographic stream that house different network connections (**Fig. 5**). The more posterior regions of the orthographic stream are likely to show connectivity with the dorsal attention network, as Vogel et al. (2012a) and Chen et al. (2019) have indicated, given their position in more posterior ITC. We are investigating this possibility.

Individualized analyses that consider fine-scale functional topography may be necessary to fully characterize the responses and network connections throughout the visual processing streams.

### Limitations

The observation of parallel streams displaying a preference for the visual stimulus category (faces, scenes, words) is in slight conflict with previous work that has focused on a specific, single region of ITC that displays category selectivity, such as the fusiform face area (FFA), parahippocampal place area (PPA), and VWFA. In the terminology of Kanwisher (2010), we observed category-preferring ‘archipelagoes’, rather than single regions, for each of the different image categories. We further showed that these archipelagoes formed parallel streams that radiated away from the occipital pole (**Fig. 5**). These streams could be due to the specifics of the stimuli used here. Although each image was cropped to a specific bounding box, the orthographic stimuli took up the least of the white space within this box, while the scenes images took up the most space. These features might affect the degree to which the visual field was engaged by each image type, which could explain why categories were distinguishable even in earlier visual areas. Arguing against this, we also observed evidence for more posterior visual regions being recruited in the NSD data, using a task in which stimuli were presented against a noise background. However, the posterior portions of the streams, as well as the parallel organization of different streams, were less clear in NSD participants compared to in DBNO participants. Similarly, the stimulus conditions could have elicited reliable differences in saccades, which were not measured here. These differences are arguably inherent to the processing of different types of visual stimuli, and we did not seek to investigate this specifically. Nonetheless, these factors should frame consideration of the parallel streams.

## Conclusions

Our study indicates that visual and language systems involved in reading may interface at an inferotemporal language region that serves as an important nexus capable of integrating features of both unimodal and transmodal processing (**Fig. 1**). The functional connectivity and mixed selectivity of this language region distinguishes it from the two systems it engages with. These findings clarify the circuit properties that underly the neuroanatomy of reading, by highlighting that the convergence of vision and language in reading is anatomically supported by the convergence between visual processing streams and the transmodal language network at the basal language region (**Fig. 1**).

We believe our work provides new insights into how brain network architectures may support the translation of sensory information into higher-level associative cognitive processes.

## Funding

This work was supported in part by National Institute of Mental Health grant R00 MH117226 (R.M.B.); an Alzheimer’s Disease Core Center grant (P30 AG072977) pilot award (R.M.B.) from the National Institute on Aging to Northwestern University, Chicago, Illinois; National Institute of Neurological Disorders and Stroke grant T32 NS047987 (to J.J.S., A.M.H.); and the William Orr Dingwall Foundations of Language Fellowship (J.J.S.). The content is solely the responsibility of the authors and does not necessarily represent the official views of the National Institutes of Health.

## Acknowledgments

This research was supported in part through the computational resources and staff contributions provided for the Quest high performance computing facility at Northwestern University which is jointly supported by the Office of the Provost, the Office for Research, and Northwestern University Information Technology, and the Center for Translational Imaging at Northwestern University.

## Data and Code Availability

All data needed to evaluate the conclusions in the paper are present in the paper. Data files can be provided upon reasonable request.

## Author Contributions

J.J.S., M.L., & R.M.B. designed the study; J.J.S., M.L., A.M.H., & R.M.B. collected the data; J.J.S. and M.L analyzed the data with input from Z.M.S., M.M.M., & R.M.B; J.J.S. & R.M.B wrote the original draft; J.J.S., M.L., A.M.H., Z.M.S., M.M.M., & R.M.B edited and wrote the manuscript; R.M.B. secured funding.

## Declaration of Interests

The authors declare no competing interests.

## Notes

### Competing Interest Statement

The authors have declared no competing interest.

### Summary of Updates

Corrected label mentions in two figure captions to match label in figures

