## Supplementary Figures for "The ventral visual stream for reading converges on the transmodal language network"

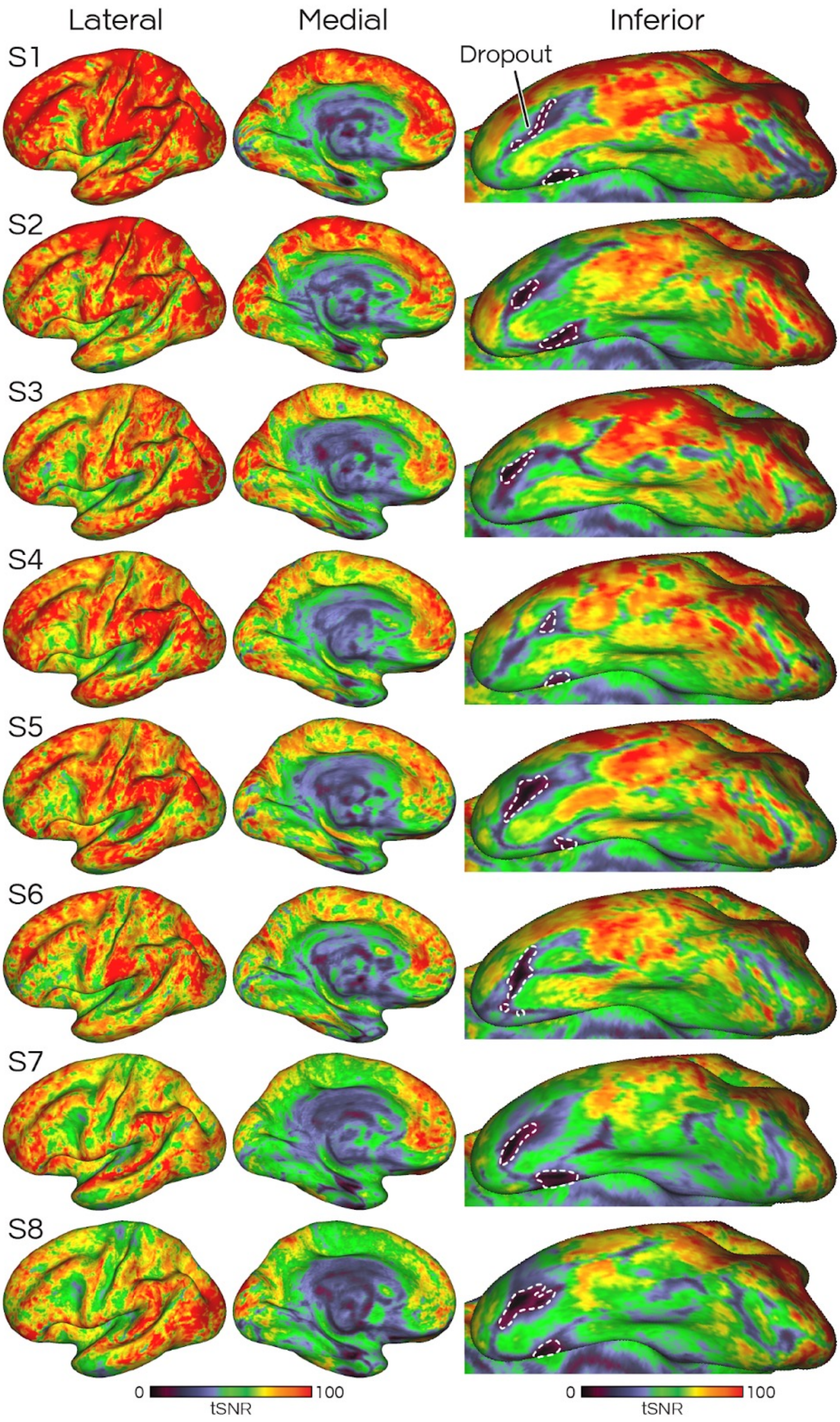


**Supp. Fig. S1: Temporal signal-to-noise ratio (tSNR) of the multi-echo functional MRI data was high across all DBNO participants, including in inferior temporal regions central to our analyses.** Rows denote individual participants, ordered from top to bottom by descending mean tSNR. For the inferior view (right), left/right is anterior/posterior and up/down is lateral/medial; white dashed lines identify hand-drawn regions of particularly low signal (tSNR < ~40). tSNR was calculated using all runs except the held-out half of resting-state runs.

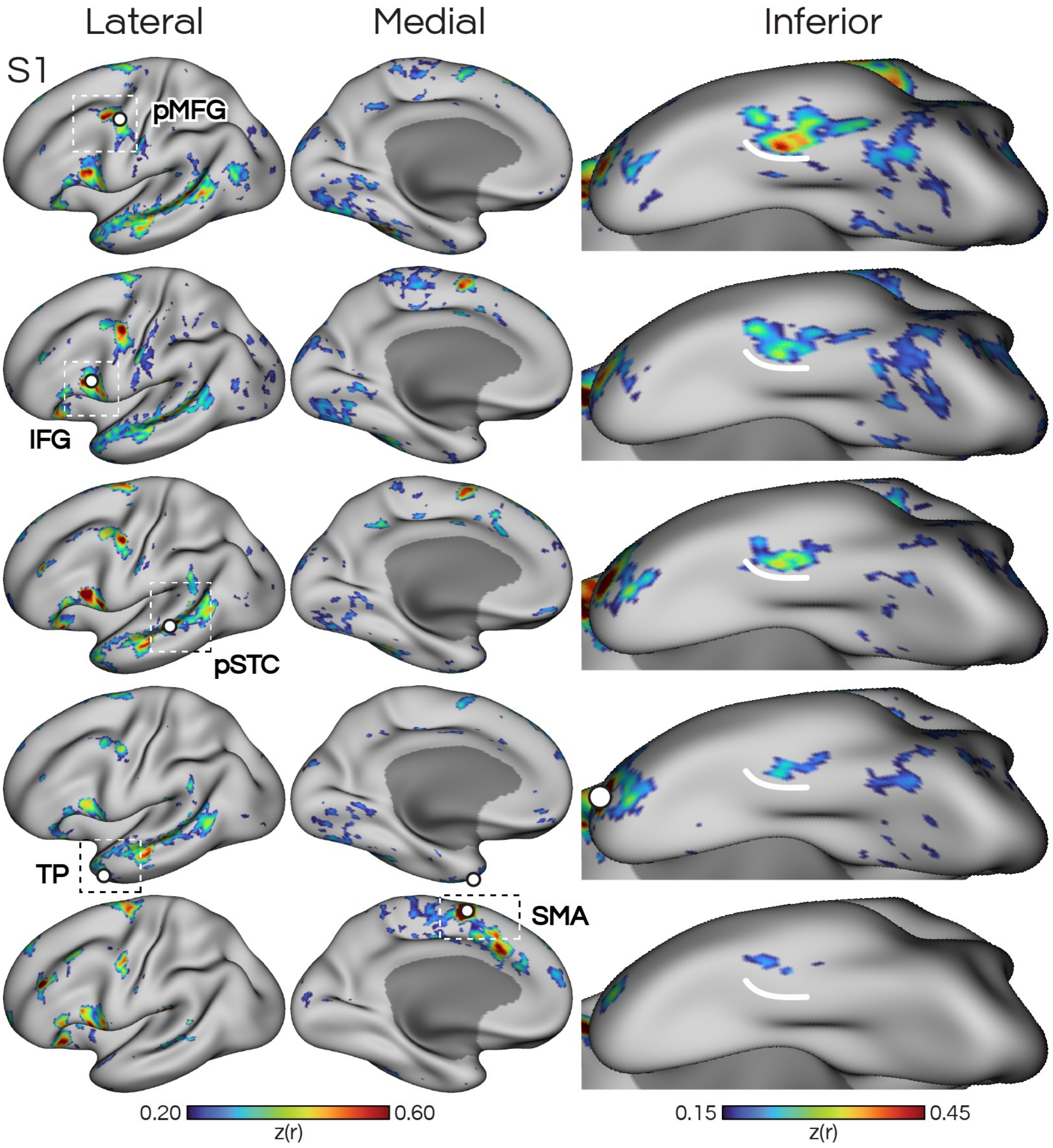


**Supp. Fig. S2: A candidate language network is defined in subject S1 using functional connectivity (iFC) from seeds in five cortical zones.** The multiple estimates provided converging evidence that the distributed LANG network includes a region approximately halfway down the inferior temporal cortex. Maps depict Z-normalized Pearson correlations. Z display thresholds were lowered for the inferior views to account for lower signal (see **Supp Fig. S1**). pMFG: posterior middle frontal gyrus, IFG: inferior frontal gyrus, pSTC: posterior superior temporal cortex, TP: temporal pole, SMA: supplementary motor area.

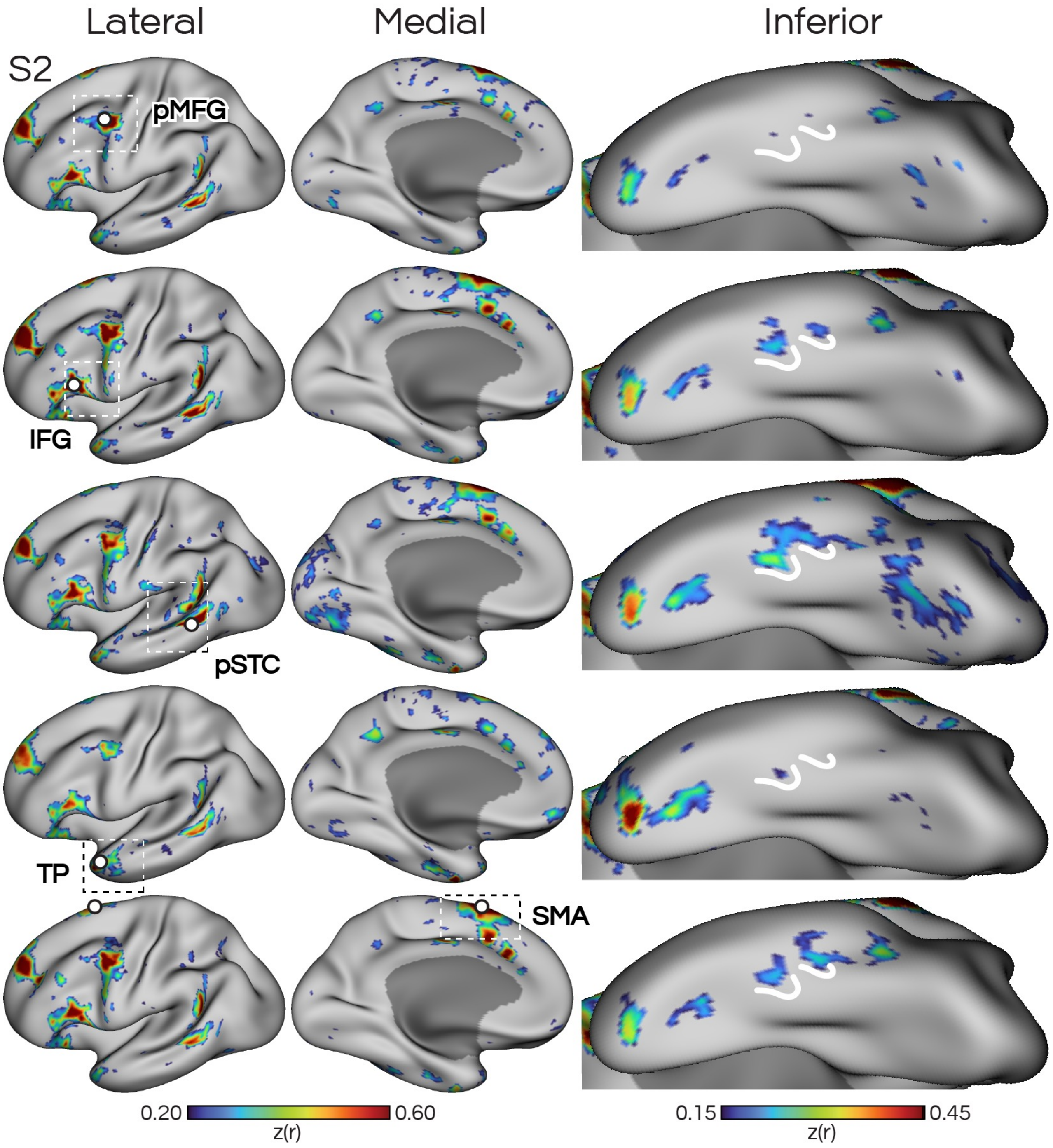


**Supp. Fig. S3: A candidate language network defined in subject S2 using functional connectivity from seeds in five cortical zones.** Formatted according to **Supp. Fig. S2**.

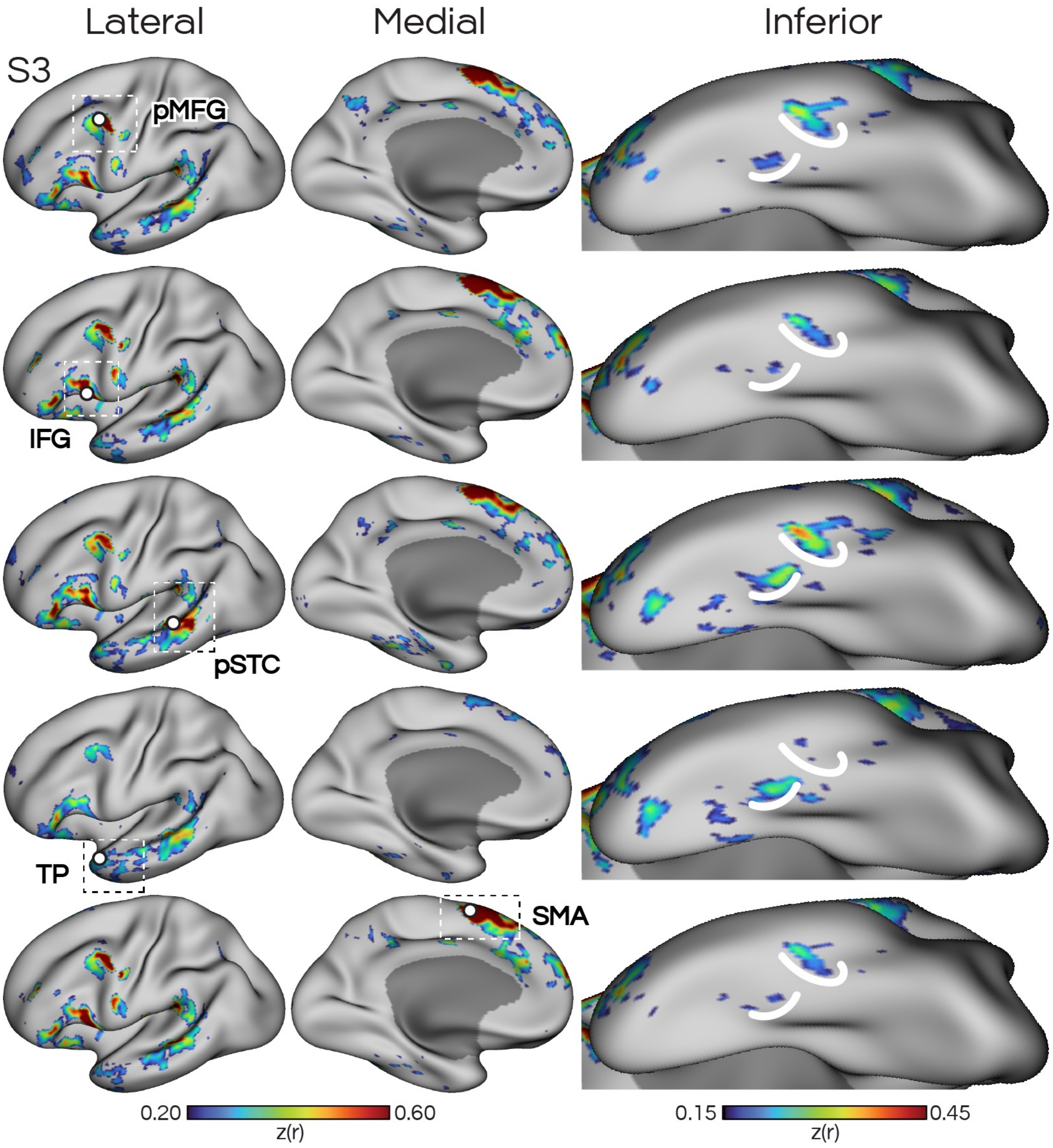


**Supp. Fig. S4: A candidate language network defined in subject S3 using functional connectivity from seeds in five cortical zones.** Formatted according to **Supp. Fig. S2.**

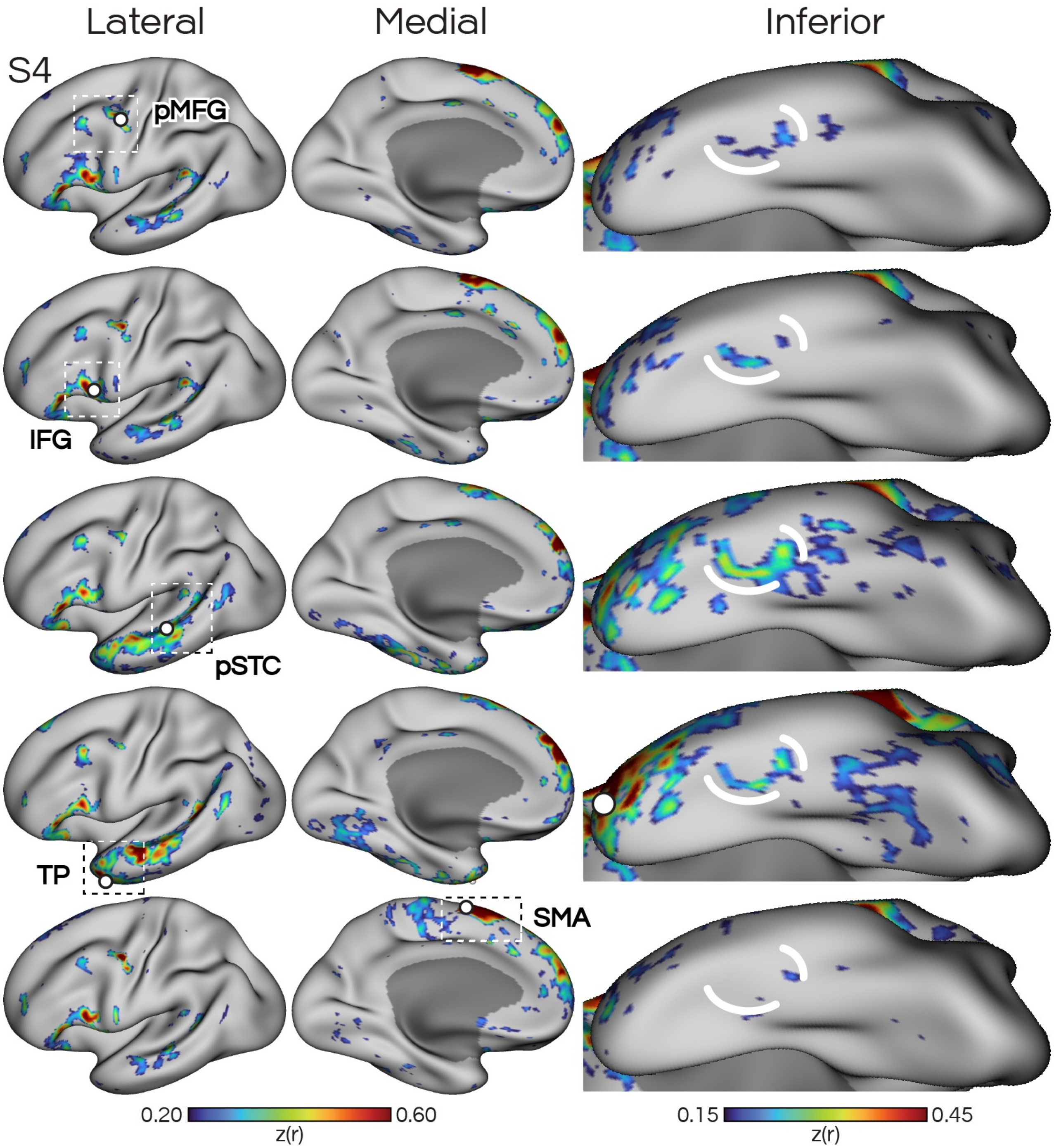


**Supp. Fig. S5: A candidate language network defined in subject S4 using functional connectivity from seeds in five cortical zones.** Formatted according to **Supp. Fig. S2.**

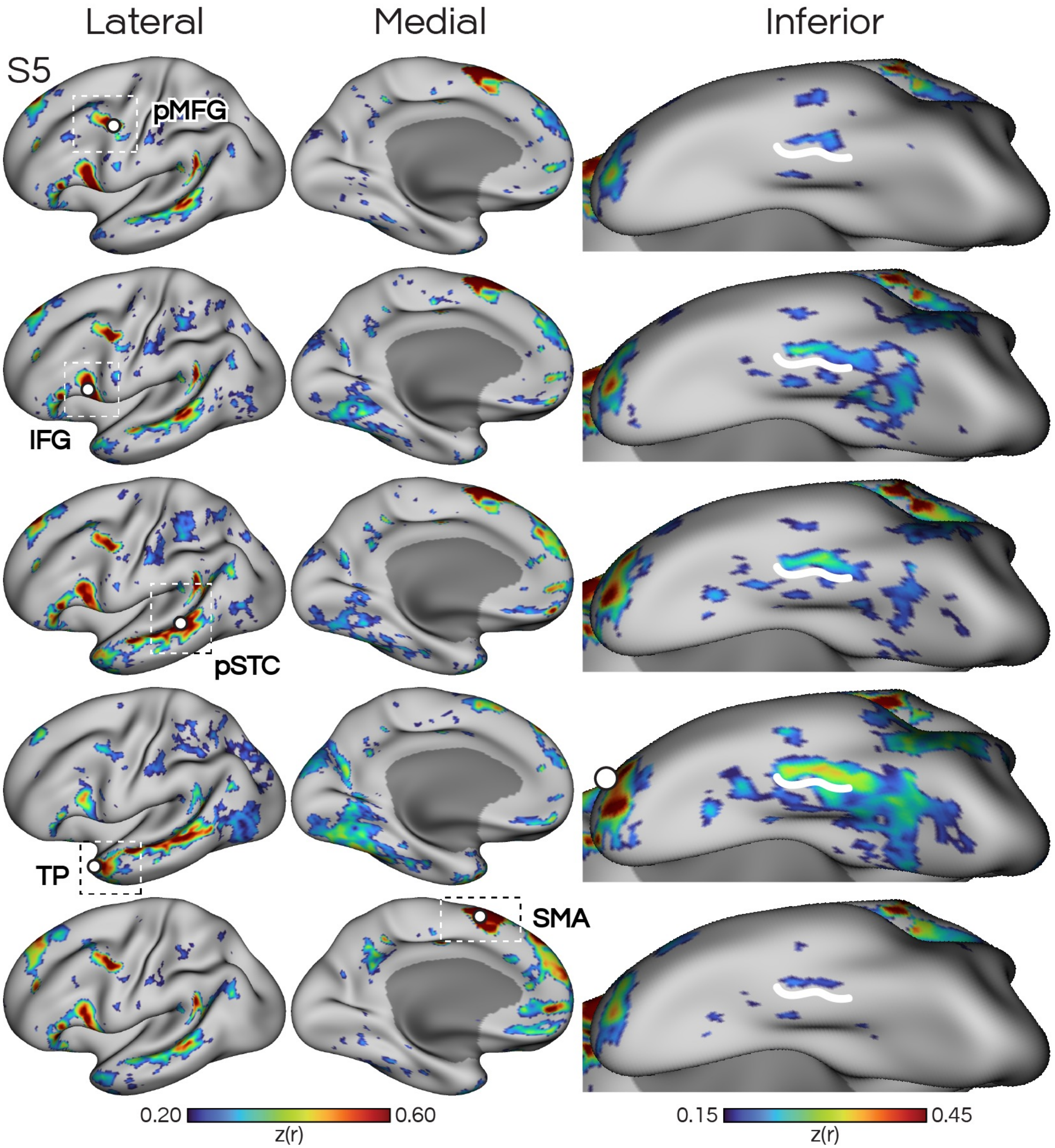


**Supp. Fig. S6: A candidate language network defined in subject S5 using functional connectivity from seeds in five cortical zones.** Formatted according to **Supp. Fig. S2.**

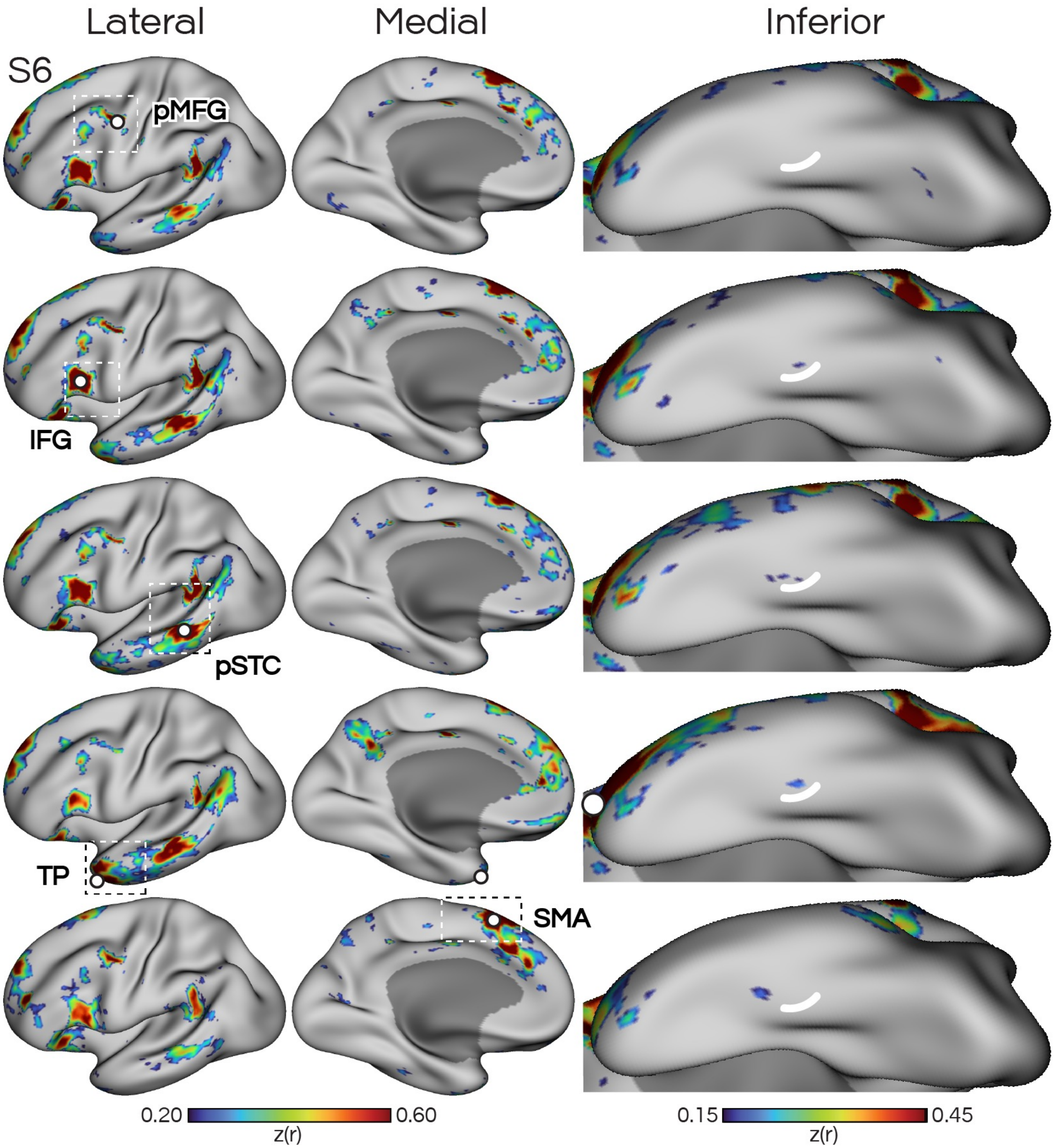


**Supp. Fig. S7: A candidate language network defined in subject S6 using functional connectivity from seeds in five cortical zones.** Formatted according to **Supp. Fig. S2.** Limited evidence of an inferior temporal LANG network region was seen in this subject, potentially due to lower data quality.

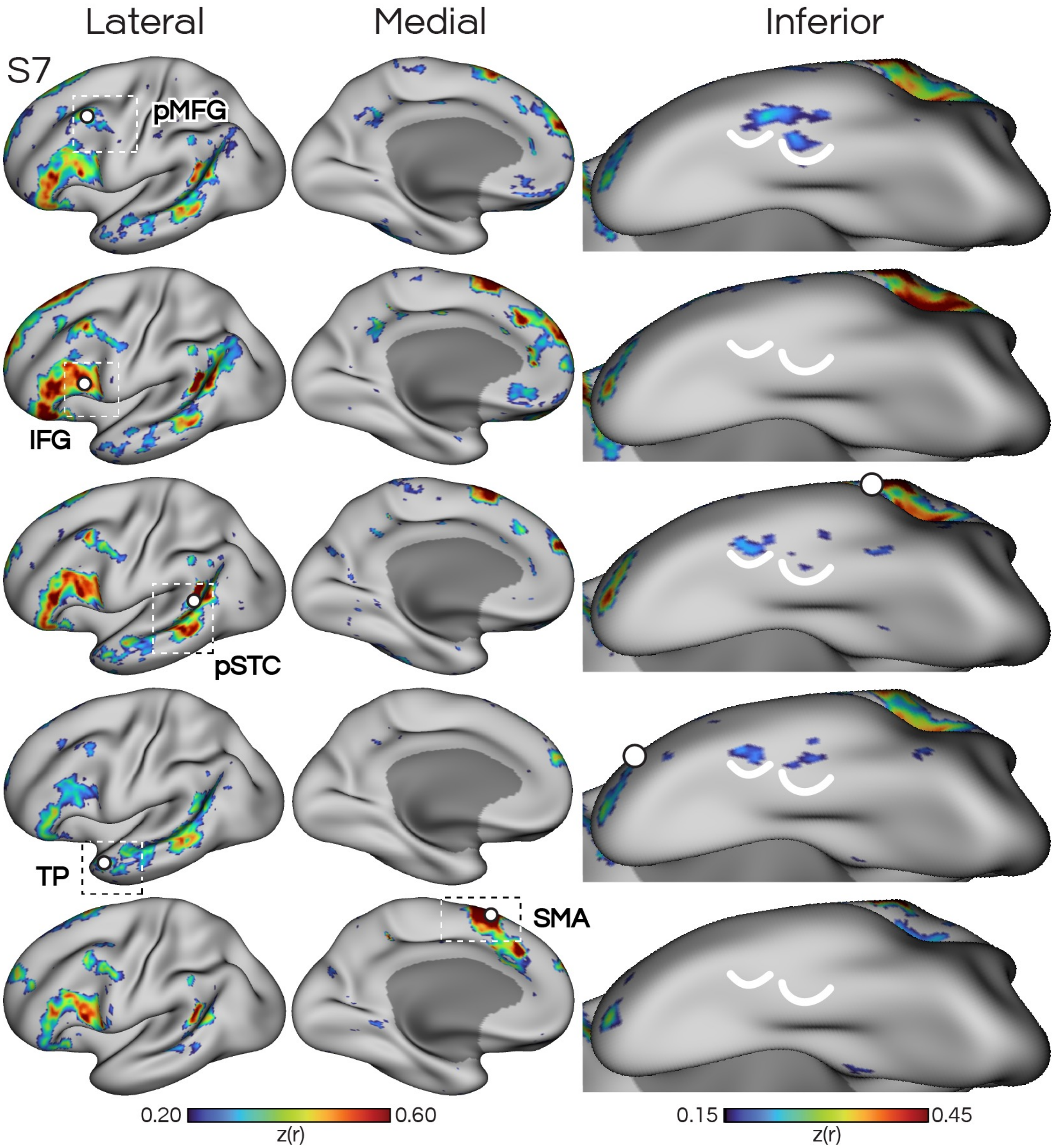


**Supp. Fig. S8: A candidate language network defined in subject S7 using functional connectivity from seeds in five cortical zones.** Formatted according to **Supp. Fig. S2.**

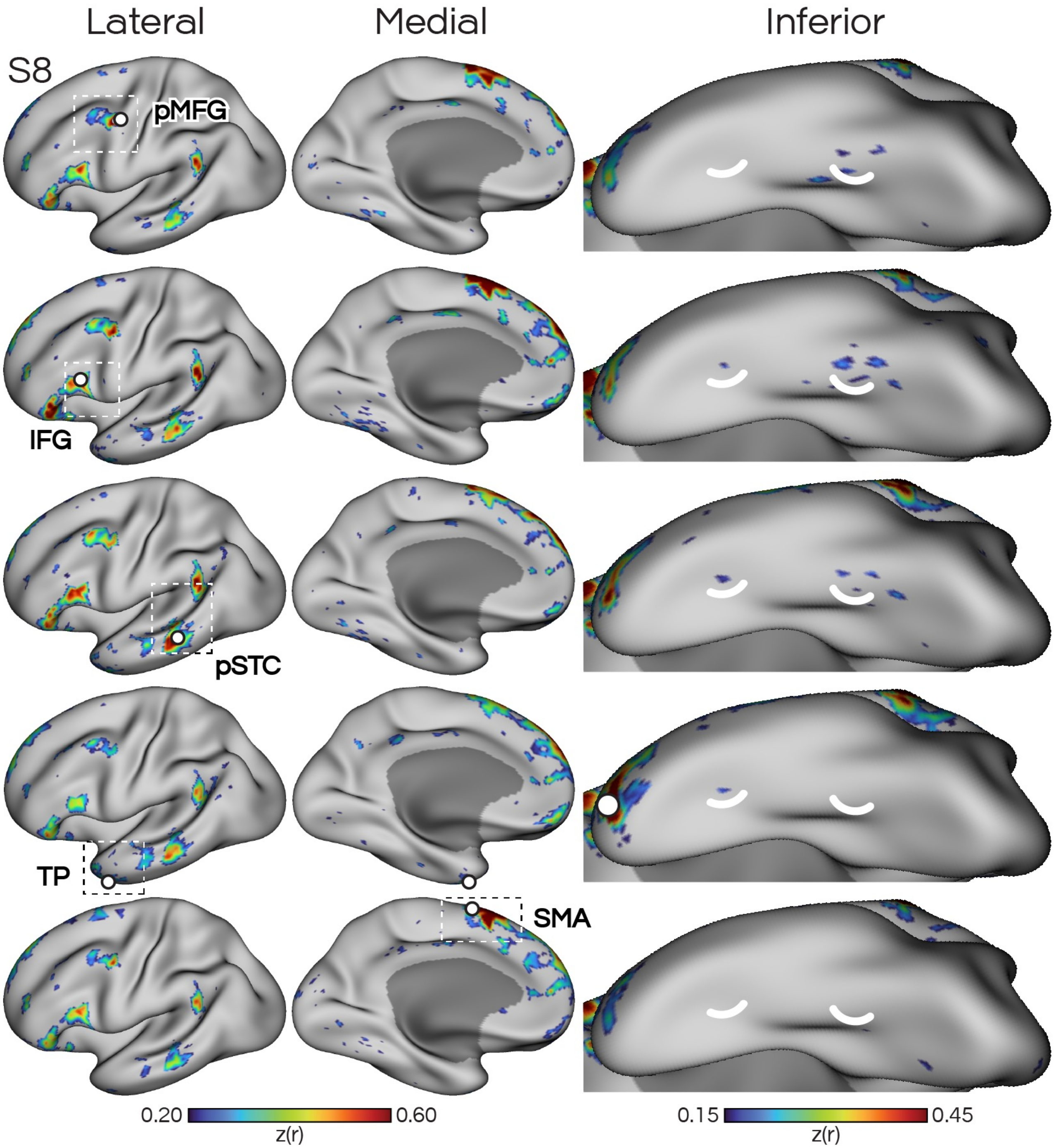


**Supp. Fig. S9: A candidate language network defined in subject S8 using functional connectivity from seeds in five cortical zones.** Formatted according to **Supp. Fig. S2.** Limited evidence of an inferior temporal LANG network region was seen in this subject, potentially due to lower data quality.

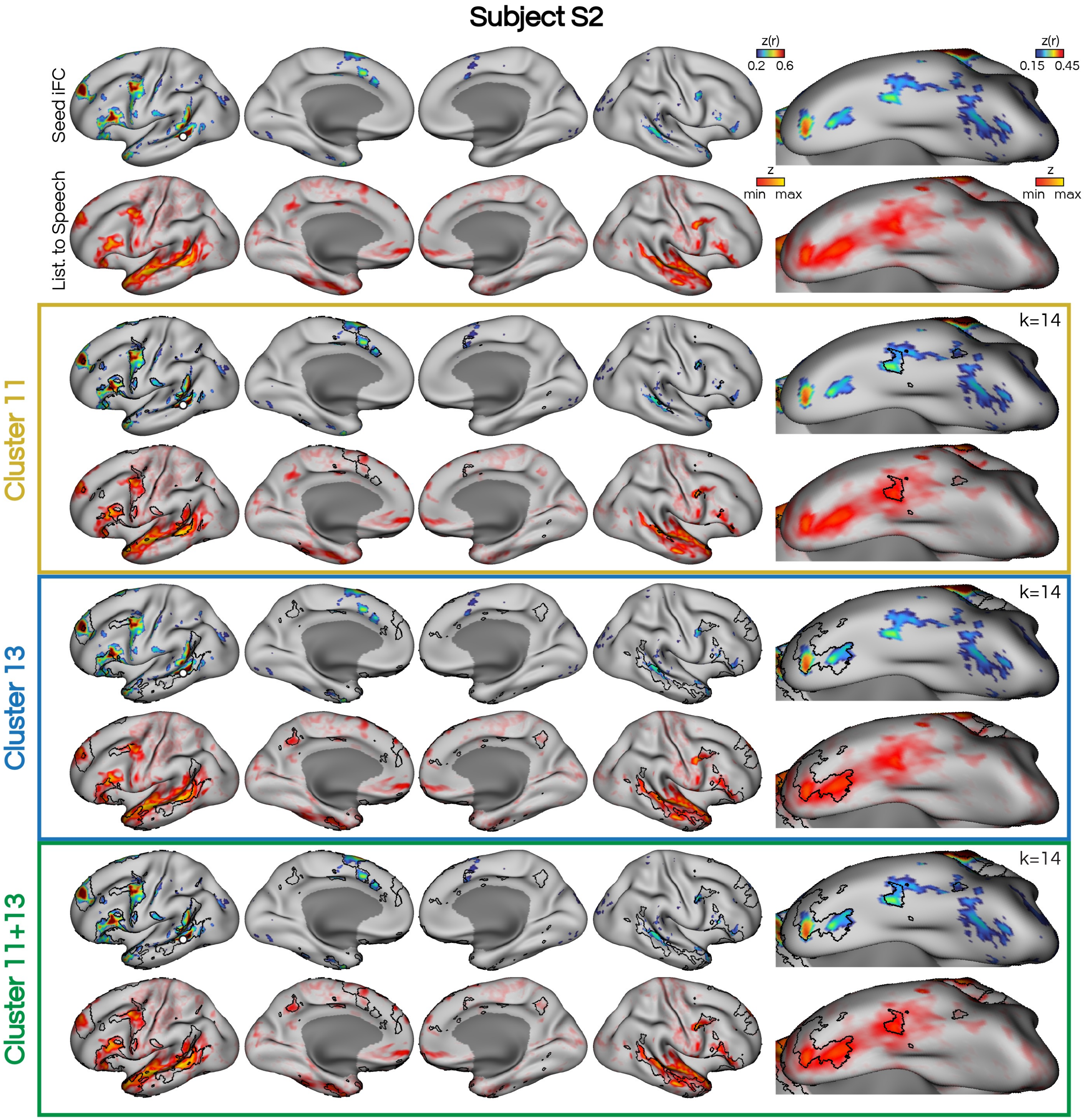


**Supp. Fig. S10: Post-hoc combination of clustering solutions for subject S2.** Following data-driven clustering, we decided to merge two clusters (11 and 13) from the k=14 clustering solution (shown as black borders) to respect that both clusters overlapped with the regions identified in the seed-based functional connectivity (Seed iFC) and task activity maps (List. to Speech).

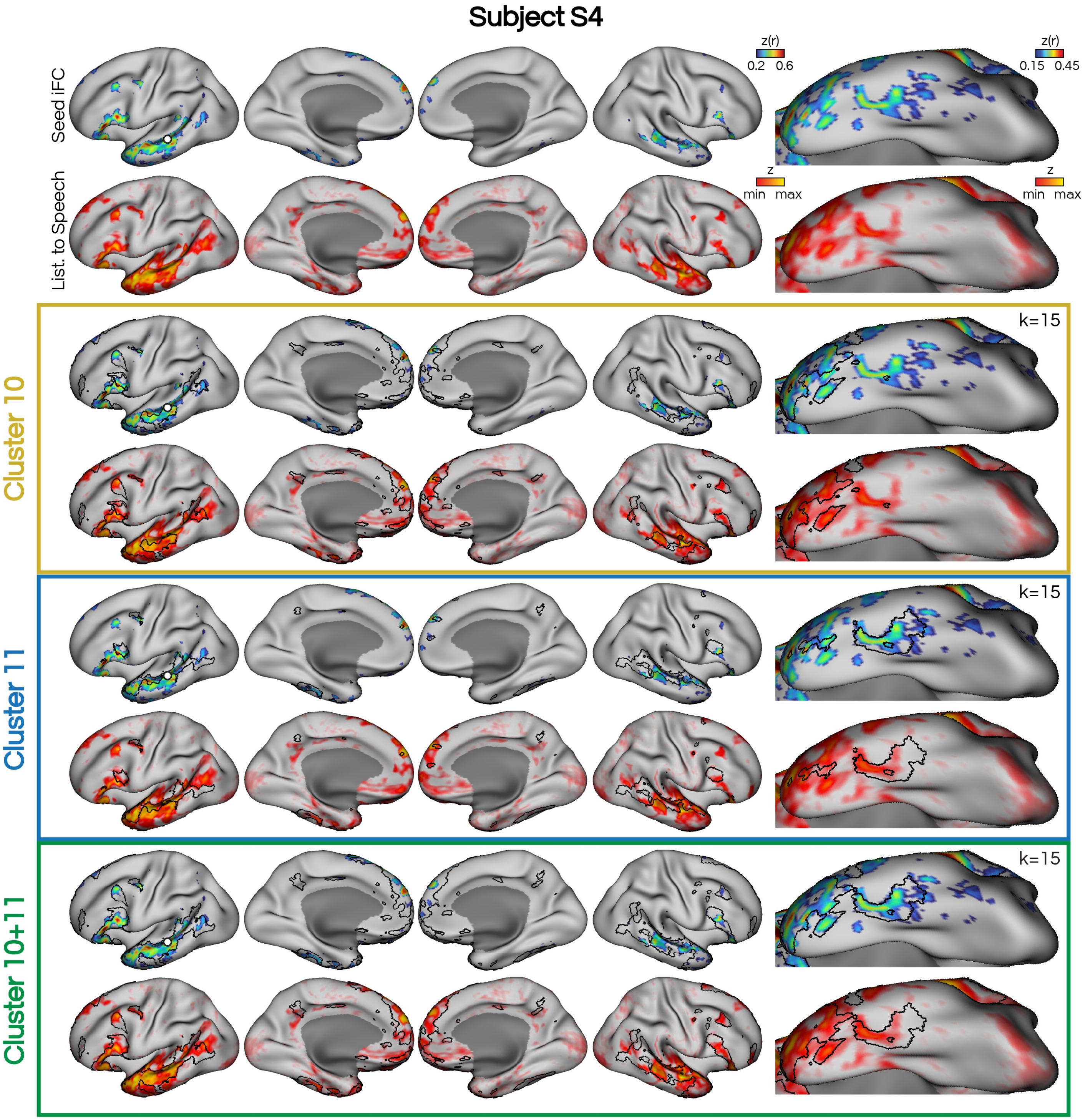


**Supp. Fig. S11: Post-hoc combination of clustering solutions for subject S4.** Following data-driven clustering, we decided to merge two clusters (10 and 11) from the k=15 clustering solution (shown as black borders) to respect that both clusters overlapped with the regions identified in the seed-based functional connectivity (Seed iFC) and task activity maps (List. to Speech).

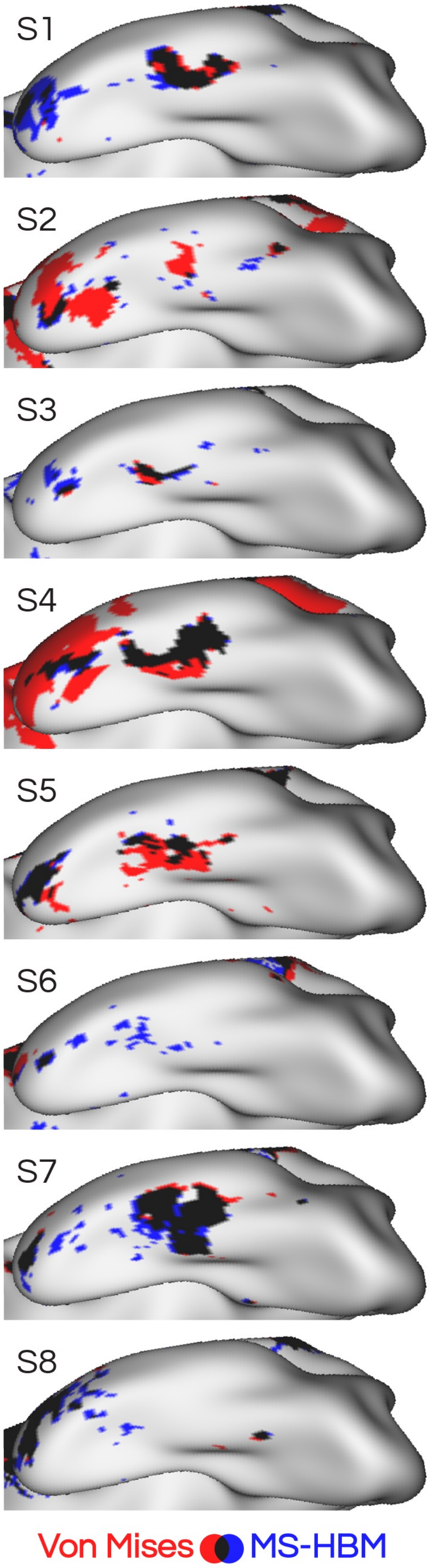


**Supp. Fig. S12: Comparison of clustering algorithm definitions of the LANG network.** Clusters derived using the Von Mises-Fisher algorithm (red) are overlaid with clusters derived using the Multi-Session Hierarchical Bayesian Model (MS-HBM; blue). Both approaches revealed an inferior region of the LANG network in the ITC in most subjects, however these varied. Shared regions are shown in black. The Von Mises-Fisher approach more often defined a larger ITC region of LANG, and was chosen as the network estimation approach for the main analyses.

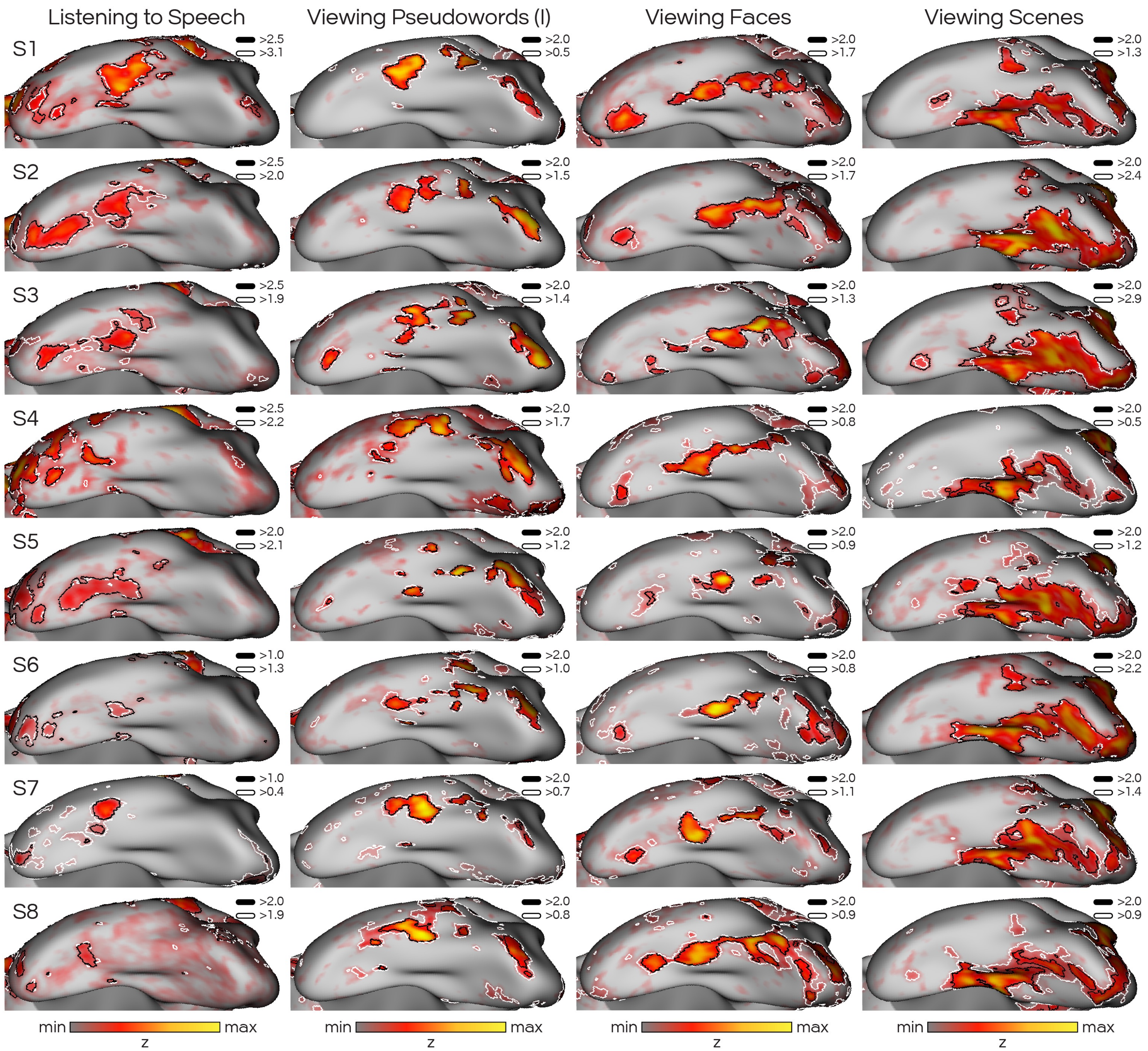


**Supp. Fig. S13: Regions showing activation in SPEECHLOC and VISCAT tasks were similarly delineated using qualitative (black borders; hand-selected) and quantitative (white borders; 90th percentile) thresholds.** Task activity is shown unthresholded using a heatmap color palette for the contrasts of Listening to Speech and Viewing Pseudowords ("I"; i.e., test), Faces, and Scenes. Both thresholds capture regions showing clear engagement (red-yellow) in each contrast.

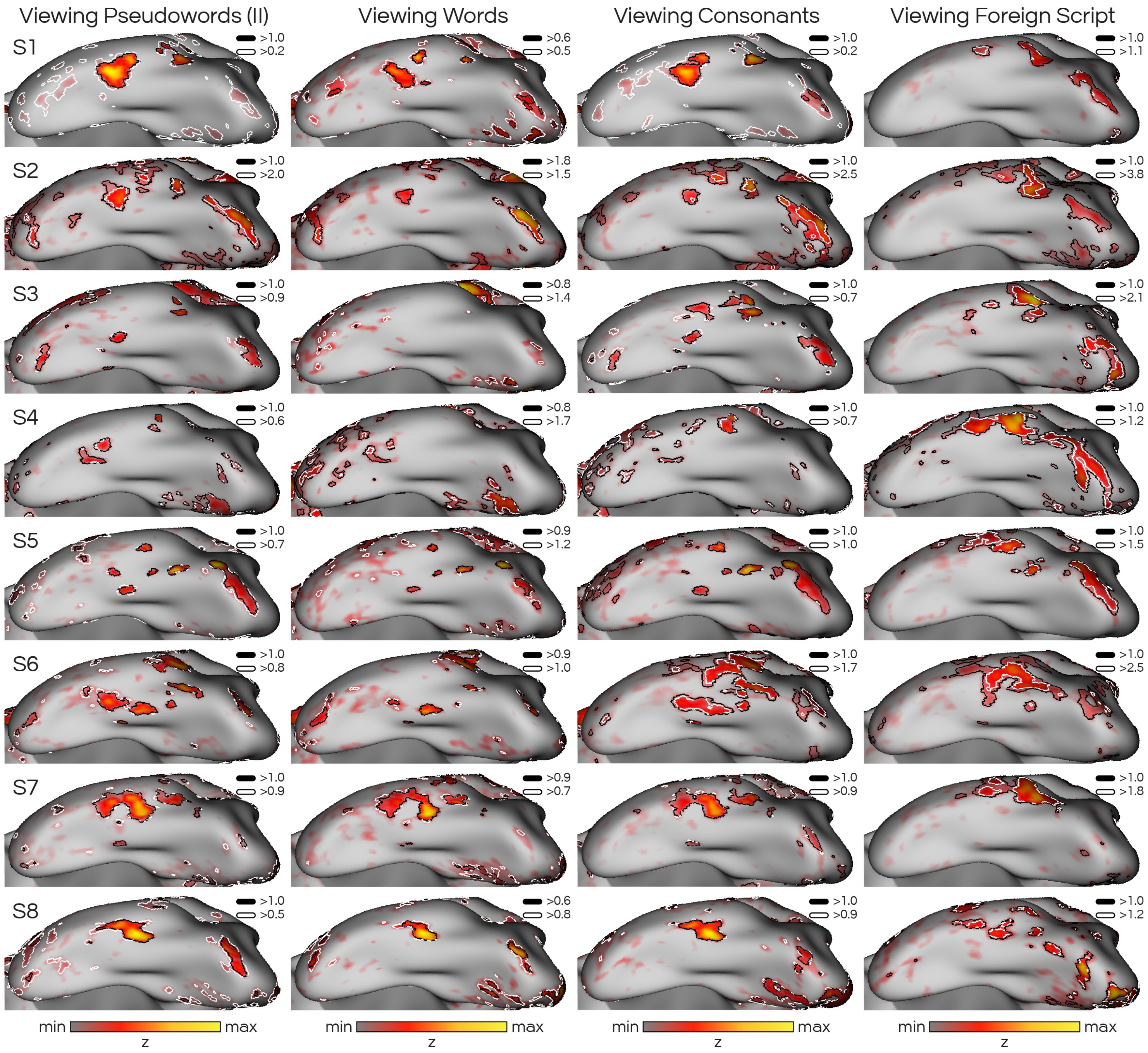


**Supp. Fig. S14: Regions showing task activation in the VWFACAT task were similarly delineated using qualitative (black borders; hand-selected) and quantitative (white borders; 90th percentile) thresholds.** Task activity is shown unthresholded using a heatmap color palette for the contrasts of Viewing Pseudowords (“II”; i.e., retest), Words, Consonants, and Foreign Script. Both thresholds capture regions showing clear engagement (red-yellow) in each contrast.

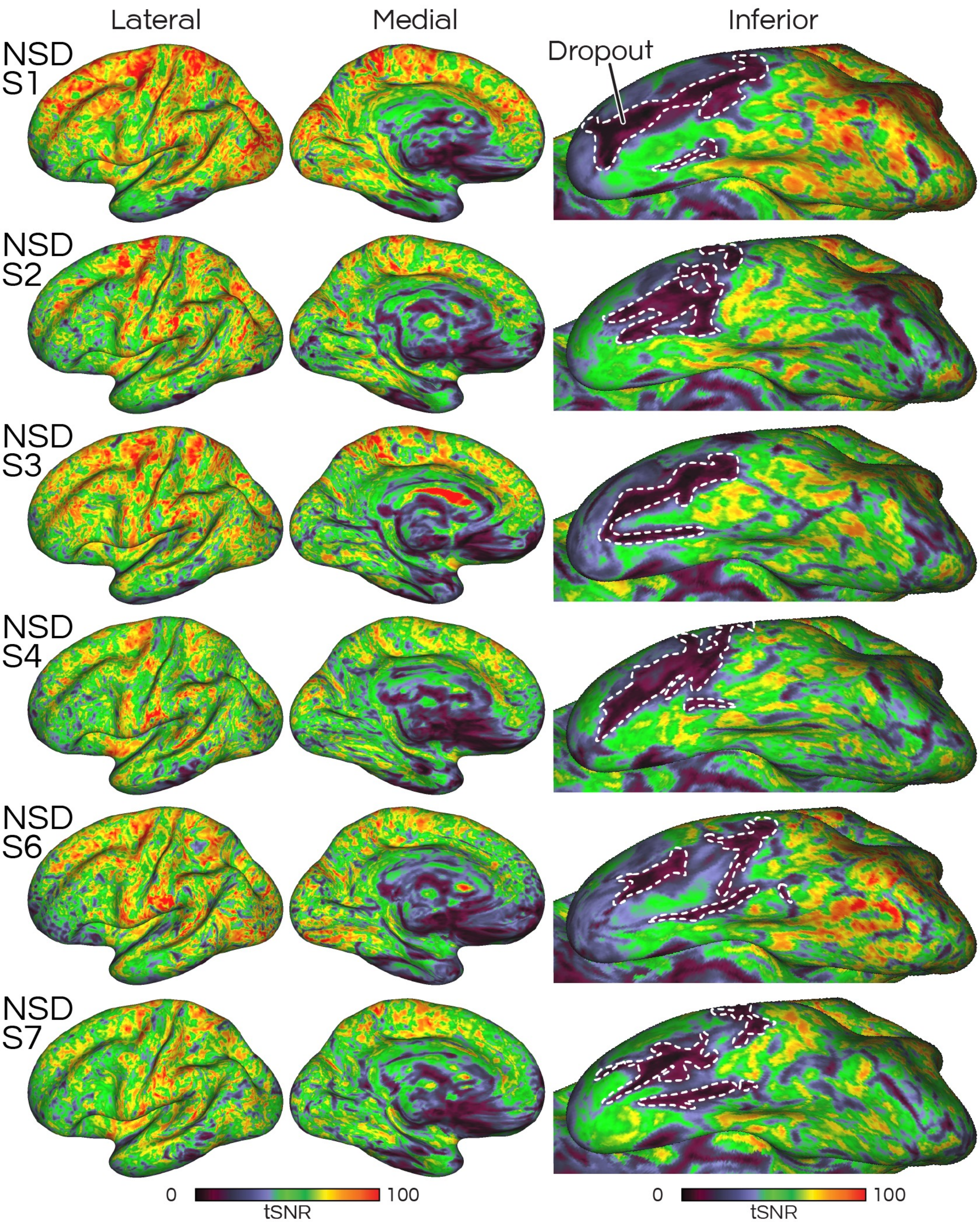


**Supp. Fig. S15: Temporal signal-to-noise ratio (tSNR) maps for each subject (NSD-S1 – NSD-S7) in the Natural Scenes Dataset (NSD).** High tSNR was achieved on the lateral surface, despite the smaller voxel size (1.8 mm isotropic), but with broader regions of signal dropout on the inferior temporal surface compared to the 3T multi-echo data (**Supp. Fig. S1**). White dashed borders have been hand drawn to indicate basal regions with the lowest tSNR values (~<20). Note that NSD-S5 and NSD-S8 were excluded during quality control of the resting-state data.

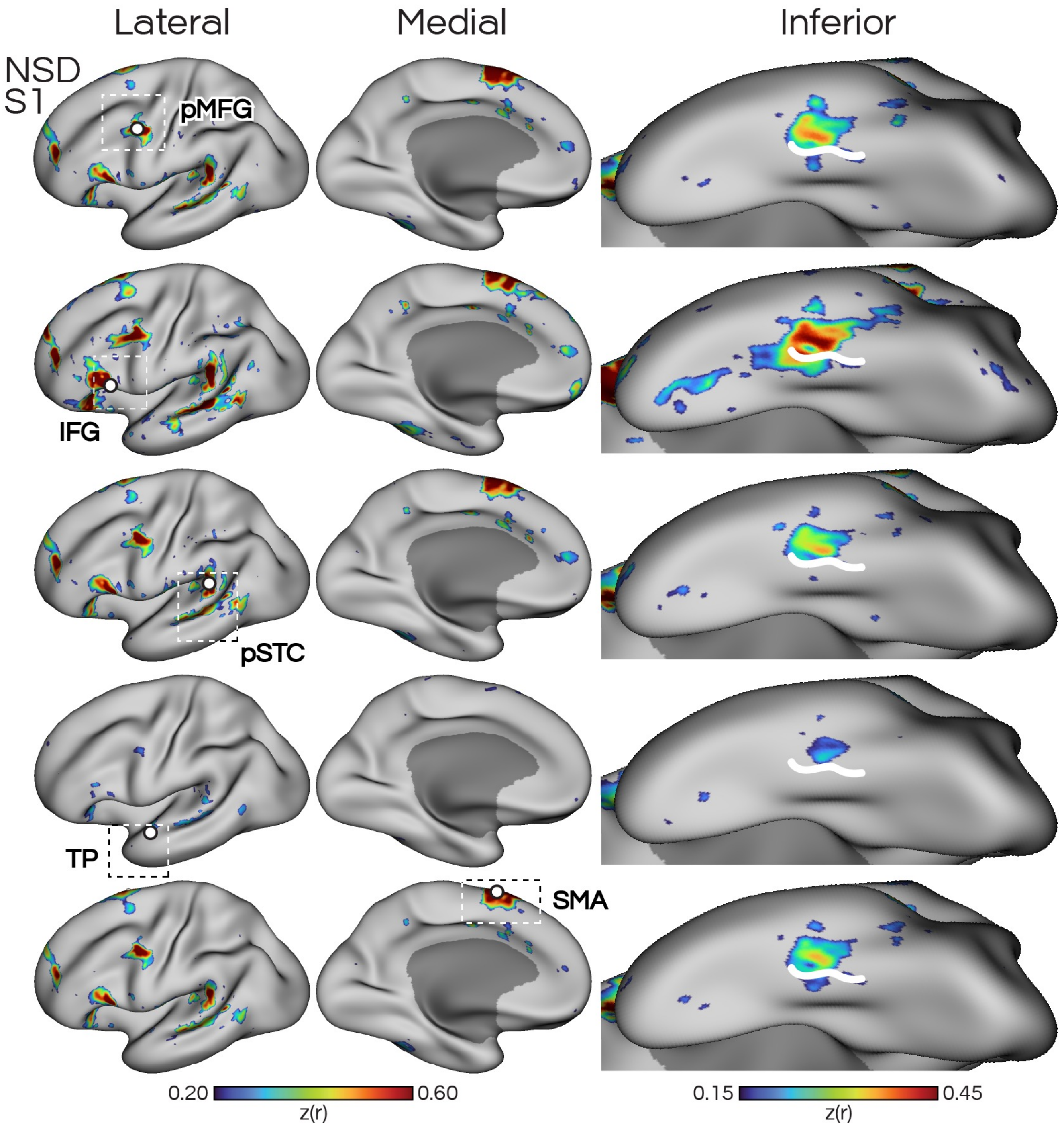


**Supp. Fig. S16: Replication of candidate language network in subject NSD-S1 using functional connectivity from seeds in five cortical zones.** The multiple estimates provided converging evidence that the distributed LANG network includes a region approximately halfway down the inferotemporal cortex. White lines are hand-drawn to serve as references and are consistent across panels. Maps depict Z-normalized Pearson correlations. Z thresholds were lowered for the inferior views to account for lower tSNR in these regions. pMFG: posterior middle frontal gyrus, IFG: inferior frontal gyrus, pSTC: posterior superior temporal cortex, TP: temporal pole, SMA: supplementary motor area.

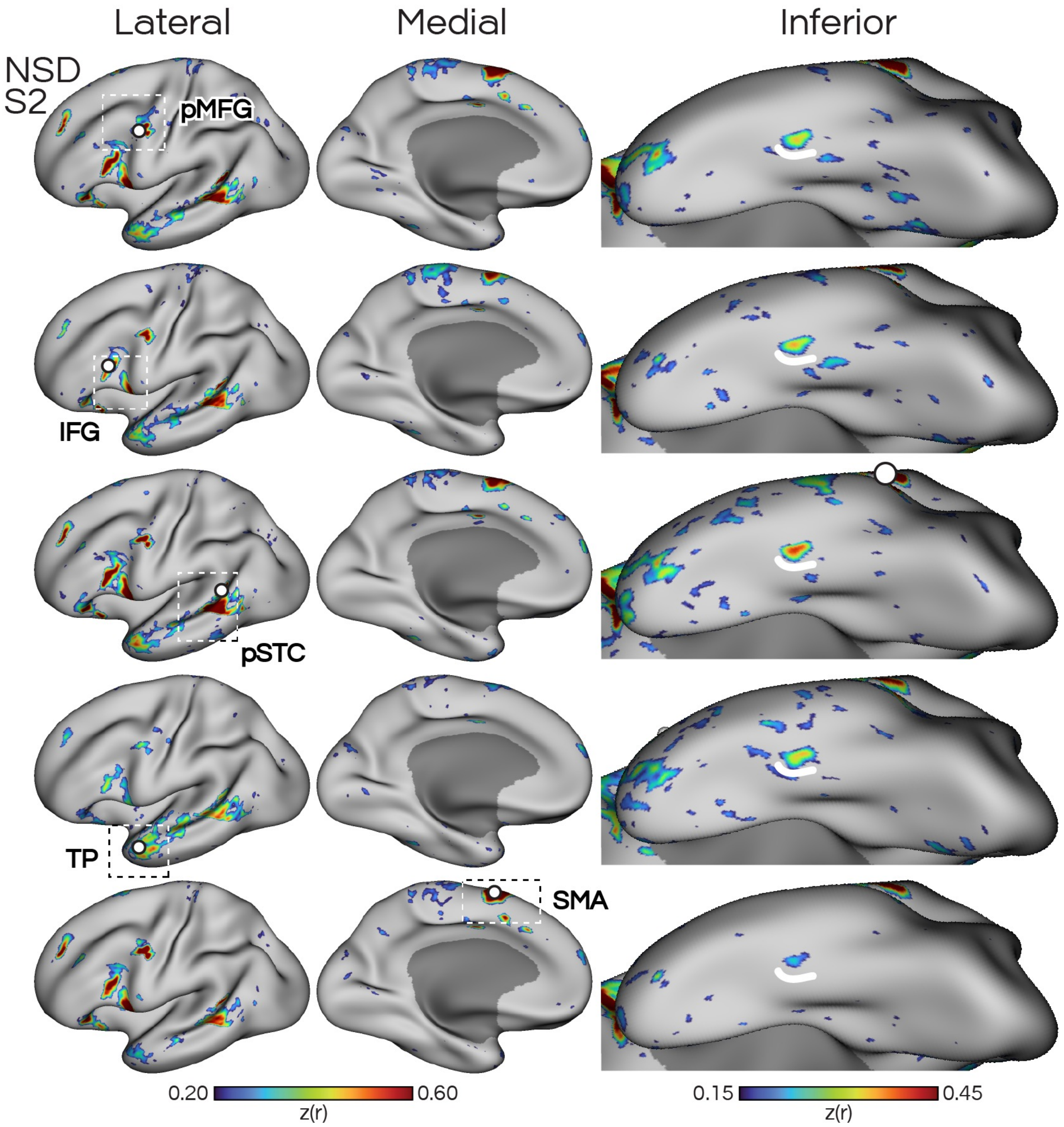


**Supp. Fig. S17: Replication of candidate language network in subject NSD-S2 using functional connectivity from seeds in five cortical zones.** Formatted according to **Supp. Fig. S16.**

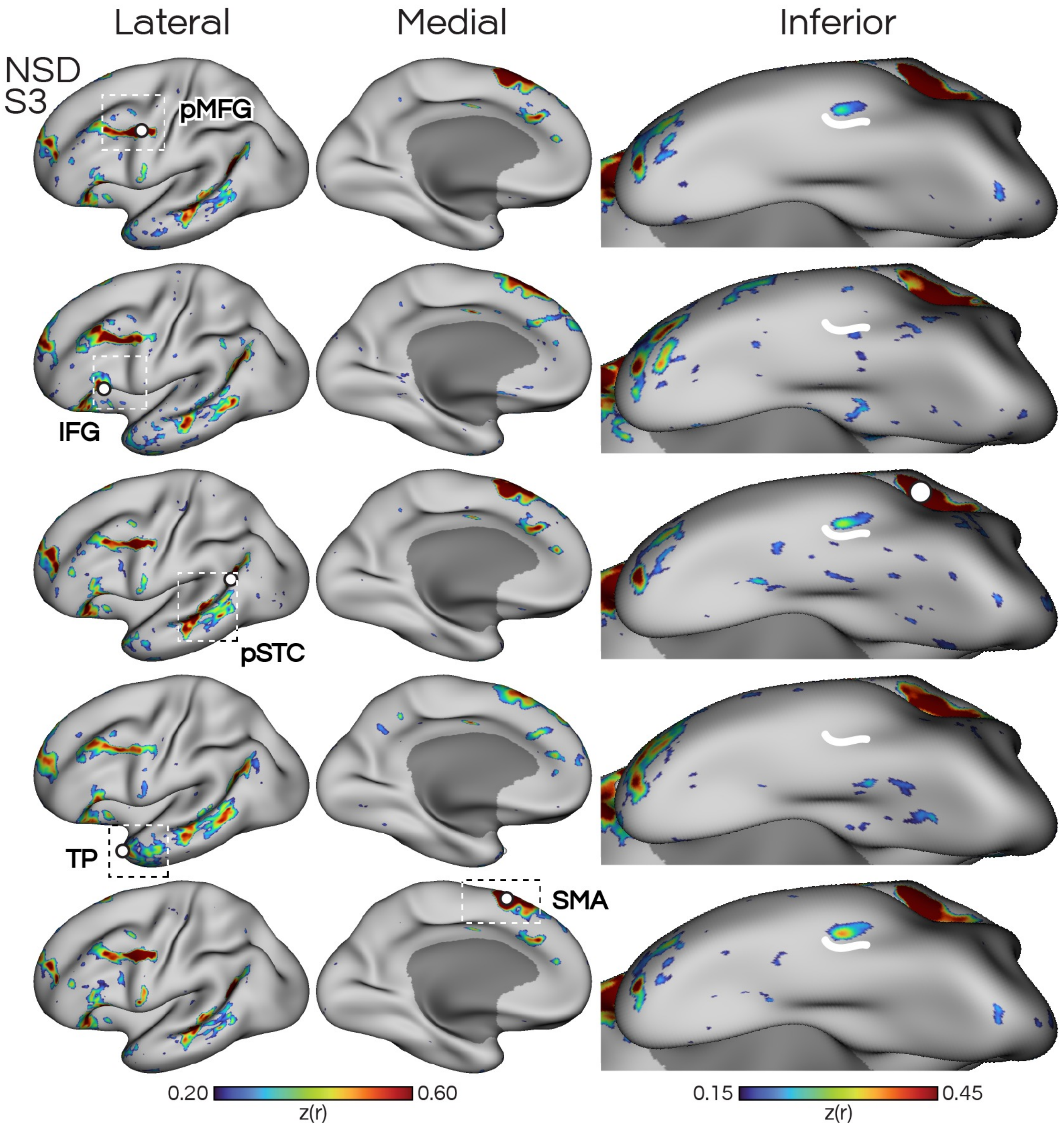


**Supp. Fig. S18: Replication of candidate language network in subject NSD-S3 using functional connectivity from seeds in five cortical zones.** Formatted according to **Supp. Fig. S16.**

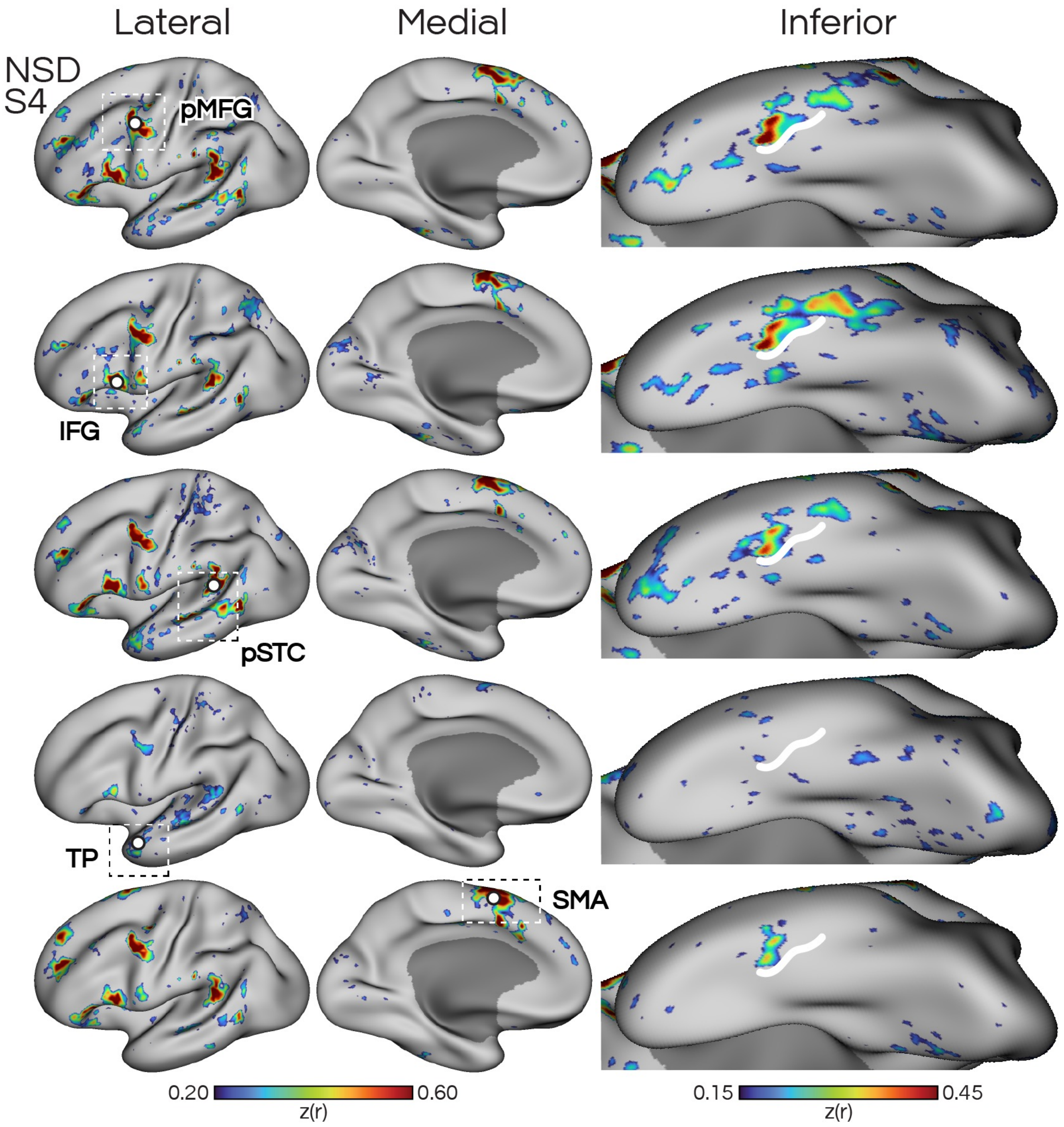


**Supp. Fig. S19: Replication of candidate language network in subject NSD-S4 using functional connectivity from seeds in five cortical zones.** Formatted according to **Supp. Fig. S16.**

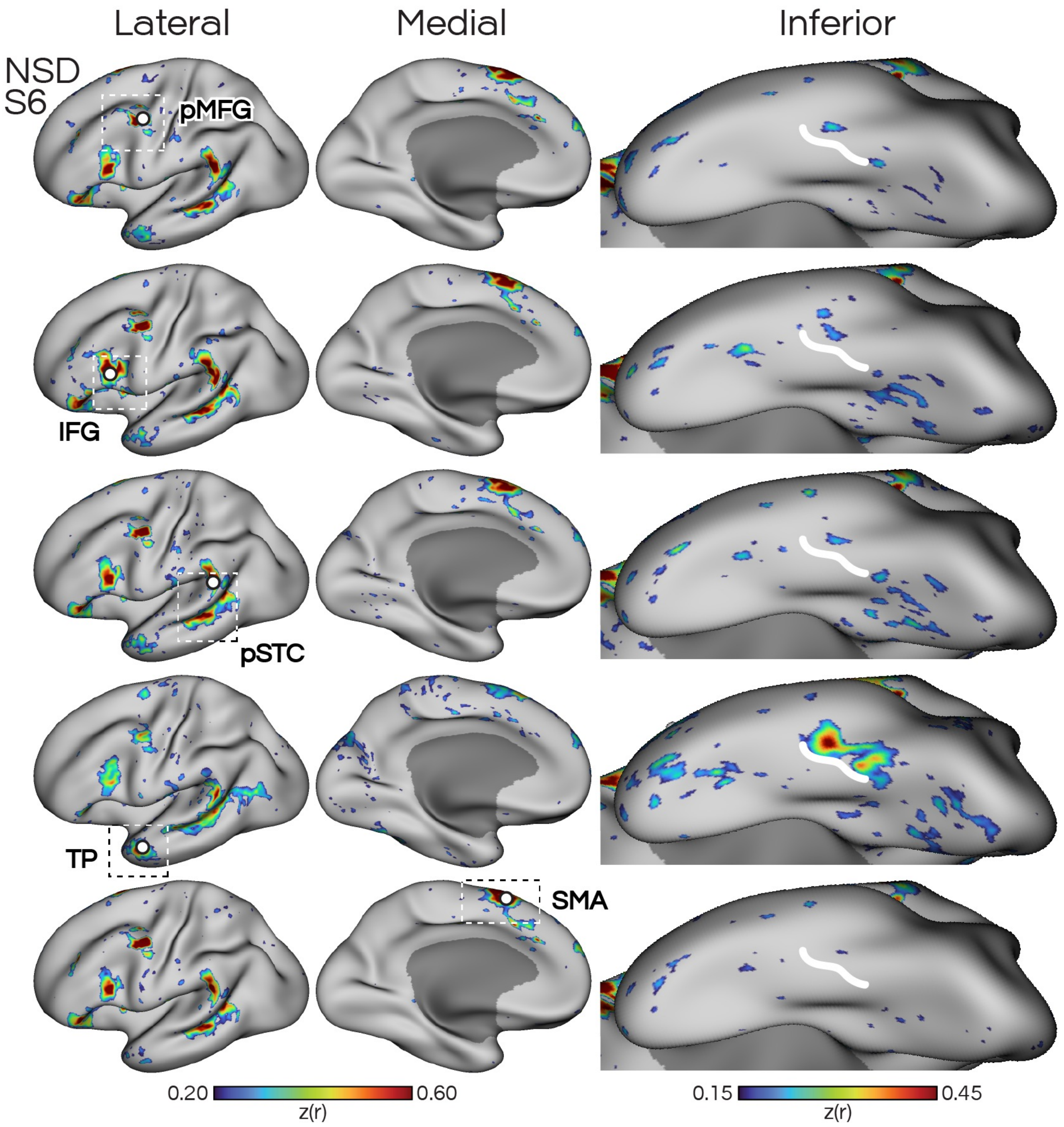


**Supp. Fig. S20: Replication of candidate language network in subject NSD-S6 using functional connectivity from seeds in five cortical zones.** Formatted according to **Supp. Fig. S16.**

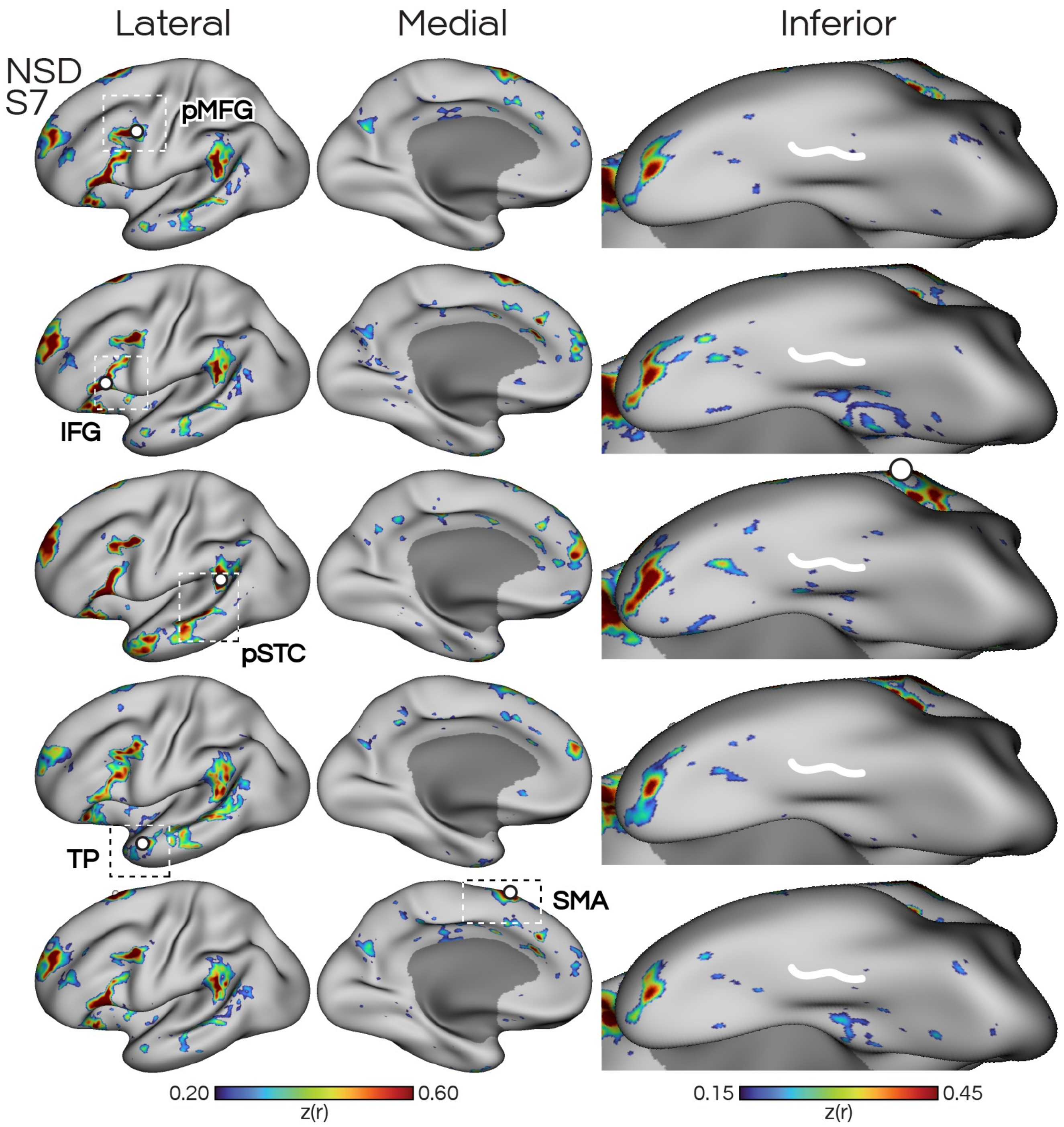


**Supp. Fig. S21: Replication of candidate language network in subject NSD-S7 using functional connectivity from seeds in five cortical zones.** Formatted according to **Supp. Fig. S16.** Limited/no evidence of an inferior temporal LANG network region was seen in this subject, potentially due to broader signal dropout (shown in **Supp. Fig. S15**)

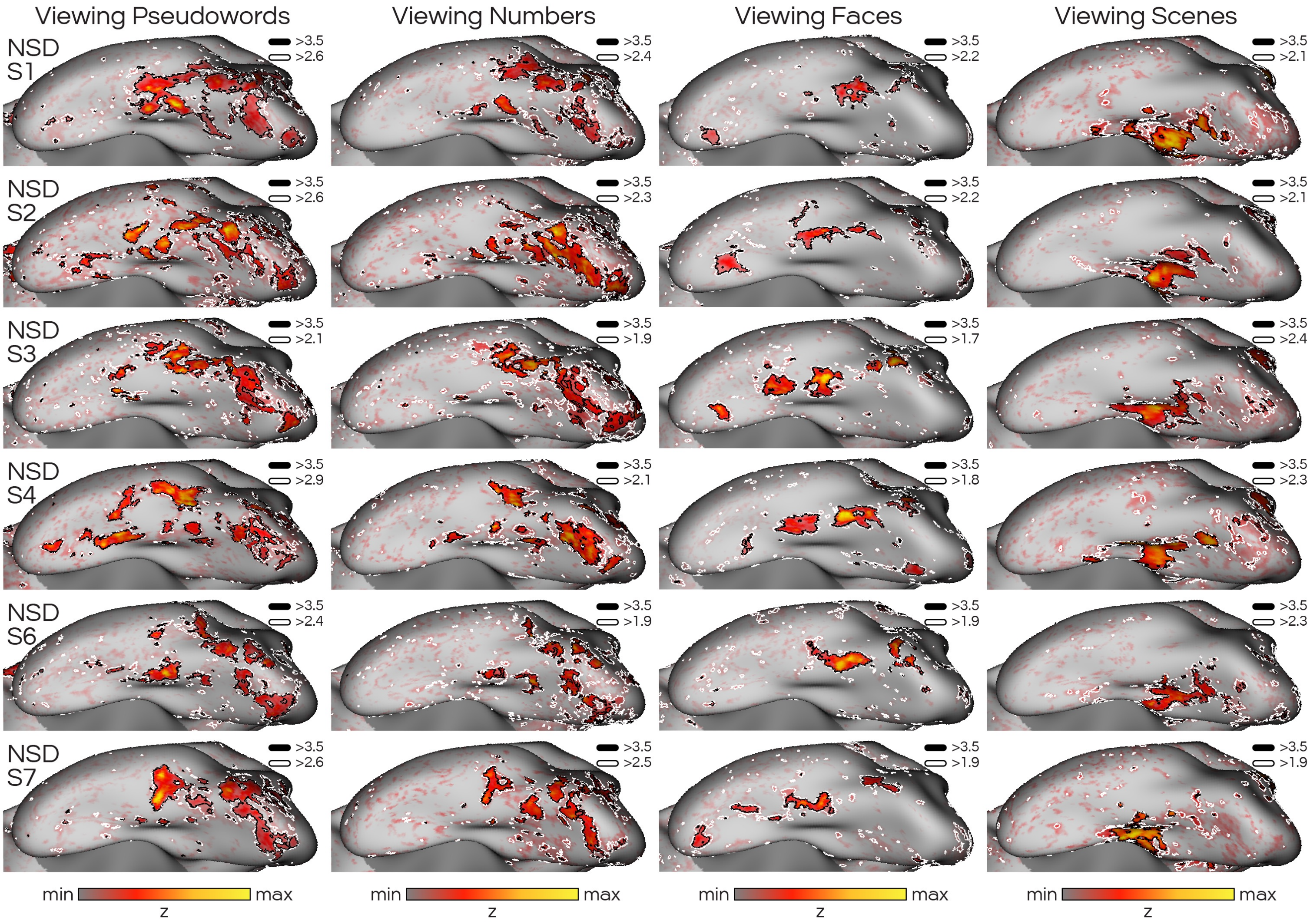


**Supp. Fig. S22: Regions showing activation in NSD fLoc tasks were similarly delineated using qualitative (black borders; hand-selected) and quantitative (white borders; 95th percentile) thresholds.** Task activity is shown using a heatmap color palette, and network boundaries are shown as black borders. Hand-selected) Display thresholds were chosen qualitatively per participant, to minimize noise-related "speckling", and to account for signal quality variations between different individuals. 95th Percentile) Thresholds were also chosen quantitatively by only including vertices within the 95th percentile. The maps look similar using both methods.

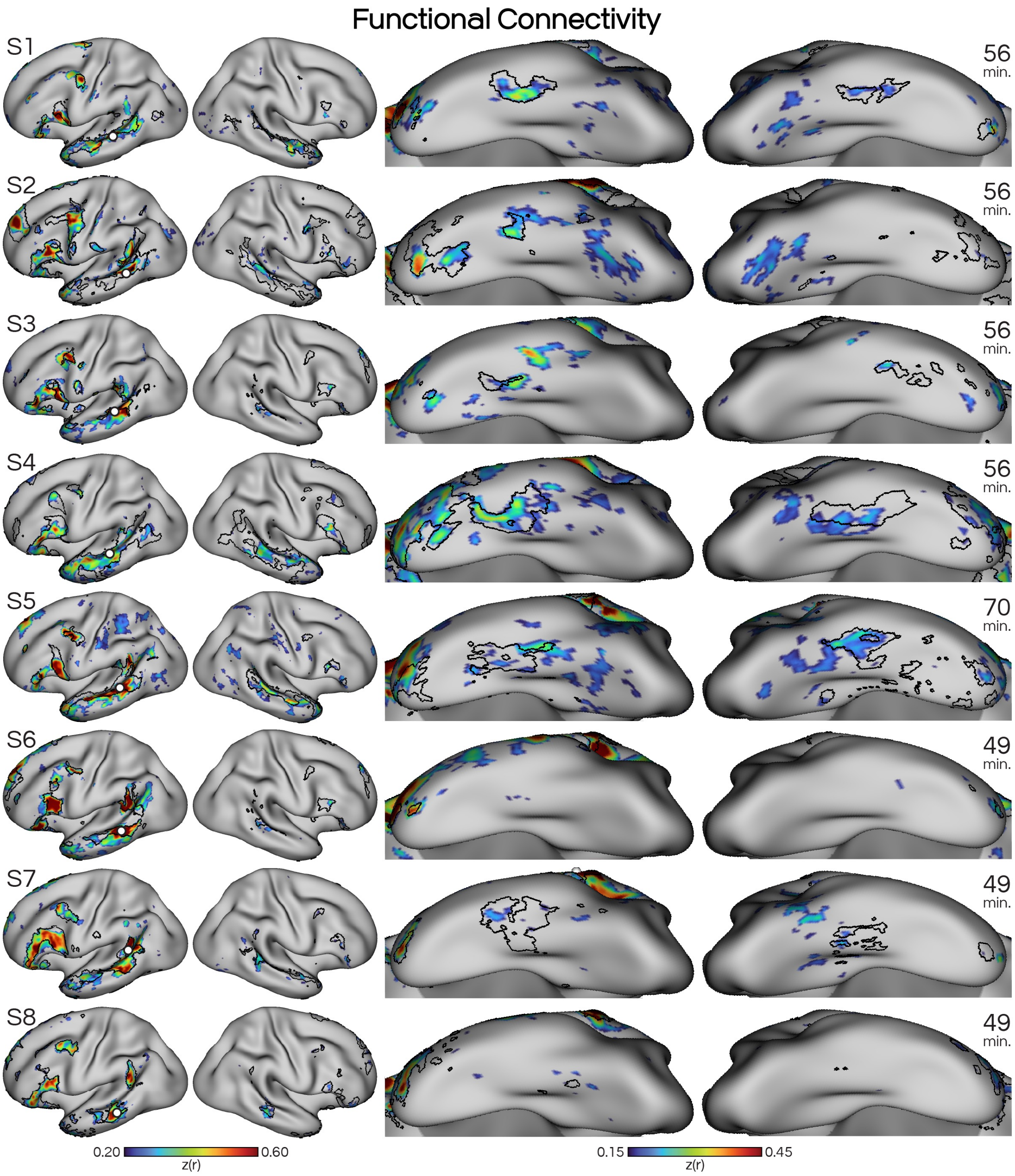


**Supp. Fig. S23: Seed-based and clustering estimates both support that the language network has larger regions on the left hemisphere.** Regions showing the greatest correlations (Turbo colormap) with the selected seed (white circle) aligned with the data-driven clustering estimate (black borders), both on the lateral and inferior surfaces. Total times of included runs for each participant are on the far right.

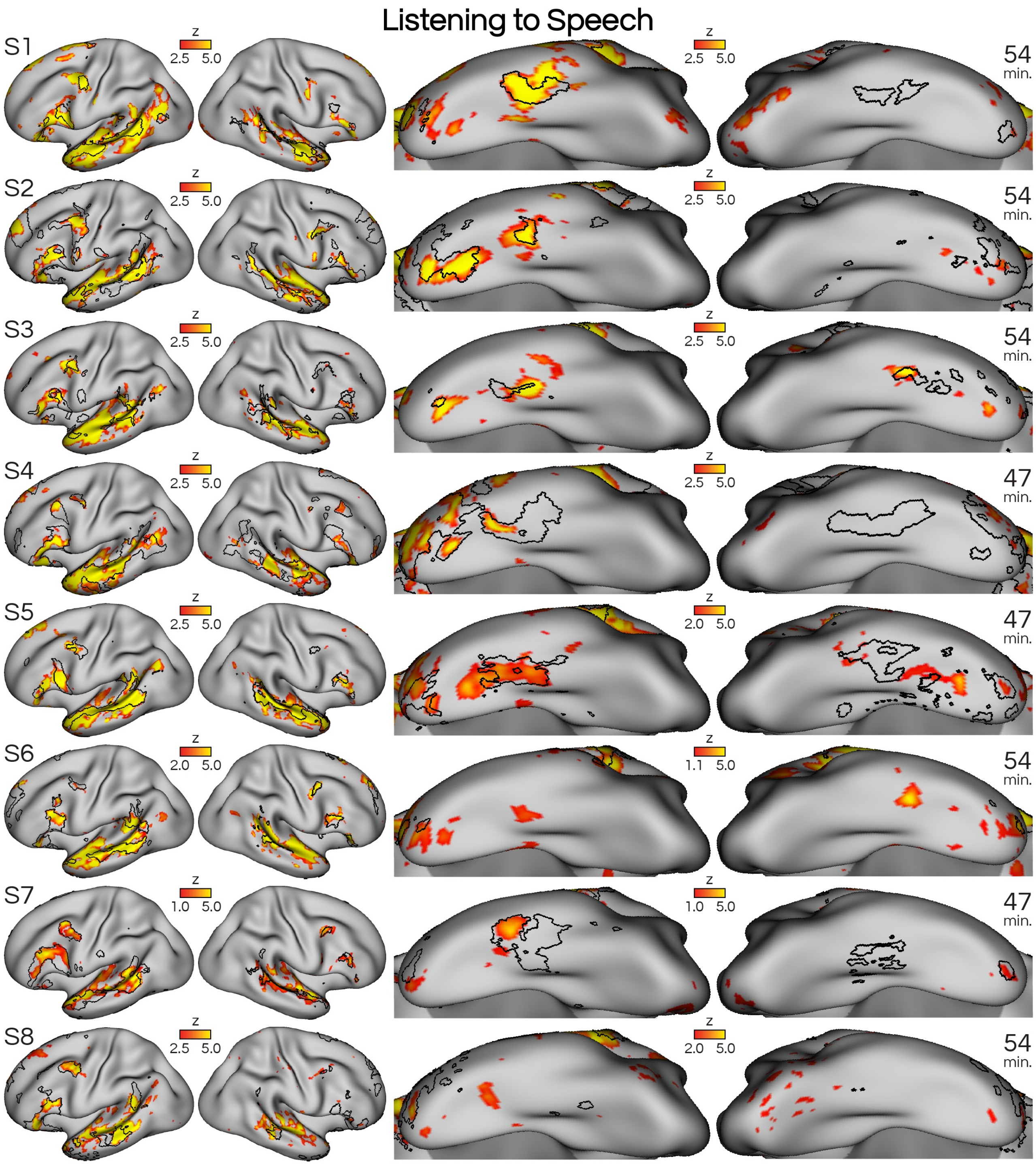


**Supp. Fig. S24: Listening to speech (contrasted with listening to distorted speech; SPEECHLOC task) activated the distributed LANG network including basal LANG region.** Regions showing greater Z-normalized beta weights (heatmaps) were generally within network regions as defined using iFC through data-driven clustering (black borders), including on the inferior surface. Speech-related activity was generally more extensive and robust on the left hemisphere, implying left lateralization. Total times of included runs for each participant are on the far right.

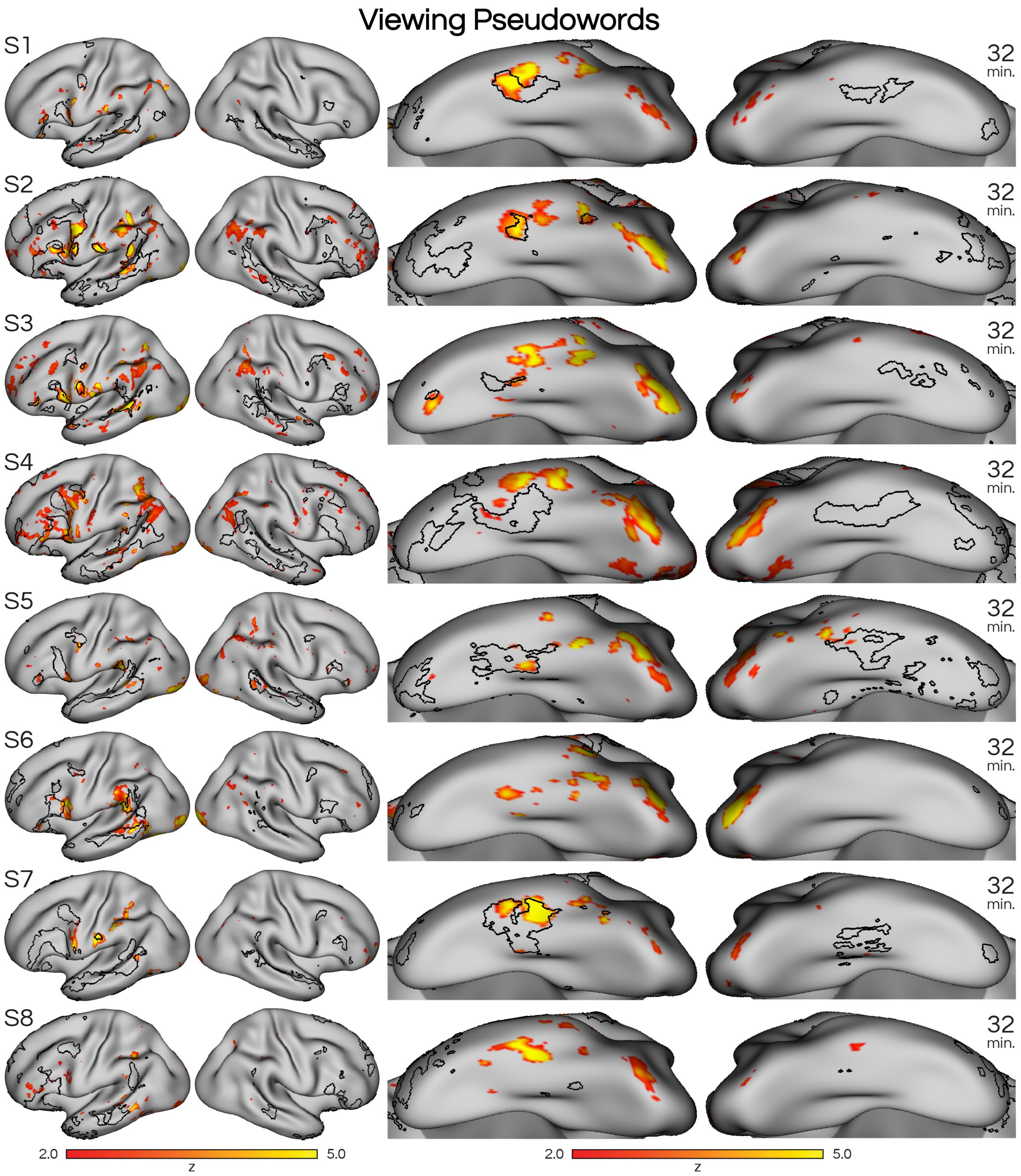


**Supp. Fig. S25: Viewing written pseudowords (contrasted with viewing faces, scenes, objects; VISCAT task) activated a stream of posterior inferotemporal regions, with the most anterior region overlapping with the LANG network at the basal LANG area.** Heatmaps shows Z-normalized beta weights. In contrast to the language localizer maps, activation was more robust on the inferior surface than on the lateral surface. Total times of included runs for each participant are on the far right.

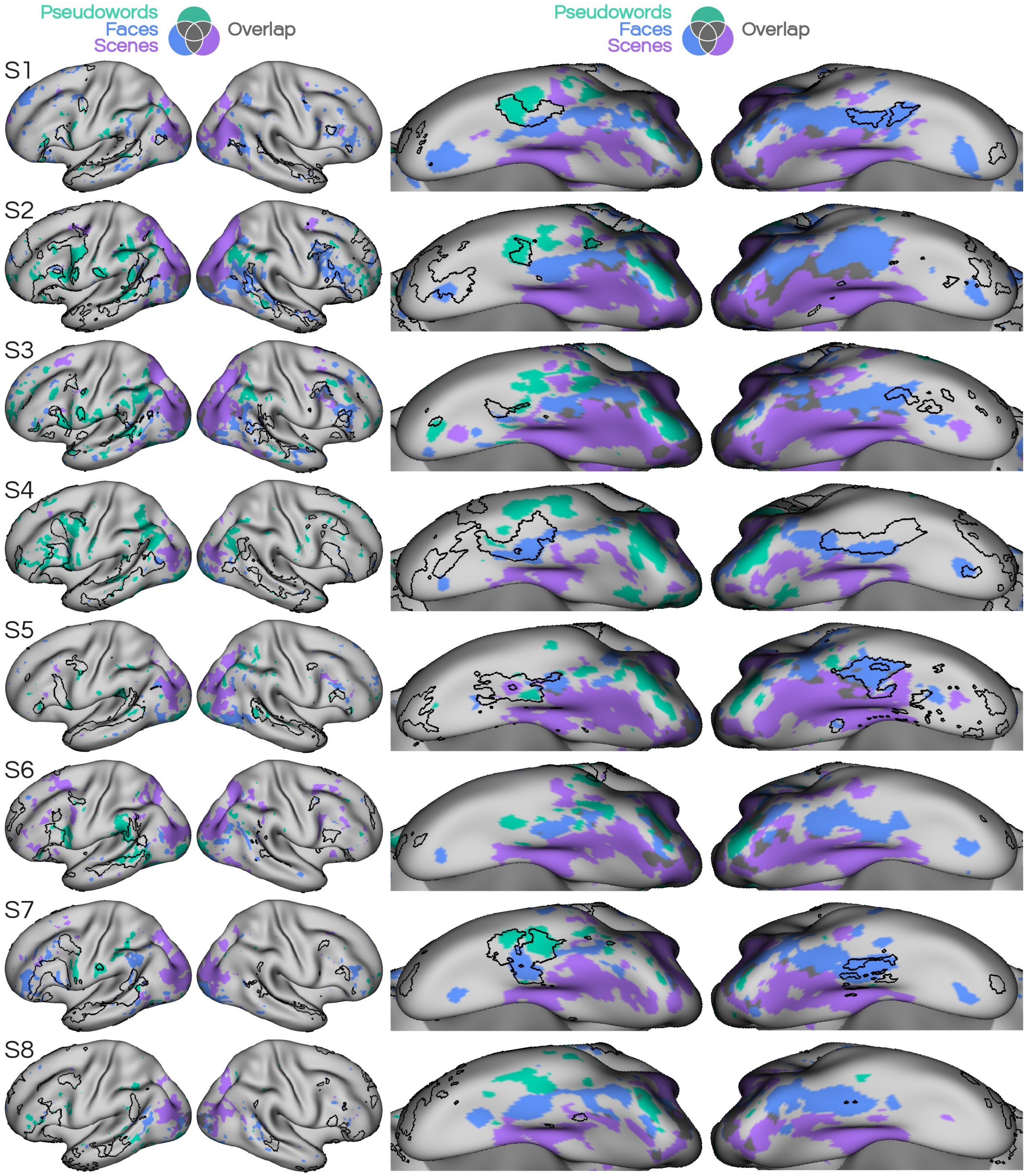


**Supp. Fig. 26: Activity evoked by viewing pseudowords is left-lateralized, while activity for faces or scenes is more bilateral.** Binarized activity from the pseudowords (teal) faces (blue) and scenes (purple) conditions displayed on left and right hemispheres. Pseudoword regions are larger and more prevalent on the left hemisphere, while face and scene regions appear more symmetrically distributed.

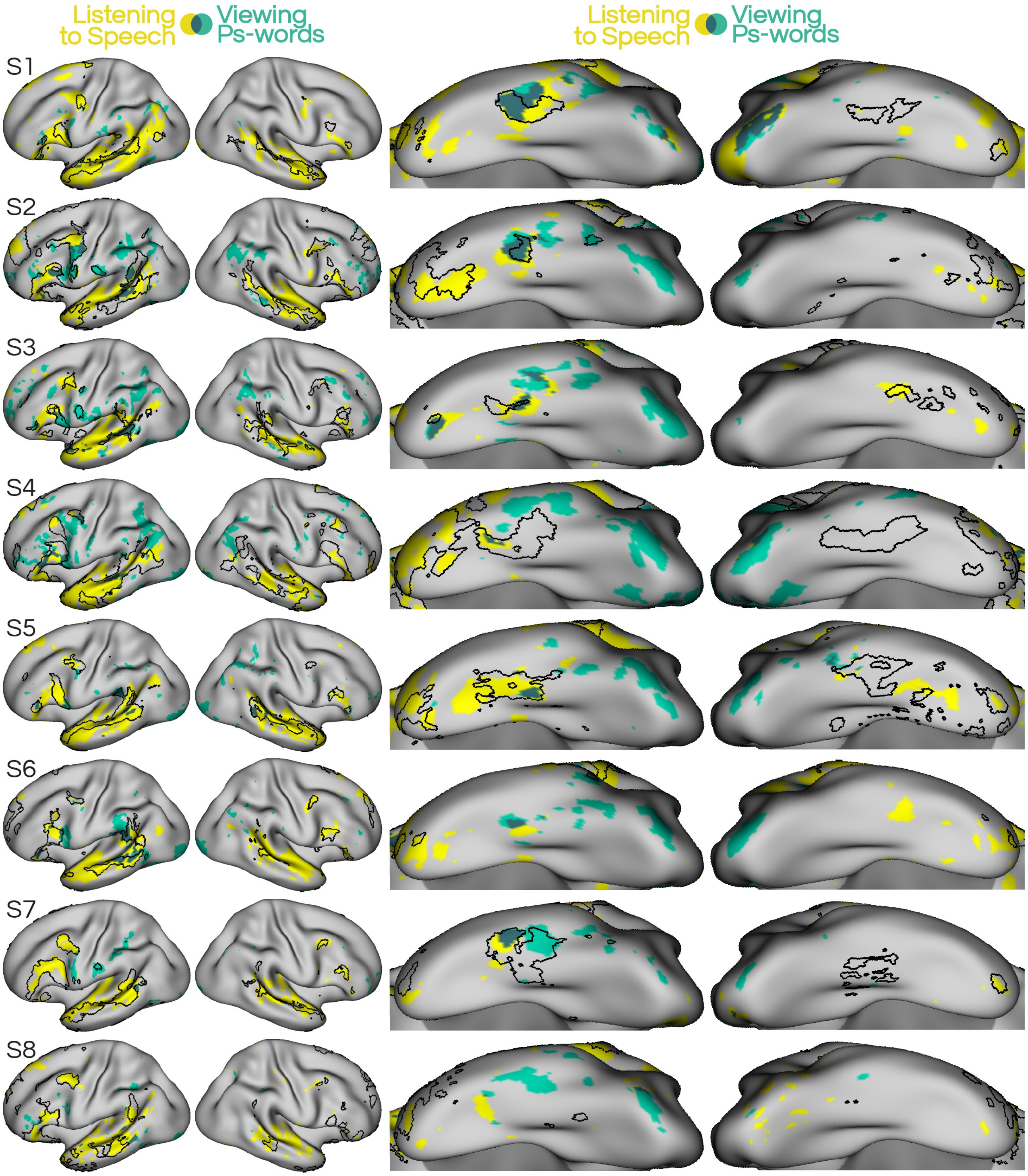


**Supp. Fig. 27: Pseudowords and speech activity lateralize to the same side.** Binarized activity from the speech (yellow) and pseudowords (teal) conditions displayed on left and right hemispheres. Pseudoword regions are larger and more prevalent on the left hemisphere, where they more often align with the basal LANG region (black borders).

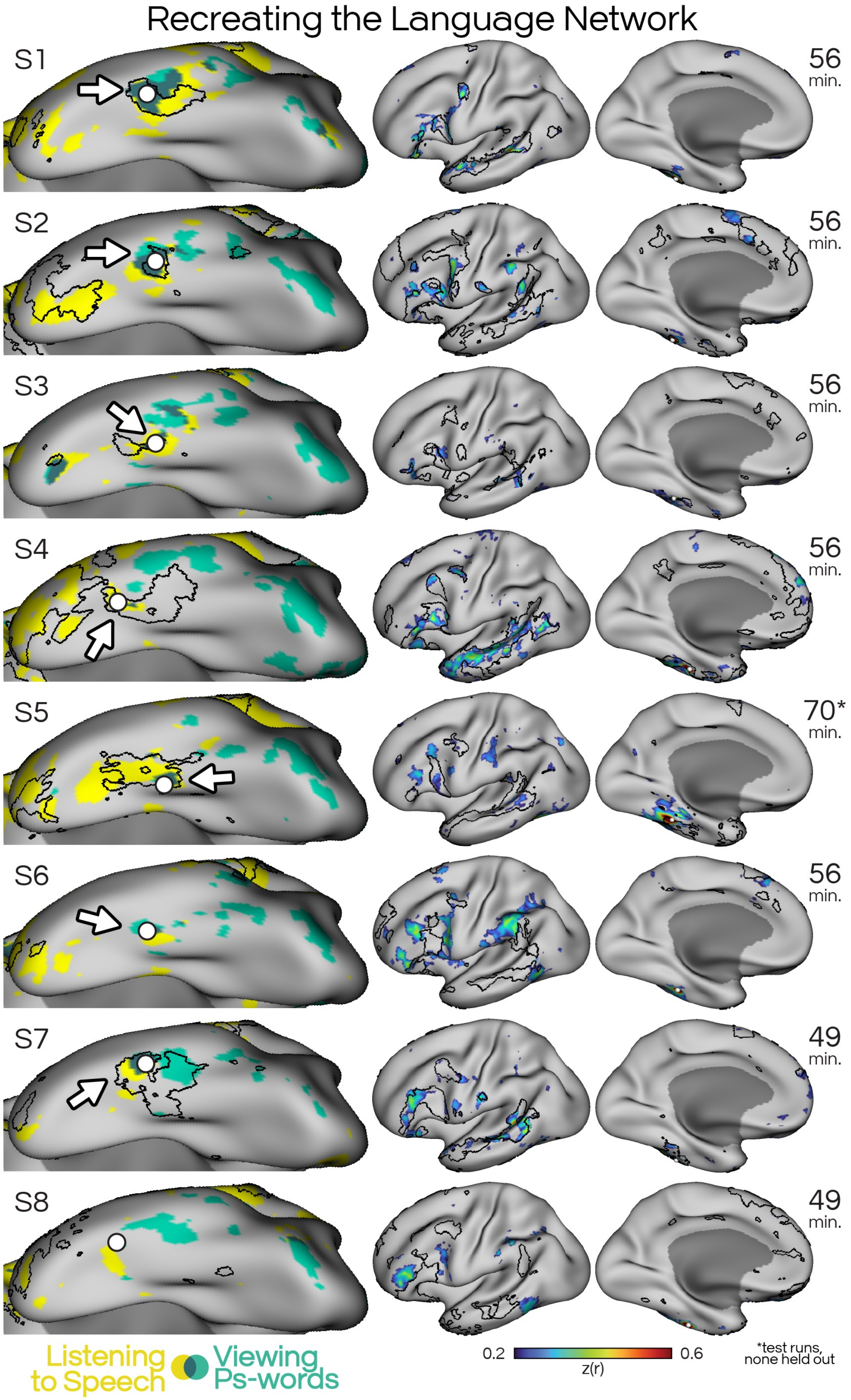


**Supp. Fig. S28: Seeds chosen in regions selective for both listening to speech and viewing pseudowords recreated the LANG network.** *Left:* For each participant, a seed (white circle) was chosen in a mid-ITC region demonstrating activity for both spoken language and pseudoword images (dark green). *Right:* Resulting seed correlation pattern (rainbow) in held-out resting-state runs is displayed relative to the language cluster boundaries estimated using the original resting-state runs (white borders). Total time of held-out rest runs for each participant are on the far right. The correlation patterns were broadly aligned to the LANG network boundaries, though the degree of correspondence varied across participants. Subjects S6 and S8 displayed maps which did not recreate the LANG network, in keeping with their results in the main analyses which failed to define a basal LANG region using iFC.

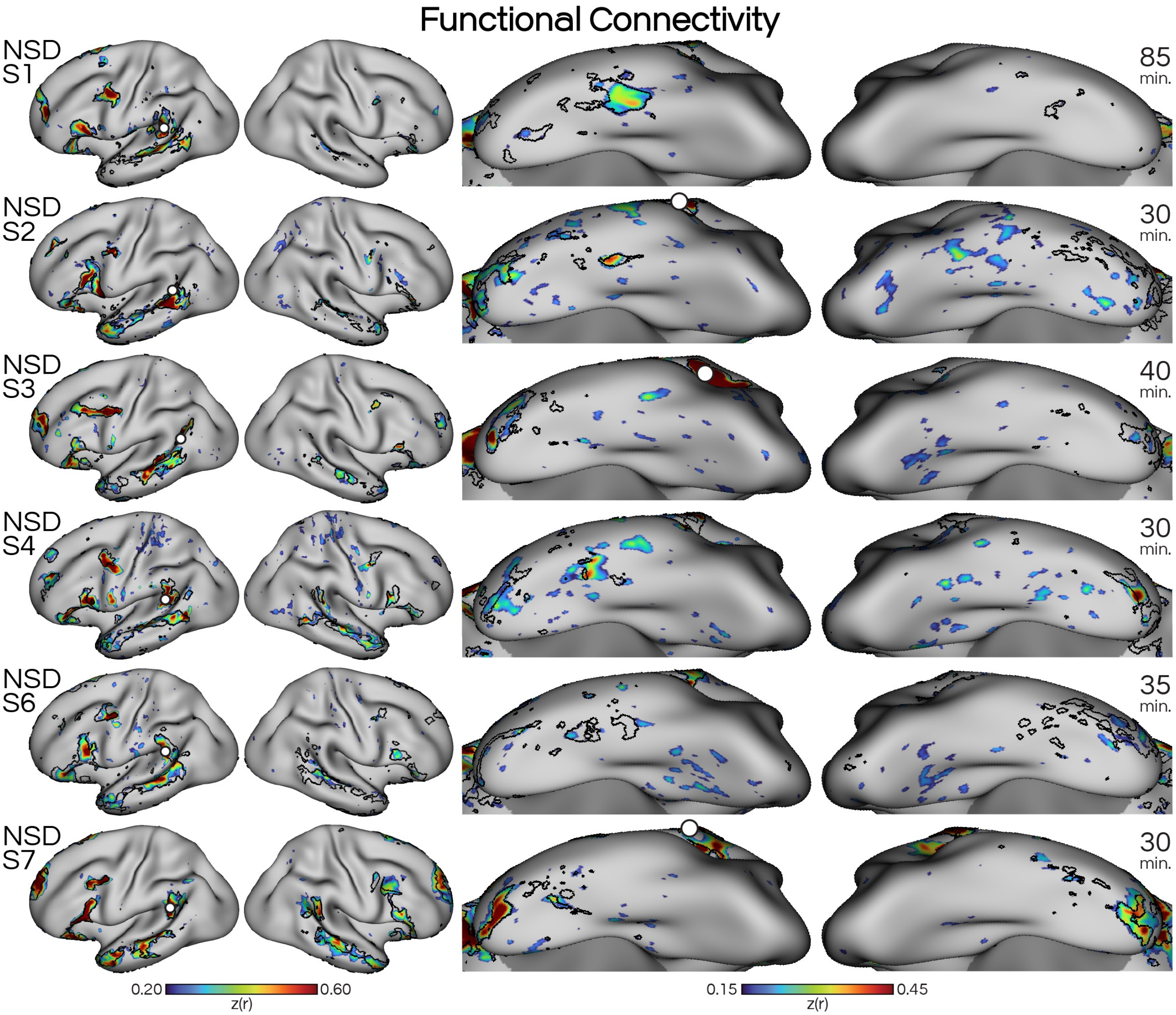


**Supp. Fig. S29: Functional connectivity estimates of the LANG network in participants from the Natural Scenes Dataset using converging approaches.** Seed-based correlation maps are shown in the Turbo colorbar, and data-driven clustering estimates are shown as black borders. Total times of included runs for each participant are on the far right.

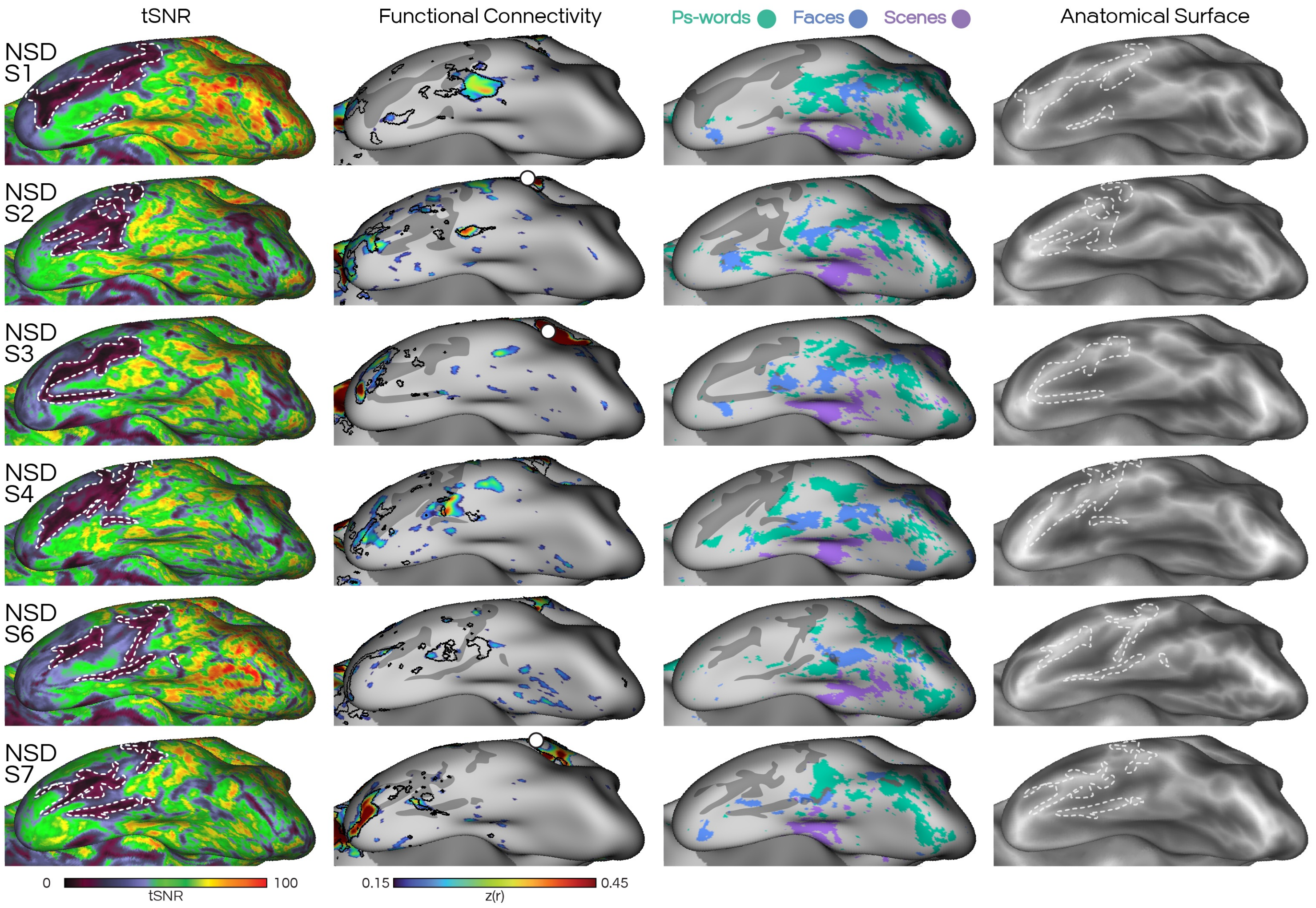


**Supp. Fig. S30: Gyral anatomy aligns with regions showing signal dropout in the Natural Scenes Dataset.** *Left-most column:* Temporal signal to noise ratio (tSNR) maps reveal regions of particularly low signal, indicating dropout (see hand-drawn white borders on left and right columns and dark gray regions in other columns). These dropout regions aligned with participant-specific gyral anatomy, and likely explain instances in which functional connectivity and task based analyses failed to identify regions in the NSD subjects. *Middle left column:* Intrinsic functional connectivity maps of the LANG network estimate using seed-based correlations (Turbo colormap) and data-driven clustering (black borders). Note how the basal LANG region in subject NSD-S6, is split in half by a region if signal dropout. *Middle right column:* Regions showing task responses to visual stimuli including pseudoword (teal), face (blue), and scene (purple) conditions are also affected by signal dropout. *Right-most column:* Overlaying the dropout maps onto each subject's sulcal depth map (grayscale) indicates that dropout regions were predominantly on the crowns of the gyri (lighter gray).

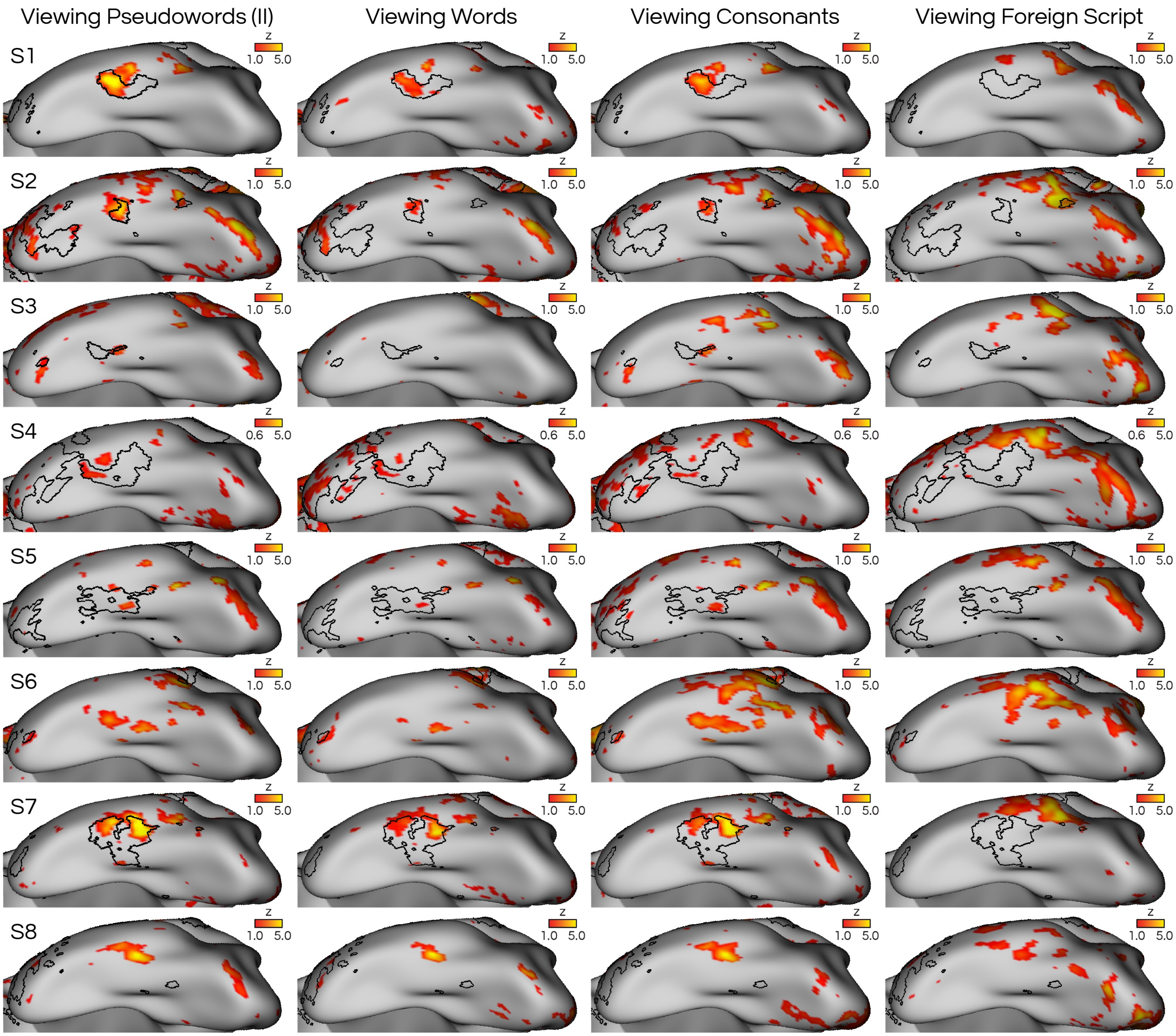


**Supp. Fig. S31: Task contrast maps for visual stimuli with decreasing resemblance to written words from the VWFACAT task.** Heatmaps display Z-normalized beta values for (from left to right) viewing pseudowords, real words, consonant strings, and foreign script, all contrasted against viewing line drawings. In most participants, a mid-ITC region that overlapped with the LANG network (black borders) was active for all conditions except the foreign script, supporting that the basal LANG region responds to familiar character strings (in this case, Roman characters) but not unfamiliar characters (in this case, modified Japanese characters).

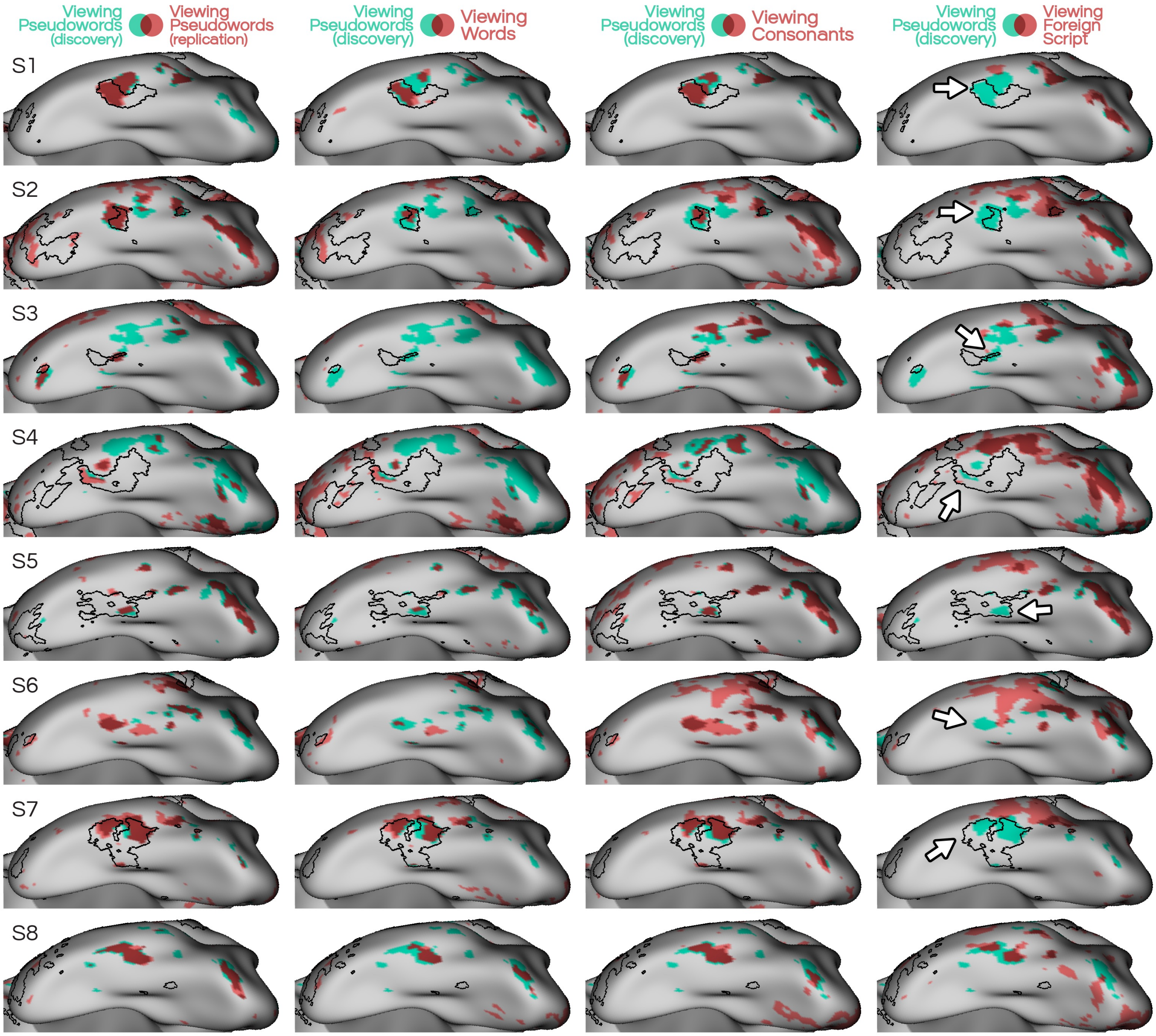


**Supp. Fig. S32: The orthographic stream is activated for familiar character strings (including real words, pseudowords, and consonant strings), but not unfamiliar foreign characters.** The orthographic stream defined from the VISCAT task (**Figs. 3 & 4**) is shown in teal in all panels, with red showing the regions active in each contrast from the VWFACAT task, along with overlap in darker red. *Left-most column:* The contrast of pseudowords > line drawings replicated that the orthographic stream responds to pseudowords in most participants, with the possible exception of subjects S2 and S3. *Middle columns:* The same stream of regions also demonstrated greater activity for real words and consonant strings compared to line drawings. *Right-most column:* In most participants, viewing foreign characters (modified Japanese) that none of the participants reported being able to read activated the more posterior regions of the orthographic stream. However, activity did not extend to the anterior-most region (white arrows) which was connected to the LANG network (black borders). This same region was active when listening to speech (**Fig. 3**). **Supp. Fig. S28** shows the non-binarized heatmaps.

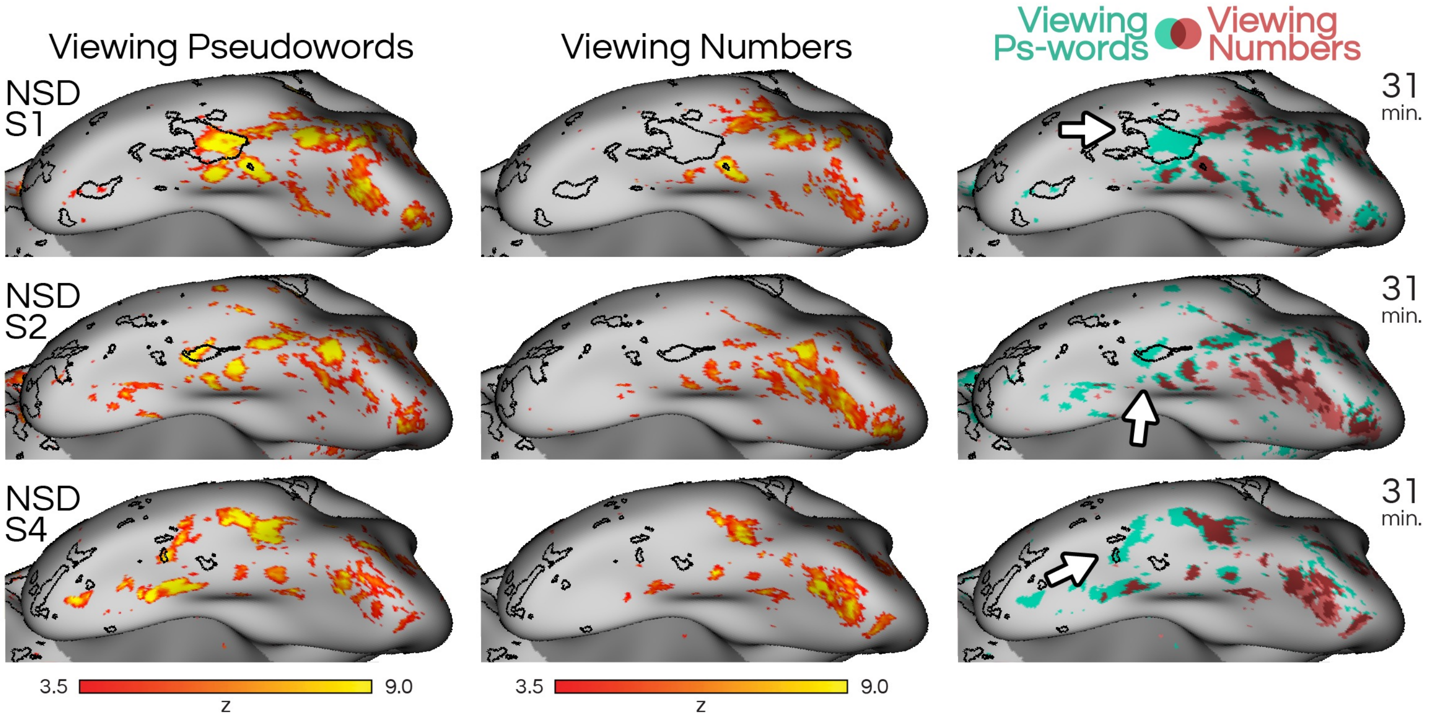


**Supp. Fig. S33: In NSD subjects, number strings also activate most of the orthographic stream apart from the most anterior region that is connected to the LANG network.** *Left column:* Heatmap shows activity when viewing pseudowords compared to other image categories (bodies, faces, places, objects). *Middle column:* Heatmap shows activity when viewing number strings compared to the same categories (bodies, faces, places, objects). *Right column:* Binarized pseudoword (teal) and number (red) activity overlaid on the same surface to reveal that number strings activated the majority of the more posterior orthographic stream regions (teal) but not the anterior-most region which was connected to the LANG network (black borders).

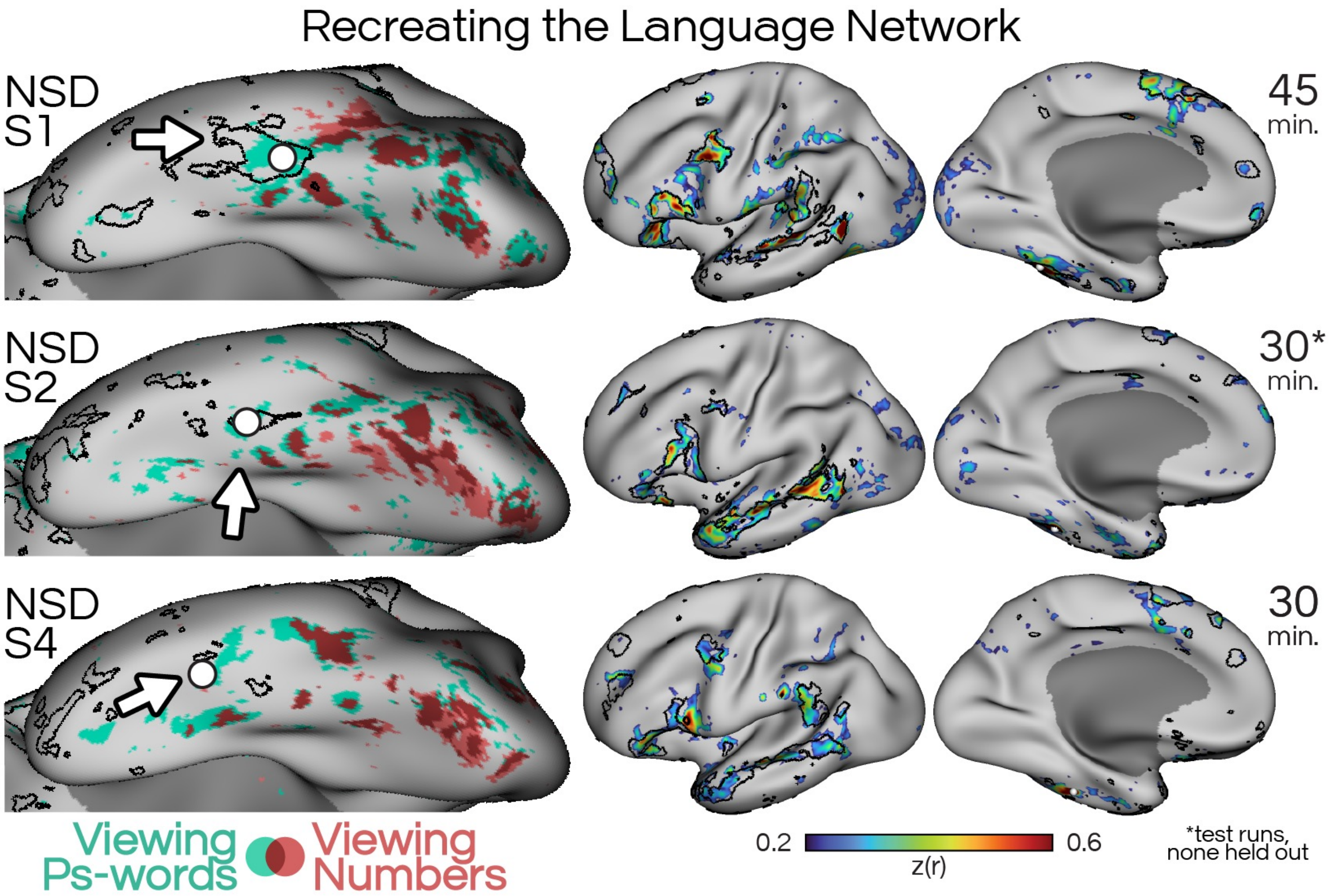


**Supp. Fig. S34: The LANG network could be replicated from inferior temporal seeds in the NSD subjects.** *Left:* Black boundaries indicate the basal LANG network region defined by data-driven clustering. For each participant, a seed (white circle) was chosen in mid-ITC targeting a region within the basal LANG region that showed increased activity for pseudoword (teal) but not for numbers (red). *Right:* The resulting seed correlation pattern (Turbo colormap), calculated using held-out resting-state data for NSD-S1 and NSD-S4, replicated the organization of the LANG network. Total time of held-out rest runs for each participant is on the far right.
